# Differing impacts of global and regional responses on SARS-CoV-2 transmission cluster dynamics

**DOI:** 10.1101/2020.11.06.370999

**Authors:** Brittany Rife Magalis, Andrea Ramirez-Mata, Anna Zhukova, Carla Mavian, Simone Marini, Frederic Lemoine, Mattia Prosperi, Olivier Gascuel, Marco Salemi

## Abstract

Although the global response to COVID-19 has not been entirely unified, the opportunity arises to assess the impact of regional public health interventions and to classify strategies according to their outcome. Analysis of genetic sequence data gathered over the course of the pandemic allows us to link the dynamics associated with networks of connected individuals with specific interventions. In this study, clusters of transmission were inferred from a phylogenetic tree representing the relationships of patient sequences sampled from December 30, 2019 to April 17, 2020. Metadata comprising sampling time and location were used to define the global behavior of transmission over this earlier sampling period, but also the involvement of individual regions in transmission cluster dynamics. Results demonstrate a positive impact of international travel restrictions and nationwide lockdowns on global cluster dynamics. However, residual, localized clusters displayed a wide range of estimated initial secondary infection rates, for which uniform public health interventions are unlikely to have sustainable effects. Our findings highlight the presence of so-called “super-spreaders”, with the propensity to infect a larger-than-average number of people, in countries, such as the USA, for which additional mitigation efforts targeting events surrounding this type of spread are urgently needed to curb further dissemination of SARS-CoV-2.

---

Since its emergence from Wuhan, Hubei Province, China, in 2019 and its established human-to-human transmission, government and local bodies have been working to control the spread of coronavirus disease (COVID-19). Severe acute respiratory syndrome coronavirus 2 (SARS-CoV-2), the pathogen responsible for this disease, is a single-stranded RNA virus that likely emerged through recombination events within animal reservoirs infected by different strains (*1, 2*). Since December 2019, its rapid spread throughout the world has already resulted in more than ten million cases and hundreds of thousands deaths, with no vaccine or specialized medication currently available.

Prior to COVID-19, the most recent global pandemic that presented a serious public health emergency was caused by influenza A H1N1 strain in 2009. H1N1’s threat exposed vulnerable public health capacities at the global, national and local levels, which we are facing once again, such as limitations of scientific knowledge, dilemmas in decision making, and communication among experts, policymakers and the public (*3*). The failure of preventive measures can lead to outbreaks crossing borders and exceeding national capacities (*4*). The likelihood of worldwide spread for pathogens characterized by human-to-human transmission, such as H1N1 and SARS-CoV-2, is extremely high with today’s globalized economy and ease of international travel and requires additional, concerted efforts of governments and public health institutions to prevent or contain outbreaks. As individual nations have responded in various ways, and at varying times throughout the pandemic, understanding the impact of these efforts on the global and regional transmission dynamics of the virus are imperative in defining a strategy to assess the current situation and prevent similar scenarios in the future.

Genomic data sampled from viral epidemics offer a unique opportunity to evaluate not only the evolution of the virus over time but also the changing viral population dynamics imprinted in the evolutionary history. These population dynamics, though often reflective of the neutral processes of evolution (*5*), can at times be traced to significant ecological and epidemiological events (*6*). Sample collection dates are critical in these connections, as when combined with the assumption of relatively stable mutation rates over time, allow for genetic differences to be rescaled as differences in time. The timing of population changes, inferred from genetic changes, can then be compared with external, contextual data (e.g., (*7*)) to investigate potential links between public health interventions (or other relevant events) and viral population growth and spread. Global analyses of viral spread using genomic data can reveal important information regarding the emergence of a novel virus, such as the phylogenetic analysis of H1N1 (*8*), which informed the community of the adaptive process of H1N1 from its original host (swine) to humans and its subsequent challenges in escape from the human immune system. Since then, large projects aimed at facilitating and optimizing the sharing of data and results for real-time projections of viral spread have helped in tracking SARS-CoV-2 global dissemination (e.g., (*9, 10*)). Yet, given that viruses evolve at a relatively rapid rate, considering separate isolated geographic areas as separate epidemics may also be warranted when attempting to understand how regional efforts drive viral evolutionary and population dynamic patterns. A virus from country X that seeds infection in country Y (founder event) becomes, over time, genetically distinct from the strains circulating in the country of origin, even in the absence of selection, due to genetic drift. On the other hand, in the era of globalization, analyses on regional epidemics limited to regional data can miss out on critical travel-mediated variables, such as separate, independent introductions of the virus (*11*).

Viral genetic data are not only useful in reconstructing the evolutionary and demographic history of a viral epidemic but also in identifying putative direct transmission events when *a priori* knowledge, or estimates, of the maximum genetic distance that separates linked individuals exists (e.g., (*12*)). Transmission clusters are of interest to public health, as they represent groups of individuals related by a common denominator, or risk factor, such as locality, social network structure, or other behavior (e.g., (*13*)); the connectedness of these individuals is reflected in closely related genetic sequences. With advanced efforts in SARS-CoV-2 genomic sequencing, phylogenetic tools can help to identify and characterize clusters of transmission, as well as regions involved in those clusters. Using such transmission cluster data, we propose that patterns in cluster formation, growth, and connectedness among otherwise separate geographical regions offer insight not only into global components of the public health response, but can also help in identifying clusters and regions for which a more specific strategy to control local spread is required.

## Results

### Cluster size and composition

Using a large genomic dataset collected from the Global Initiative on Sharing All Influenza Data (GISAID) aggregation of SARS-CoV-2 data, we hypothesized the existence of a relationship between international travel restrictions and overall transmission cluster dynamics, as well as sufficient variability among clusters in both space and time that would reveal the impact of varied local public health interventions. Based on the CDC definition of molecular transmission clusters for HIV, well-supported clades of viral sequences (one sequence per patient) comprised of at least 5 individuals with similar genetic distances were considered in this study to qualify as putative transmission clusters (also used in (*14*) for SARS-CoV-2). The criterion for similar genetic distances within these clades was a median patristic distance (branch length separating sequences within the tree) of <0.009% (see Supplementary Materials) in a maximum likelihood (ML) phylogenetic tree inferred from 11,069 SARS-CoV-2 genomic sequences. This distance, representing the median genetic difference between two sampled individuals is less than the median difference observed within a single individual (0.014%) in the study by Shen et al. (2020) (*15*) on patients admitted with COVID-19 pneumonia. By only considering individuals that share a genetic distance less than would be expected during the evolution of the virus within a single host, we have greater confidence in infections separated by a short amount of time and thus an epidemiological linkage or connection; increasing genetic distance leads to increased uncertainty as to the relationship of individuals within a cluster and a greater likelihood of the inclusion of multiple risk factors. The majority of clusters identified using this method included *<* 25 individuals, with one large outlier cluster of 185 individuals (184 from the USA and 1 from Denmark) (Figure 1A). In terms of country representation within transmission clusters, the majority of the identified clusters included viral sequences isolated from patients in 1 *−* 5 countries, with the majority country in clusters comprised of only two countries representing 50-99% of the sequences (Figure 1B). Clusters including 6-10 countries were more evenly distributed, while the majority country represented 70% of the sequences in the only cluster with strains from 11 countries. The results collectively indicate a significant role for travel in connecting several countries through putative direct transmission events, rather than isolated epidemics seeded by single introductions, consistent with previous studies (*16, 17*).

**Figure 1:**
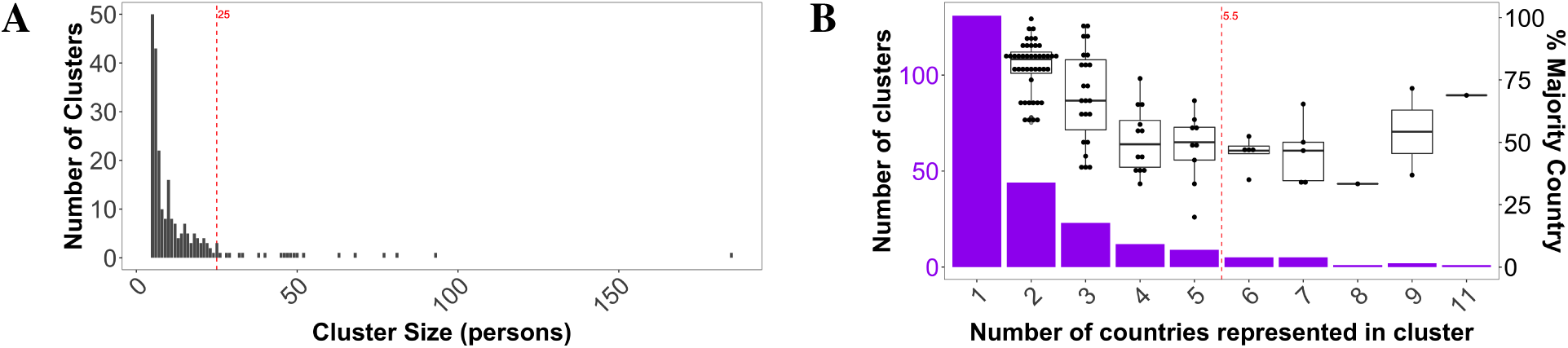
Transmission cluster distributions against cluster size (A) and number of countries represented in each cluster (B). Percentage of sequences comprising the majority represented country has also been plotted (box plots) in (B). Red, dashed lines indicate median values.

As with conventional epidemiological analyses, sampling bias can impact results and interpretation and has been inherent to SARS-CoV-2 sample collection throughout the pandemic (*18*). In our analysis, the number of clusters involving each of the countries with available sequence data as of April 24, 2020, were distributed similarly to the number of available genomes indicating, unsurprisingly, that the number of clusters detected for a particular country is limited by the number of available samples from that locale (Figure 2A). Hence, no direct comparison could be made regarding the number of clusters involving different countries. This was not the case, however, for the average size (number of persons) of clusters associated with each country, since large clusters were not always detected in countries with more samples available, such as the United Kingdom (UK) and United States of America (USA) (Figure 2B). In other words, despite potential lack of information from countries with reduced datasets, a global perspective on transmission cluster patterns related to cluster size might nevertheless be investigated. It is important to keep in mind that low sequence presence is not necessarily an indicator of low infection rates, as the fraction of total infected individuals that were sampled may, in fact, be high for these countries, resulting in a different type of regional sampling bias. For example, although as of April 24, over 3,000 sequences were submitted for the UK, this comprised *<* 25% of total infections, whereas Hong Kong reportedly submitted at least one genome for every confirmed case (Figure 2C). Given that individuals linked through transmission in a small amount of time share a small genetic difference, on which phylogenetic cluster inference relies, missed sampling can prevent the inclusion of individuals within a cluster. Hence, we anticipated that sampled individuals from countries with sequencing more representative of the infected population (i.e., genome per confirmed case value closer to 1) would be more likely to cluster, resulting in a larger fraction of clustered individuals. However, there was no clear relationship (linear regression *R*^2^ *<* 0.0084) between the percentage of individuals within a country that are included in clusters and the number of retrieved genomes per confirmed cases (Figure 2C). This finding indicates that country-specific contributions to clustering were not biased toward countries with larger fractions of sampled individuals from the infected population. Therefore, sub-sampling as an effort to mitigate the effects of sampling bias at the country level, though often performed for phylogenetic analyses of viral geographical spread (*19*), was not deemed necessary for our study. On the other hand, it is important to notice that due to the lack of information on sampling strategies used to gather available sequence data, we could not rule out possible effects of selection bias (i.e., preferential sampling). For example, in certain countries for which representative sequencing was low but clustering rate was high, we cannot exclude that sampling efforts were focused on presumed contact networks in order to locate and quarantine infections.

**Figure 2:**
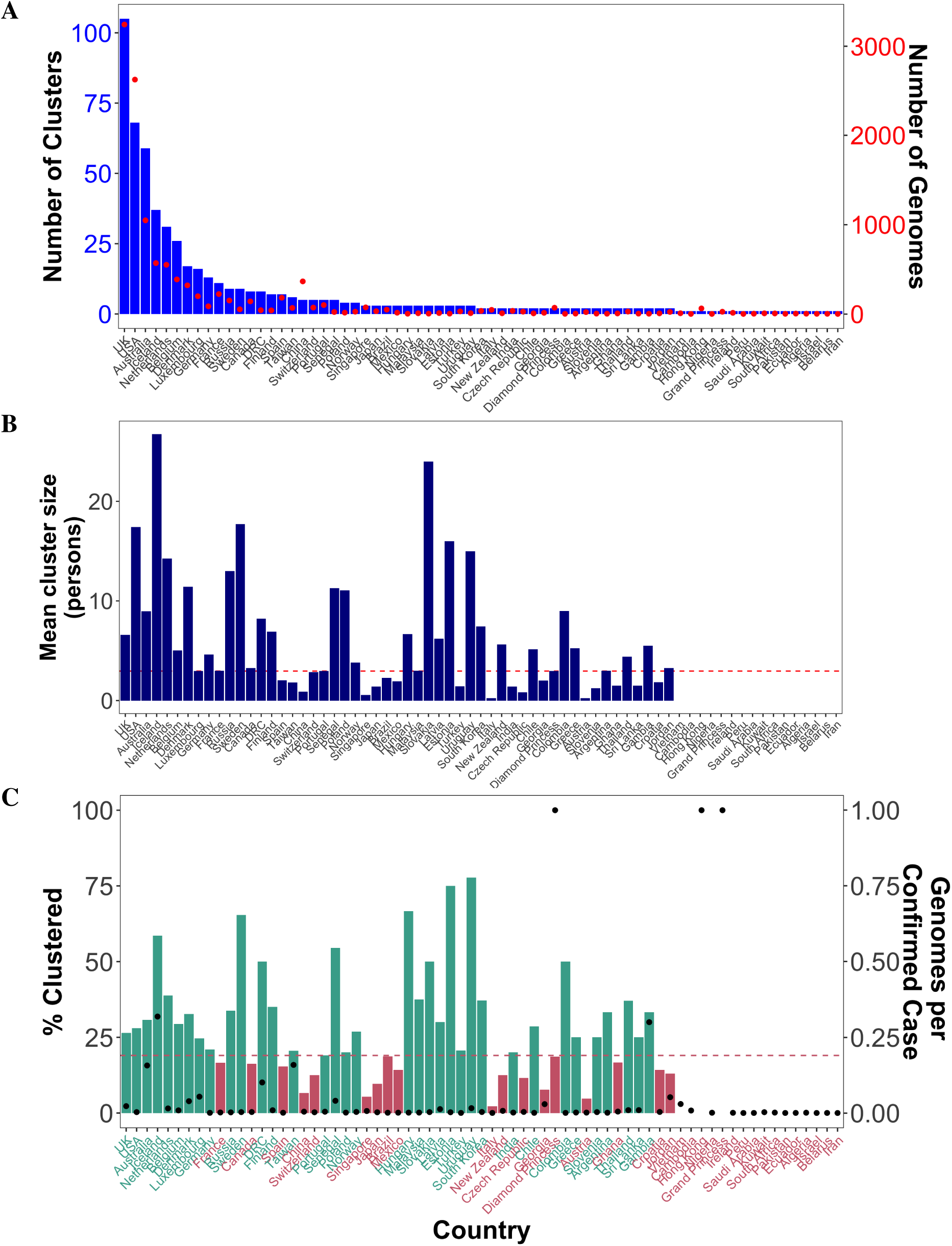
Transmission cluster characteristics for each country involved. Number of clusters (A), mean cluster size (B), and percentage of individuals that clusters (C) are plotted for each country, as well as the number of available genomes (A) and genomes per confirmed case reports (C) for comparison.

### Cluster origin and epidemiology

In order to derive epidemiological information for each detected cluster, branch lengths within the tree were scaled in time by enforcing a molecular clock, which assumes the accumulation of substitutions has occurred at a constant rate over time. The time to the most recent common ancestor (TMRCA) of all sequence data was estimated to be December 6 [25 Nov - 10 Dec] 2019 (Figure S1, consistent with previous demographic model-based estimate by Andersen et al. (2020) (*20*). The result was a good indication that date estimates for remaining internal nodes within the tree were also reliable.

For the majority of the transmission clusters detected in the tree, TMRCAs (i.e., a cluster’s temporal origin) dated back prior to Feb 20, 2020, with a peak observed between the first week of February until the first week of March (Figure 3A). These results were robust to genetic distance thresholds of 0.006% and 0.013% (Figure S2), the latter value representing the 2.5th percentile of tree-wide distances and maximum diversity observed in Shen et al. (2020) (*15*), as described above. Following this time, a sharp decline in number of newly formed clusters was evident. Furthermore, despite differing patterns in the number and size of clusters across individual countries, the global peak in clusters size coincided with the peak in number of new clusters, around the end of February, as did the number of countries represented in each cluster (Figure 3A). The overlapping peaks in cluster number and size, as well as number of countries/cluster in time, suggest a common underlying factor responsible for the decrease in the rate of cluster formation, growth, and connectivity at the global level. Thus, it appeared reasonable to hypothesize that efforts to reduce international travel at the beginning of the epidemic played a role.

**Figure 3:**
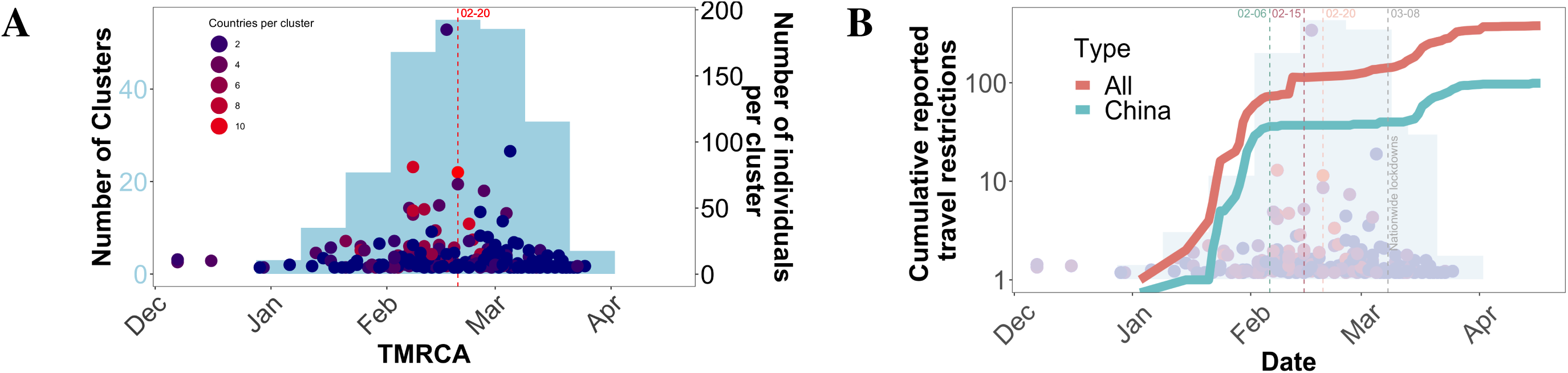
Timing of transmission cluster characteristics. (A) Number of clusters (light blue) sharing a similar temporal origin, as inferred from the time to the most recent common ancestor(TMRCA) of corresponding cluster sequences. Dots correspond to the size (number of individuals) within the corresponding clusters at individual time points. Dots are colored according to the number of countries represented in each cluster. (B) Number of cumulative reported international (red) and China (teal) travel restrictions over time superimposed onto (A), ending in the first reported case of local travel restrictions (grey) for visual clarity. Red, dashed lines indicated median values.

Data on reported travel restrictions (Tables S1 and S2) were obtained from various sources (see Supplementary Materials), and the cumulative number of reports involving all international travel, as well as those specifically involving China, were plotted over time to provide context into travel-related events that would potentially result in the global cluster behavior described above. The early, elevated rate of accumulation of travel bans, largely comprised of restriction on immigration from China began to slow on February 15 (slowed accumulation for China-specific bans on the 6th) (Figure 3B). Accumulation in the number of restrictions remained low until the onset of nationwide lockdowns (first reported March 08), at which time the steep decline in overall cluster TMRCA, size, and geographical range began. These results strongly suggest that reduced international travel slowed the formation, growth, and connectivity of transmission clusters, whereas more local interventions were necessary, and more effective, in preventing local virus spread.

The median lifespan, or duration, of a transmission cluster was estimated to be approximately 2 weeks, with the largest cluster extending for over six weeks (Figure 4A). Such a short duration time (on the order of the longer end of the incubation period (*21*)), combined with average cluster sizes of up to 25 sampled individuals, is indicative of SARS-CoV-2 rapid transmission (*22*). The expected number of secondary cases directly generated by a primary case in the population, otherwise known as the basic reproductive number (*R*_0_), was calculated as a function of early changes in the estimated viral effective population size (*23*) and a normally distributed infectious period of approximately 2*−*8 days). *R*_0_ for the entire pandemic was estimated at 5.65 [95% credible interval (CI): 4.37*−*6.68], consistent with previously reported *R*_0_s by Shen et al. (2020) (*15*) and Tang et al. (2020) (*24*), as well as other published (but not peer-reviewed) estimates reviewed in Liu et al. (2020) (*22*). However, in contrast to previous epidemiological studies, we utilized phylogenetic methods at an increased resolution to estimate the transmission potential for individual clusters, which we have already shown can vary in size and composition over time. In the majority of clusters, *R*_0_ ranged from *<* 1*to*2.32, though *R*_0_ values of up to 12 were reported (Figure 4B), indicative of clusters formed through “super-spreading” events (SSEs), or cases of larger-than-average transmissibility (*25*). While it is important to note that accuracy of *R*_0_ estimation is reduced for outbreaks characterized by true *R*_0_ *≥* 5 (*23*), clusters representing increased secondary infection rates as compared to the majority population are of importance in the control of infection. As *R*_0_ *>* 1 is indicative of sustainable transmission and vice versa, we investigated the timing and duration of both low(*<* 1)- and high(*>* 1)- *R*_0_ clusters. High-*R*_0_ clusters were more frequently observed between February and early March (Figure 4C), consistent with the temporal peak in global cluster size. There was no relationship between *R*_0_ and duration (*R*^2^ *<* 3.69*E −* 06), suggesting clusters with *R*_0_ *<* 1 were still sustained for various lengths of time. Although seemingly counter-intuitive, this finding can be explained by dynamic transmission patterns, such as a change in the contact network, or even a later introduction of unsampled super-spreaders. As the *R*_0_ calculation is derived from early estimates of the viral effective population size, this value does not depict the full transmission potential of the group of individuals within the cluster.

**Figure 4:**
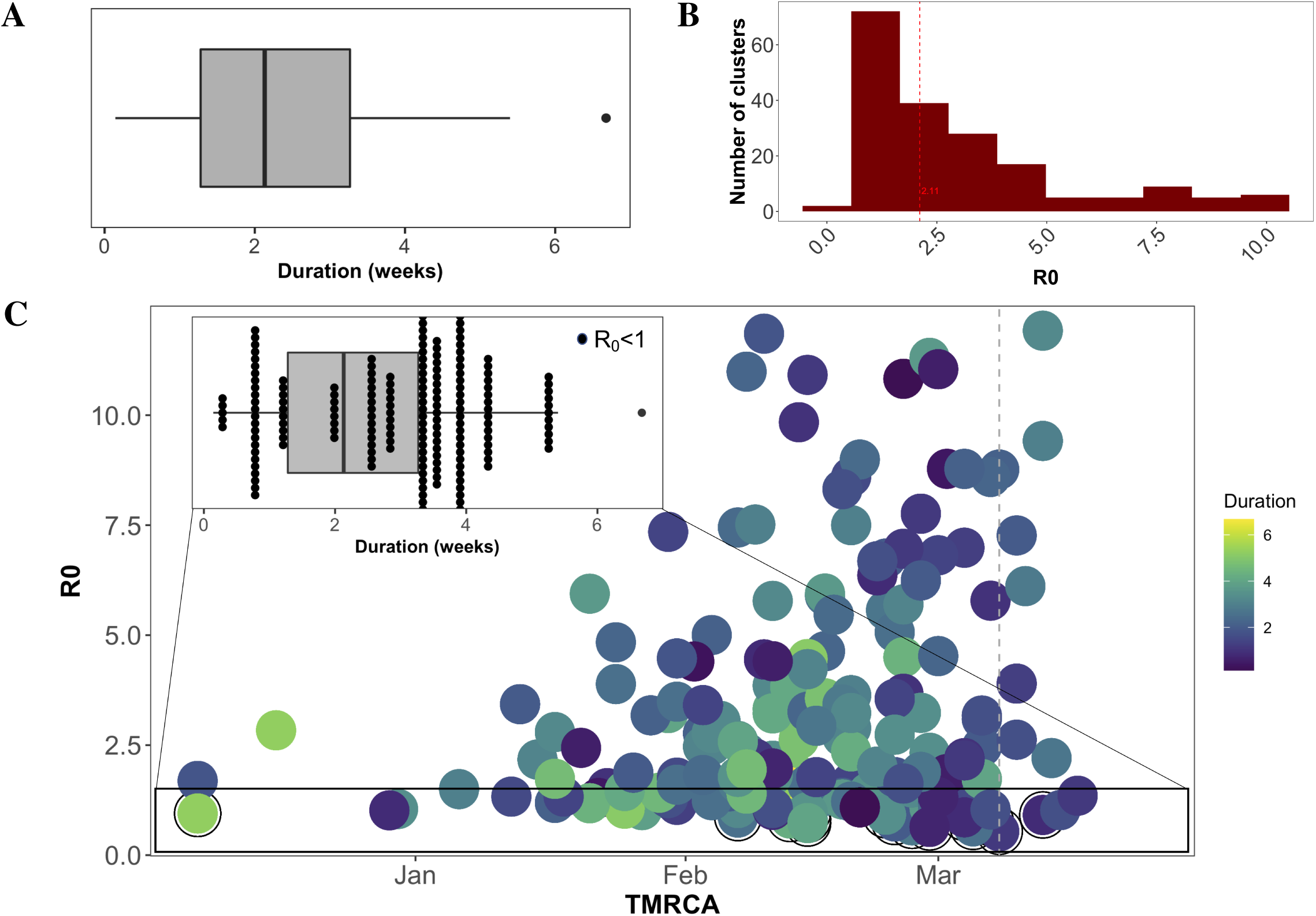
Estimated transmission cluster duration and *R*_0_. (A) Duration was defined as the time from TMRCA to most recent sampling date for each cluster. (B) *R*_0_ was derived from estimates of changes in effective population size, as described in (*23*). (C) Relationship of *R*_0_ with time of cluster formation and cluster duration. Inset focuses on the distribution of *R*_0_ *<* 1 clusters within the overall distribution of cluster duration.

Our results point to a drastic reduction in cluster formation, growth, and connectivity following the first week of March. Yet, clusters with *R*_0_ *>* 2 (earlier estimates of SARS-CoV-2 *R*_0_) were observed during this time (Figure 4C). We next sought to investigate which countries were involved in the few, albeit seemingly rapidly spreading, clusters that formed following the onset of lockdown measures (e.g., social distancing). *R*_0_ values across clusters for each country were averaged after being scaled based on the percentage of sequences belonging to that country, resulting in a weighted mean *R*_0_ (Figure 5A). Belgium, Luxembourg, and the UK and USA (alphabetical order only) were estimated to have a weighted mean *R*_0_ *>* 2 in March, whereas Australia, Iceland, and the Netherlands were approximately 1 or less (*R*_0_ *<*1.1, 1.2, and 1, respectively) (Figure 5A). In line with global reduction in connectivity, clusters formed after March 08 consisted of 2 countries or less Figure 5B. The USA formed two clusters comprised of only US individuals, both with *R*_0_ *>* 2, one of which was estimated as the highest *R*_0_ value (*>* 10, CI: 7.57-11.94) of all clusters at this time. The three clusters following in ranking comprised either Belgium alone (2) or Belgium and neighboring Luxembourg (2), with *R*0 *>* 4. All of these clusters exhibited evidence of sustained transmission, with a duration of greater than 2 weeks, though it is important to note that the addition of more up-to-date sequences may extend duration times for clusters initiated during this time period. The UK formed three separate clusters - one isolated, one with primarily UK individuals, and one with primarily Australian individuals. The two clusters comprised of majority UK sequences were both estimated to have *R*0 *>* 2, whereas the cluster with primarily Australian sequences was characterized as *R*0 *<* 1. Similarly, the second Australian cluster (Australian sequences only) was also characterized as having a relatively low *R*_0_ of *<* 1.3 (CI:1.20-1.32). Results suggest that the virus was already spreading rapidly in the USA, Belgium and Luxembourg, and the UK at the time of implementation of regional mobility restriction efforts and that, despite efforts to restrict international travel, the UK and Australia maintained travel sufficient to sustain inter-regional transmission.

**Figure 5:**
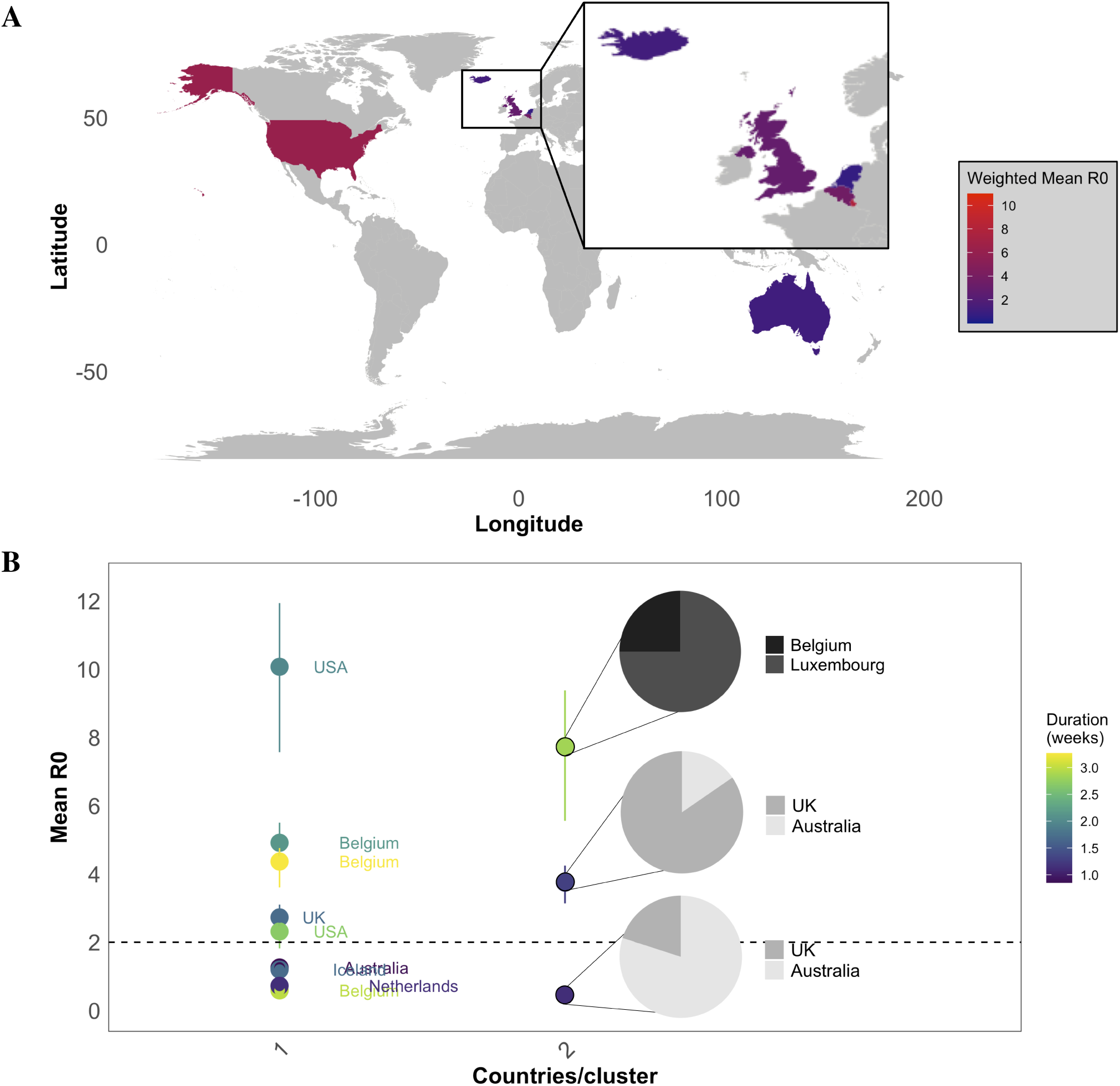
Summarized transmission patterns for countries involved in clusters formed after March 08, 2020. (A) Mean *R*_0_ for each country was calculated by weighting each cluster-specific *R*_0_ according to the fraction of sequences belonging to the country.(B) Estimated mean *R*_0_ and 95% confidence intervals for each cluster (dot). For clusters with more than one country, the fraction of sequences belonging to each country is depicted in the corresponding pie chart. Clusters are also colored according to duration. Dashed line depicts *R*_0_ = 2

Recent evidence implicating a mutation in residue 614 of the spike protein of SARS-CoV-2 in increased infectivity (*26*) and higher mortality (*27*) offered a possible explanation for increased transmission potential of certain clusters identified in this study. However, despite the clear advantages at the cellular level of glycine in place of aspartic acid at this site, there was no relationship between prevalence of glycine (% of individual sequences) within individual transmission clusters and *R*_0_ (linear regression *R*^2^ *<* 0.00011) or cluster size (*R*^2^ *<* 0.0083) (Figure S3), though generation time was not explored.

## Discussion

Unlike conventional epidemiological surveillance data, viral genetic data can be used to link infected individuals involved in direct transmission events even when the contact structure is unknown. Identification of these putative transmission clusters, coupled with phylogenetic inference of cluster dynamics and relevant epidemiological parameters, has provided evidence of a global pattern in cluster formation, growth, connectivity, as well as transmission potential over time. The slow rise and subsequent rapid fall in the number and size of forming clusters over the period of December 30 to April 20, 2020, may have been the results of a change in global transmission patterns and/or a rate of sampling that did not adequately capture, as recently reported (*28*), the increased rate in the number of infected individuals. According to the latter scenario, missed sampling of individuals involved in a transmission cluster can prevent the detection of links via genetic data, or even conventional surveillance data. However, the similar temporal pattern observed for the number of countries connected by individual clusters also suggests a relationship between cluster formation, growth, and connectivity and that an outside force was responsible for changing global transmission patterns, rather than problematic sampling over time.

When placed in the context of the timing and accumulation of public health interventions, the data provide evidence supporting the benefits of both global and regional response efforts. Given an incubation period extending to up to 18 days within an individual (*29*), it is plausible that specific restriction on international travel involving China, peaking on February 06, could have resulted in the slowed rate of cluster activity beginning the first week of February. It is also possible that the rapid increase in overall international travel restrictions, peaking around February 15, was partially, if not equally, responsible. Whereas additional modeling using recorded travel data would be highly beneficial in teasing apart the effects of international and China-specific travel restrictions on transmission characteristics, the drastic halt in cluster activity beginning the first week of March directly coincides with the date of onset of nationwide lockdowns. This particular finding speaks to the effectiveness of additional non-pharmaceutical measures taken at the regional level to curb the spread of infection, consistent with previous reports (*30, 31*).

Whereas a global reduction in cluster activity in response to efforts to reduce mobility might be expected given the known role of travel in pathogen spread (*32*), we anticipated cluster- and region-specific variation in transmission characteristics, consistent with known difficulties in controlling regional epidemics. Using viral sequence data and the phylogenetic relationships among sampled individuals, particularly for an exponentially growing epidemic of limited data availability (*23*), we can model relevant epidemiological parameters of interest used in determining the transmission potential for clusters involving individual countries. It is important to keep in mind that the results of any phylogenetic study that are dependent on a single tree assume that that a tree best describes the underlying phylogenetic relationships. While it is often best to summarize results across a sample of similarly plausible trees using Bayesian methods (*33, 34*), Bayesian tree reconstruction methods are highly parametric and have difficulty converging on a reliable distribution of trees and related evolutionary parameters for datasets as large as that of SARS-CoV-2. For this reason, we only focused this study on portions of the maximum likelihood tree that were considered to be well supported. Similarly, invaluable methods exist for the detection and epidemiological characterization of transmission clusters within the Bayesian framework, such as the multi-state birth death model (bdmm) (*35*); however, even the bdmm is limited to less than 1000 sequences (unpublished work by Scire et al. ((*36*)). While sub-sampling strategies used to reduce dataset size and potential sampling biases are widely appreciated in the field of phylogenetic epidemiology (e.g, (*19*)), they inherently result in loss of smaller clusters, which play an important role in assessing the effect of social distancing interventions. As our study is based on hypotheses regarding not only cluster characteristics but also cluster size distribution, a skew towards detection of larger clusters as a result of this loss of information was undesirable. *Post hoc* transmission characterization of individual clusters using a Bayesian framework for population dynamics estimates that has demonstrated accuracy for low-signal sequence data (i.e., small clusters) (*23*) was thus ideal for an in-depth investigation of transmission cluster dynamics at the global and regional scales. Using this method, four countries (the USA, Belgium and neighboring Luxembourg, and the UK) were identified as harboring isolated clusters with elevated transmission potential following the global initiation of nationwide lockdowns, potentially fostered by super-spreaders. Federal policies regarding lockdowns were put into place for Belgium, Luxembourg, and the UK beginning mid-March, which were likely necessary in absolving highly active clusters such as those observed in this study; though federal guidelines were issued at a similar time in the USA (March 16th), mandatory US policy regarding lockdowns was not put into motion, allowing individual states to adopt their own policies at different times. Depending on the location of transmission clusters within the USA, delayed lockdowns could have resulted in continued rapid transmission of the virus. It is also important to keep in mind that at the time of this study, a dramatic rebound was being observed in the number of detected daily cases in the USA, whereas the UK, Belgium and Luxembourg were demonstrating a consistent decline with occasional peaks (*9*). Moreover, the epidemic at the time was exponentially spreading in Brazil and India, as well as steadily in Russia - all countries that were not captured by our analysis based on sequence data up to the last week in April, 2020. In this context, our results emphasize the importance of additional country-specific transmission cluster analysis for data collected more recently than April 20.

It would be reasonable to suspect that the identification of the USA, Belgium, Luxembourg, and the UK as problematic countries was aided by the availability of sequence data from these locations (i.e., attributed to a sufficient number of individuals and genetic diversity to classify and characterize corresponding clusters), which at first points to a potential problem with sampling bias. Although these countries were indeed at the high end of the spectrum in terms of available sequence data, our analyses relied on averaged values across clusters, which we show is unrelated to genome availability, unlike number of clusters per country. These countries exhibited a wide range of clustering frequency (20-60% of individuals) with no relation to sampling representation (fraction of the infected population sampled), which varied from 1-30%, indicating with a high degree of confidence, that they were not identified as a result of sampling bias. Moreover, countries (Iceland, Australia, and the Netherlands) identified as having low transmission potential after March 08 (*R*_0_ less than or approximately 1) had a comparable number of available genomes within the distribution, lending support to the conclusion. This is not to say that selection bias, a form of sampling bias, may not be associated with the results. For example, Iceland’s low *R*_0_ during this time may not be surprising, given that the country’s genetic powerhouse, deCODE, began screening high-risk (symptomatic) individuals and those returning, or in contact with an individual, from high-risk countries as early as January 31 (*37*). Therefore, while in our study, concerns for the sampling representation of the overall infected population is negligible, more sophisticated quantitative measures to assess the impact of sampling biases, specifically selection bias, will be necessary in future investigations.

In summary, we propose that phylogenetic identification and characterization of transmission clusters using the vast resources of viral genomic data currently available can provide both global and regional perspectives on viral spread. When collected early in the course of an epidemic, as was the the case for the SARS-CoV-2, this approach may help to pinpoint locations for which increased efforts at the level of local government might be necessary to mitigate growth on a pandemic scale. At the time of the submission of these results, relaxation of these efforts was on the rise, particularly in the USA. The detection of isolated transmission clusters with elevated transmission potential in March, despite the rapid decline in global patterns of cluster activity, points to an important role played by super-spreaders in the current pandemic that likely pose a threat to relaxation (*38*) unless measures are taken to quickly recognize and predict these events. A better understanding of the underlying risk factors associated with related super-spreading events, including host, environmental, and behavioral factors (*39*), is necessary for targeting efforts aimed in avoiding recurrent rebounds in epidemic waves. Given the increased efforts in testing and sampling in the USA, as well as other countries, transmission cluster dynamic inference can help to identify these events and underlying risk networks for more precise intervention strategies.

## Acknowledgements

We would also like to acknowledge the health workers and researchers who generated the data,without whom this work would not have been possible. Funding for this work was provided by the National Science Foundation (DEB 2028221) and National Institutes of Health (R21AI138815).

## Author contributions statement

M.S., M.P., and B.R.M. conceived of the analyses, S.M. and C.M. retrieved the data, A.Z. and used their expertise (with guidance from O.G.) in sequence data quality control and tree reconstruction to produce the trees, M.P. used his expertise in Phylopart to identify transmission clusters, B.R.M. analyzed the clusters and prepared the manuscript, A.R. monitored and gathered data regarding region-specific public health interventions, and all authors helped to craft the discussion and review the the manuscript. The authors report no competing interests at the time of manuscript preparation and submission. All data is available in the manuscript or the supplementary materials.

## Supplementary Materials

### Materials and Methods

#### Sequence data, metadata, and phylogenetic reconstruction

A total of 11,262 sequences and associated metadata (time and country of sampling) were downloaded from GISAID (GI-SAID) on April 25, 2020. Sequence IDs can be found in Table S3 The number of confirmed cases for each corresponding country was retrieved from European Centre for Disease Prevention and Control. Cruise ships Diamond Princess and Grand Princess were treated as separate geographical regions for all analyses. Data regarding travel restrictions were obtained from WorldAware, COVID-19 Travel Restrictions Database, and Trip.com on June 4th, 2020; data on border closures from The New York Times on June 1st, 2020; airline restrictions from Bloomberg and Business Insider on June 4th, 2020. Data specifically for the Diamond Princess was retrieved from Business Insider on June 5th, 2020. Data on nationwide lockdowns, health screening measures, closure of non-essential business, the use of masks and restrictions of mass gatherings were obtained from different online newspapers, including CNN, South China Morning Post, The New York Times, The Miami Herald, and Reuters on June 2nd.

Sequences were quality filtered and aligned, as described in (*40*), keeping one representative per identical sequence group and excluding sequences without precise dates (month and day), resulting in 11, 316 sequences (29, 726 nucleotides in length). We reconstructed a maximum likelihood tree with RAxML-NG (*41*) (GTR+I+G6 model, with all parameters optimized) starting from a distance tree (reconstructed using mimimum evolution in FASTME (*42*)), rooted it at the most recent common ancestor of the likely first-generation strains from (*1*), and put back the identical sequences (as zero-branch polytomies with the corresponding representative sequence tip). We then collapsed the branches without phylogenetic signal (i.e., of length *≤* 1*/*2 mutation per genome). Support for branching events within the tree were calculated using the the Shimodaira-Hasegawa approximate likelihood ratio test (*43*), performed in IQ-TREE v2 (*44*) on the final, fixed RAxML-NG tree topology.

Least squares dating in LSD2 (*45*) was used to date internal nodes of the tree (given sampling dates of taxa) and to identify, and remove, outlier sequences. Outlier sequences were defined as taxa whose mutation rate (estimated as the distance to the root divided by time since the first sequence) was larger than 3 standard deviations from the median. A strict molecular clock was assumed, resulting in an estimated mutation rate of 2.3[2.0*−*2.4] *·* 10^4^ mutations per site per year.

#### Transmission cluster identification

Clades comprised of *≥* 5 distinct sequences and a reliability of *≥* 90% were recognized as potential transmission clusters when the median pairwise patristic distance (i.e., branch length separating sequences) within the clade was below a pre-specified percentile threshold of the whole-tree patristic distance distribution. A range of percentile thresholds spanning 0.0005% *−* 25% of the whole-tree distance distribution was used to choose an optimal threshold point and to verify robustness of cluster composition. The minimum percentile threshold that maximized the number of clusters was chosen as the optimal threshold by performing multiple clustering runs on randomly sampled patristic distance distributions (1 million for each run) in Phylopart v2 (*46*). Clusters and corresponding information (including sequence IDs) can be found in Table S4.

Potential clusters with more than two time points were used for the calculation of the basic reproductive number (*R*_0_) from the viral effective population size (*N_e_*) estimated using the skygrowth (*23*) package in R (*47*). Viral effective population size (*N_e_*) estimates were allowed to vary weekly, and the default smoothing parameter (*tau*) of 0.1 was used. As described in Volz Didelot (2018) (*23*), the mean effective reproductive number (*R_e_*) was calculated as a function of the change in *Ne* and 2-8 day (mean=5.475, SE=1.825) infectious period (*psi*). *R*_0_ was defined as the first *R_e_* value in time. Clusters with outlying *R*_0_ values (*>* 3.4 standard deviations above or below the mean over all clusters) were discarded as unreliable.

---

**Figure S1:**
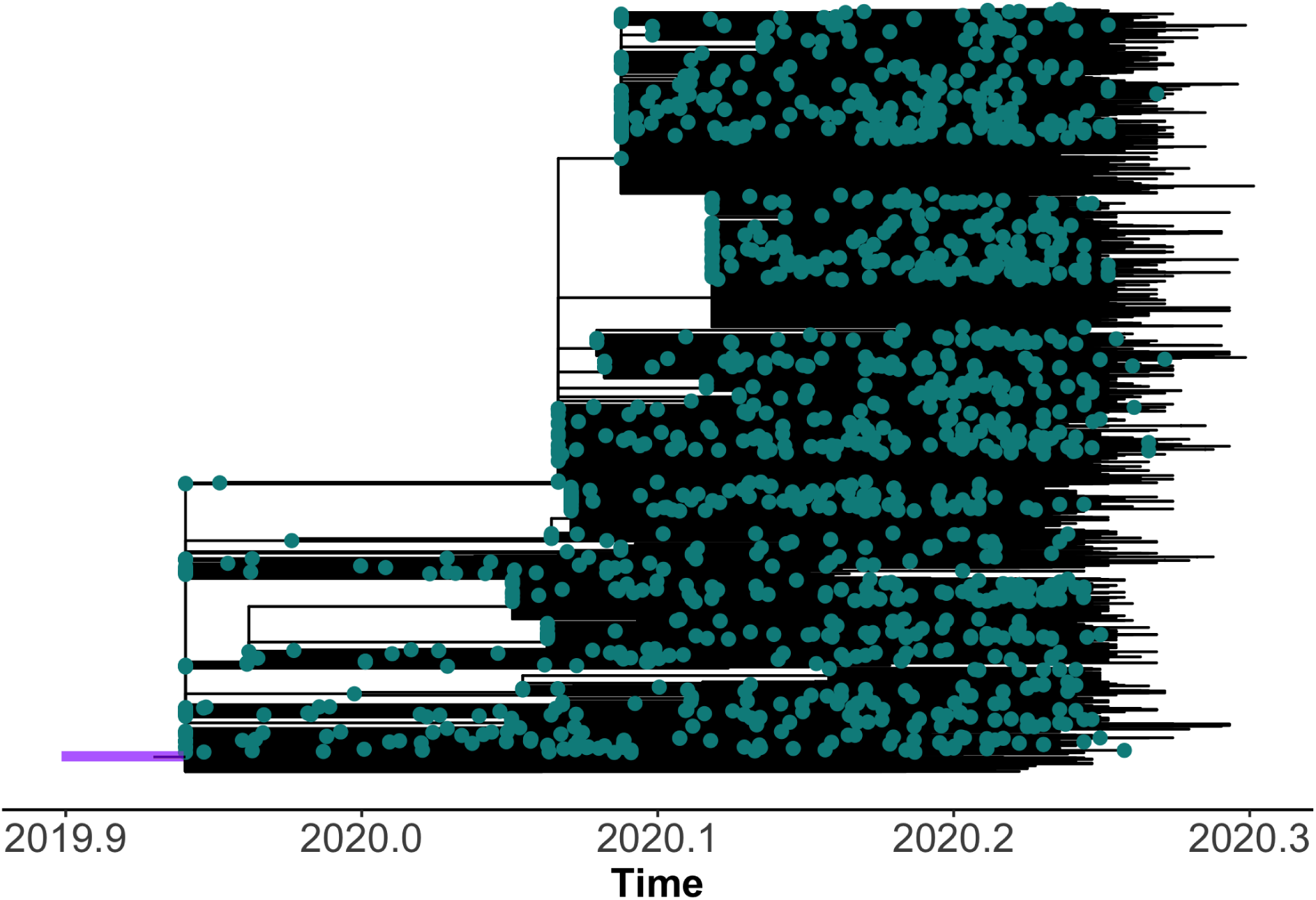
Maximum likelihood tree for 11,069 SARS-CoV-2 sequences scaled in time using least squares dating (*45*). Purple bar represents the 95% credible interval for estimate of the time to the most recent common ancestor for all sequences. Outlier sequences have been pruned and were excluded from downstream analyses (see Materials and Methods).

**Figure S2:**
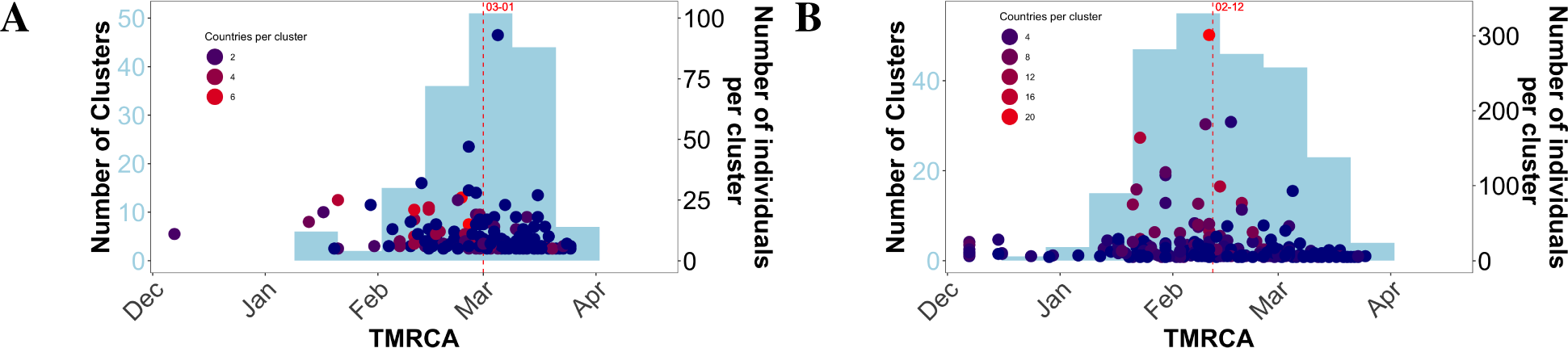
Timing of transmission cluster characteristics for additional distance thresholds defining transmission clusters. (A) Clusters with median patristic distances within 0.5% of the whole-tree patristic distance distribution. (B) Clusters with median patristic distances within 2.5% of the whole-tree patristic distance distribution. Bars represent the number of clusters (light blue) sharing a similar temporal origin, as inferred from the time to the most recent common ancestor (TMRCA) of corresponding cluster sequences. Dots correspond to the size (number of individuals) within the corresponding clusters at individual time points. Dots are colored according to the number of countries represented in each cluster. Red, dashed lines indicated median values.

**Figure S3:**
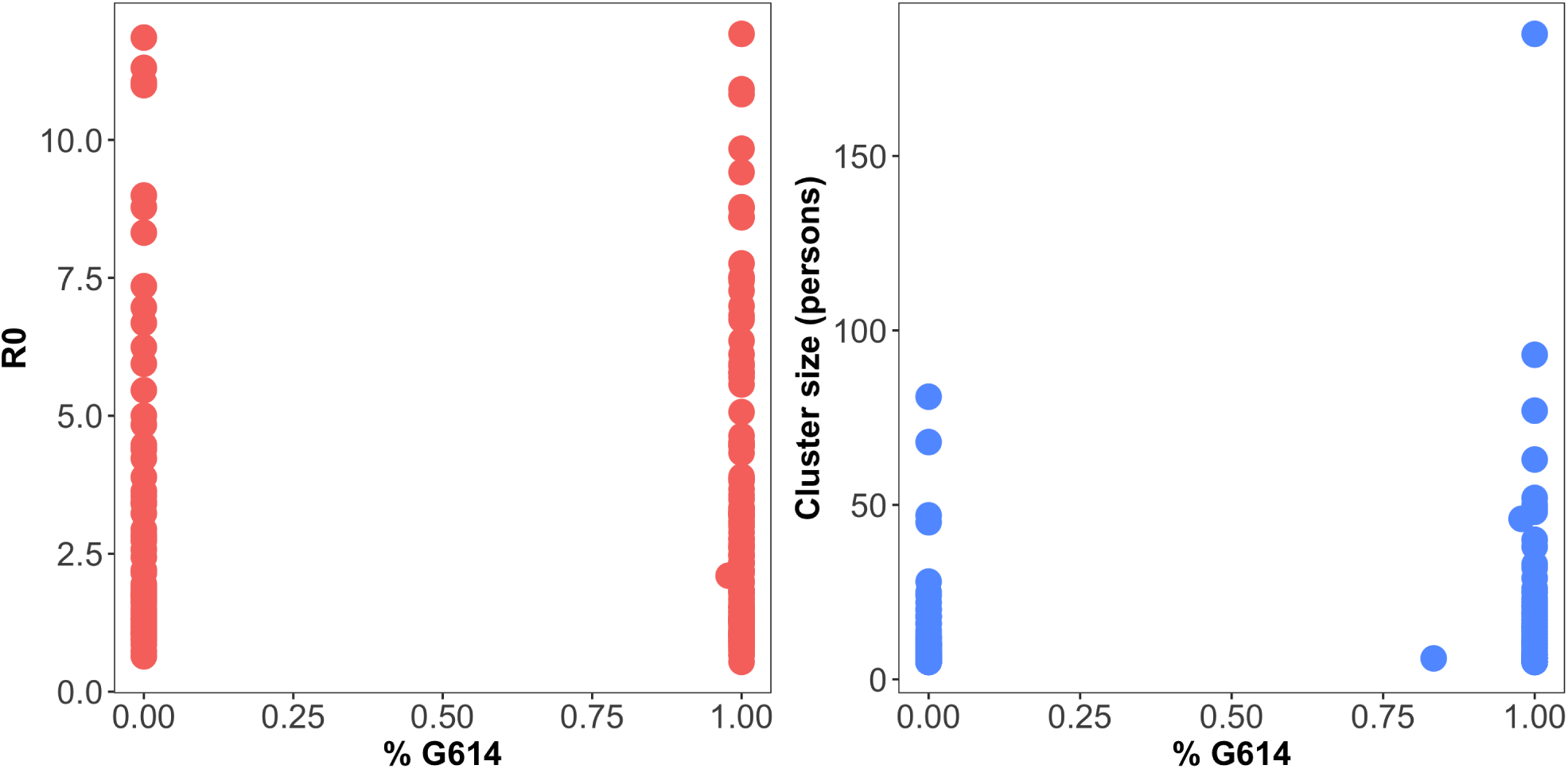
Relationship of *R*_0_ (A) and (B) cluster size with fraction of individuals with D614G mutation for each cluster. Each dot represents an individual cluster. Clusters in (A) represent a subset of clusters in (B) (see Materials and Methods).

**Table S1:**
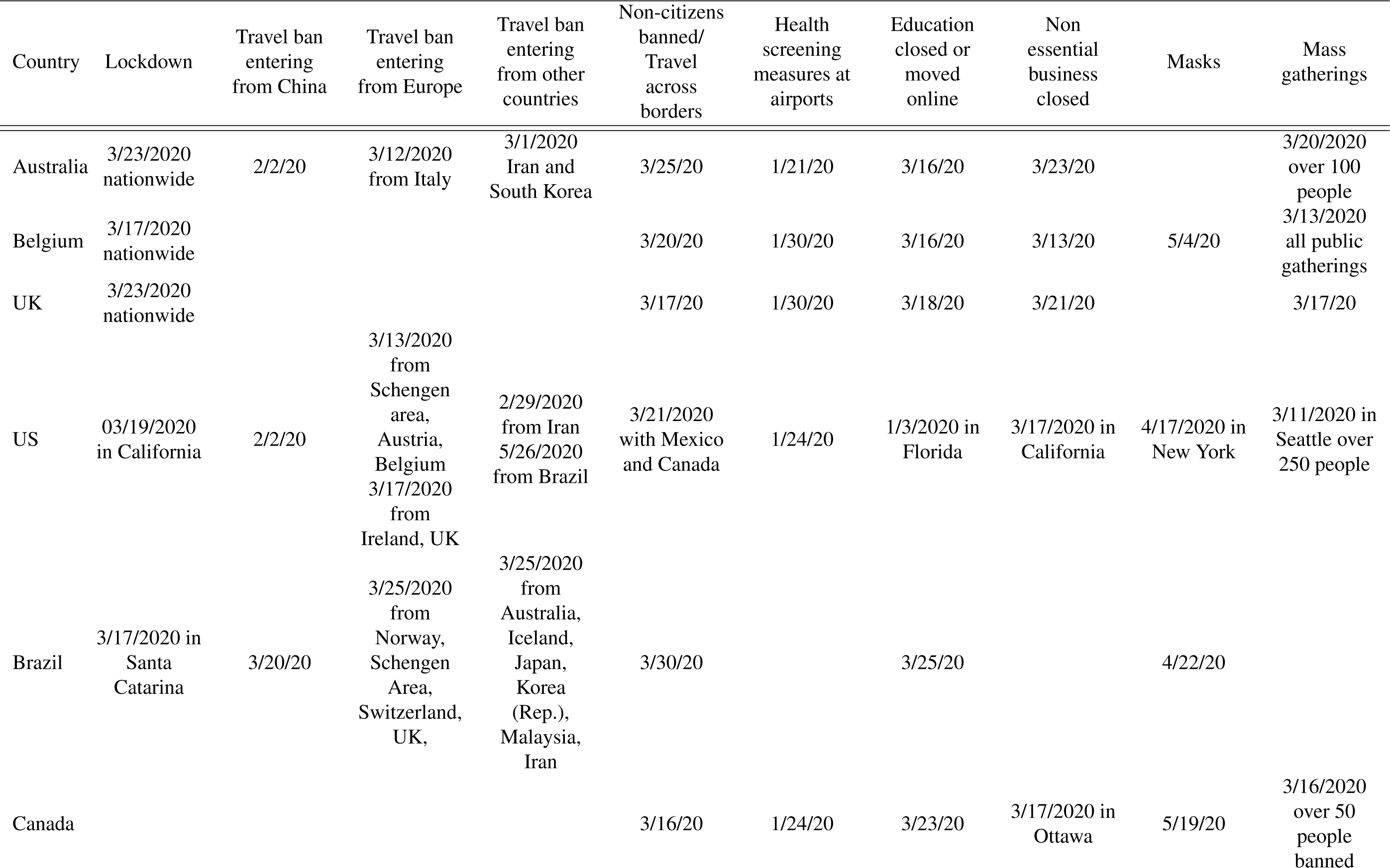
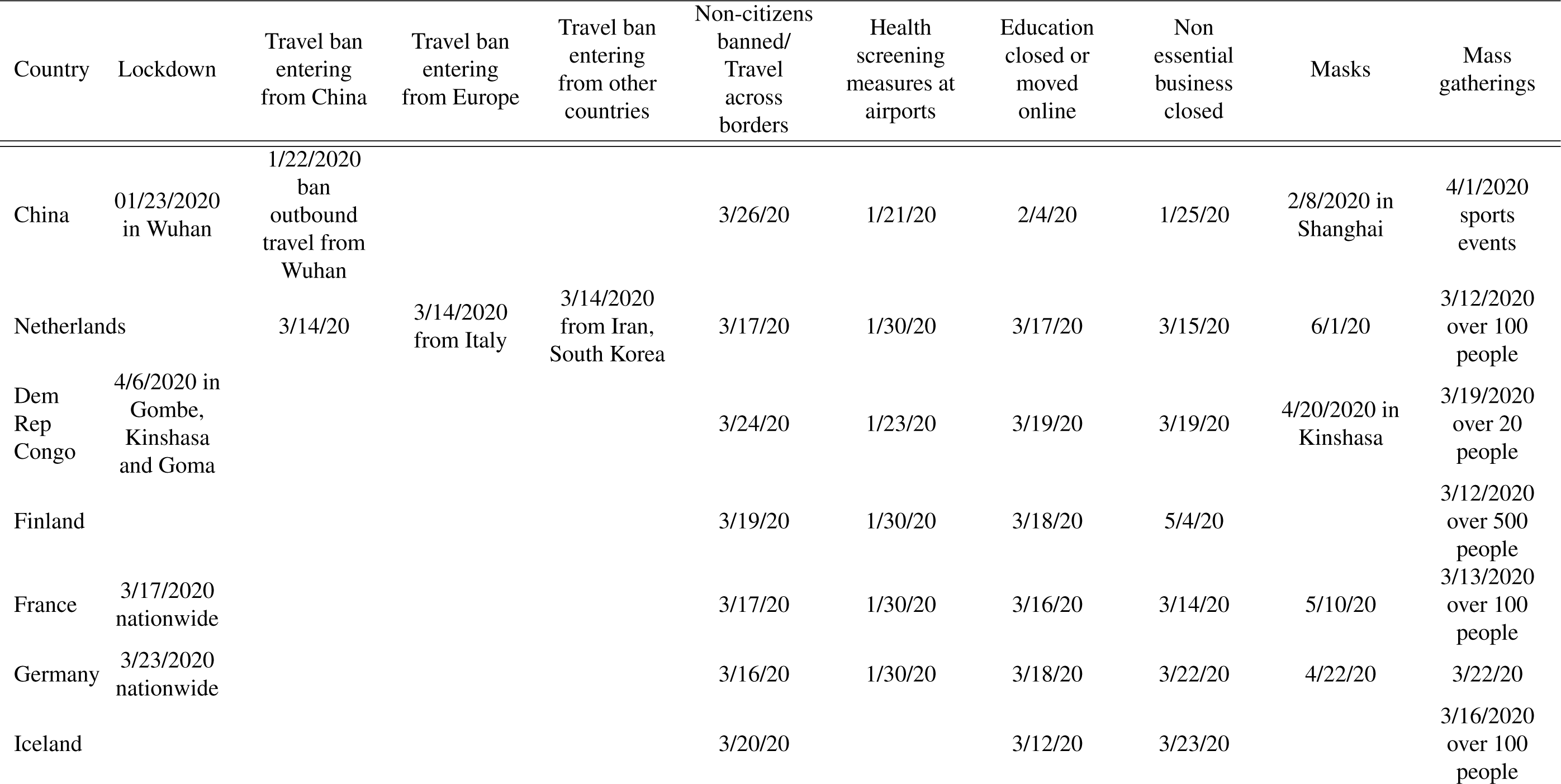
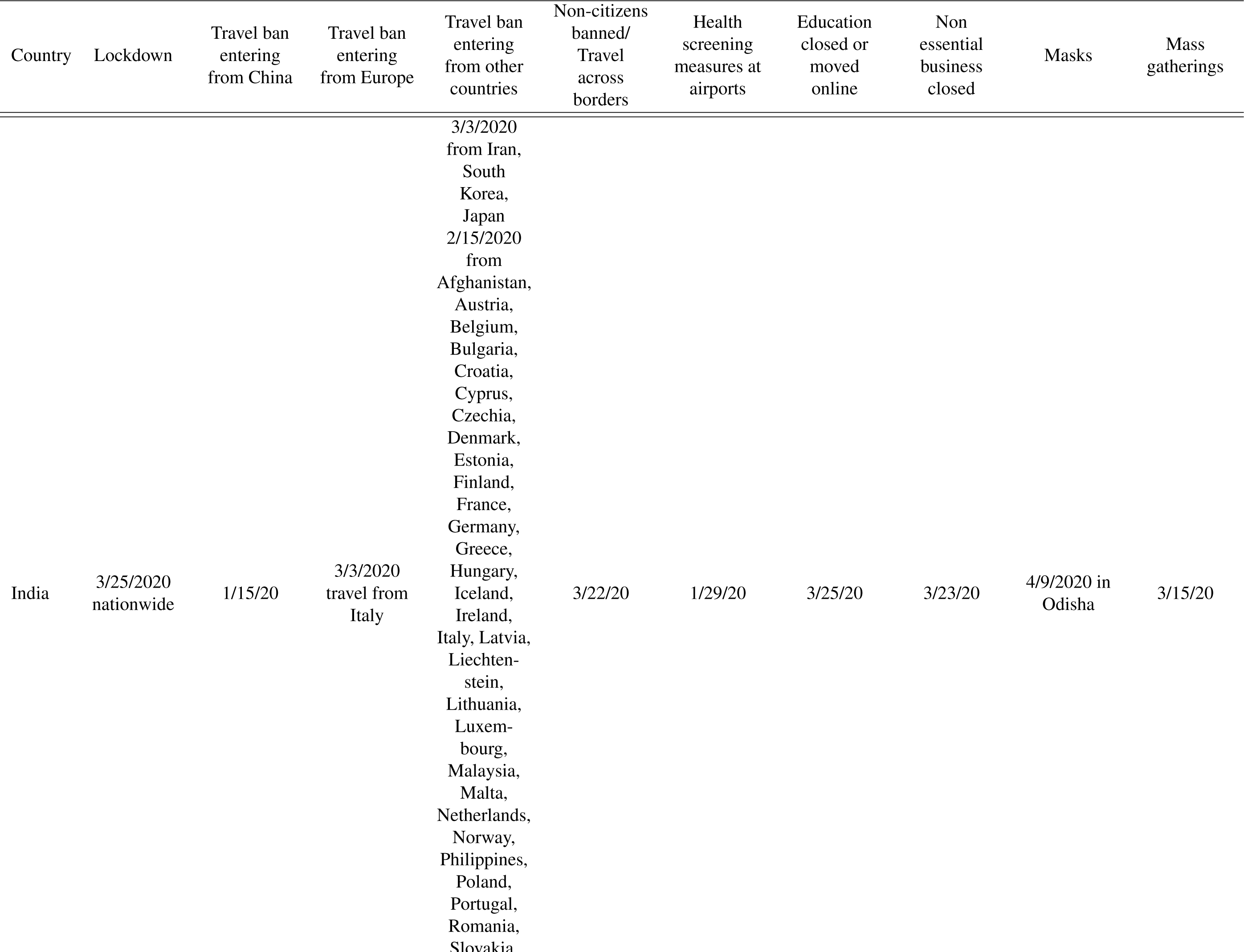
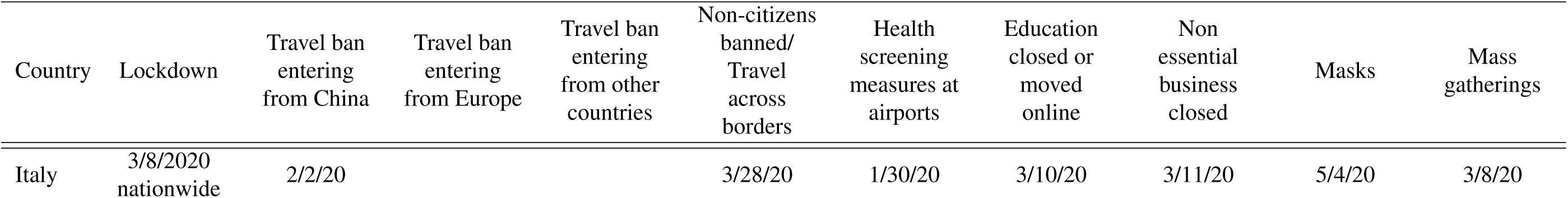
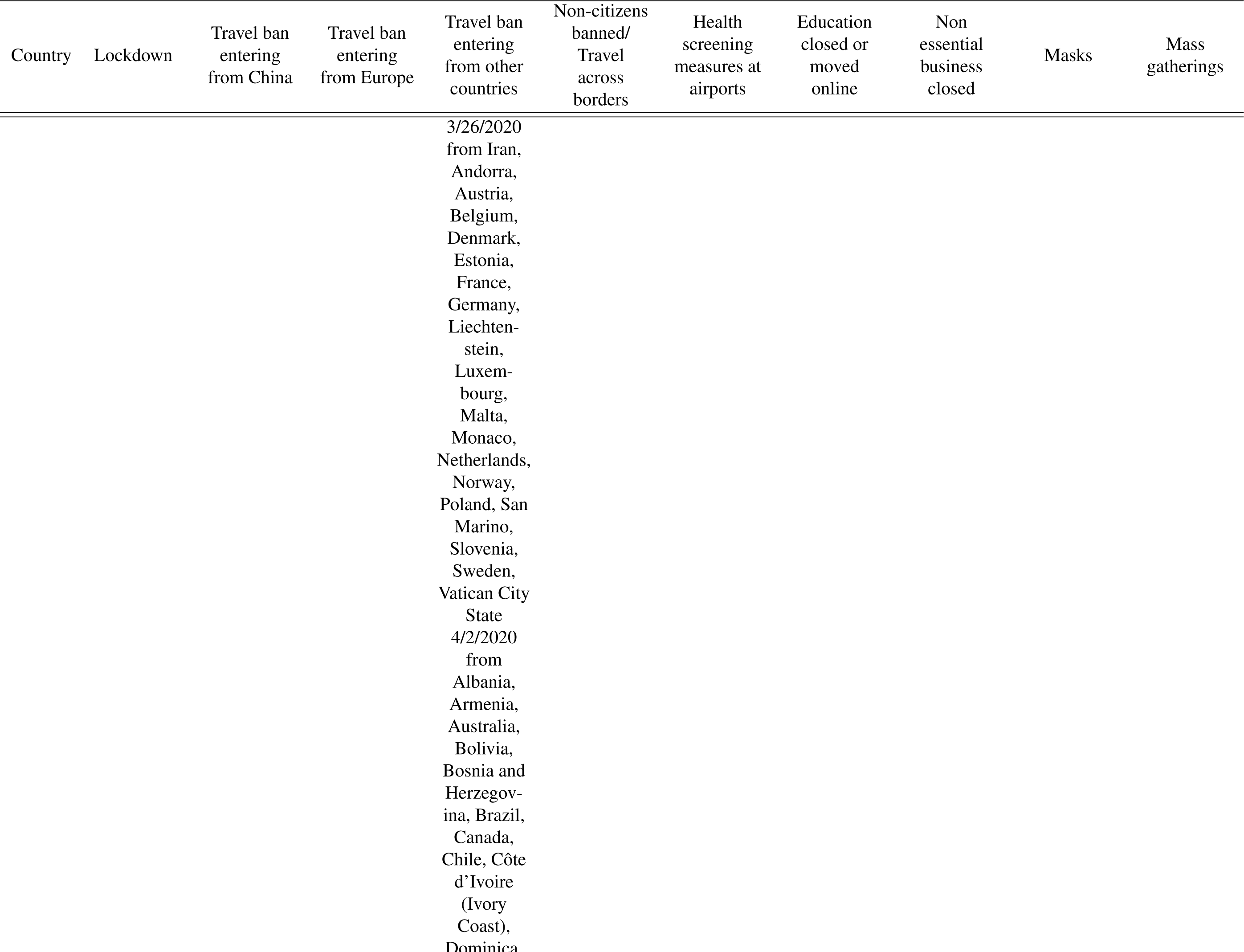
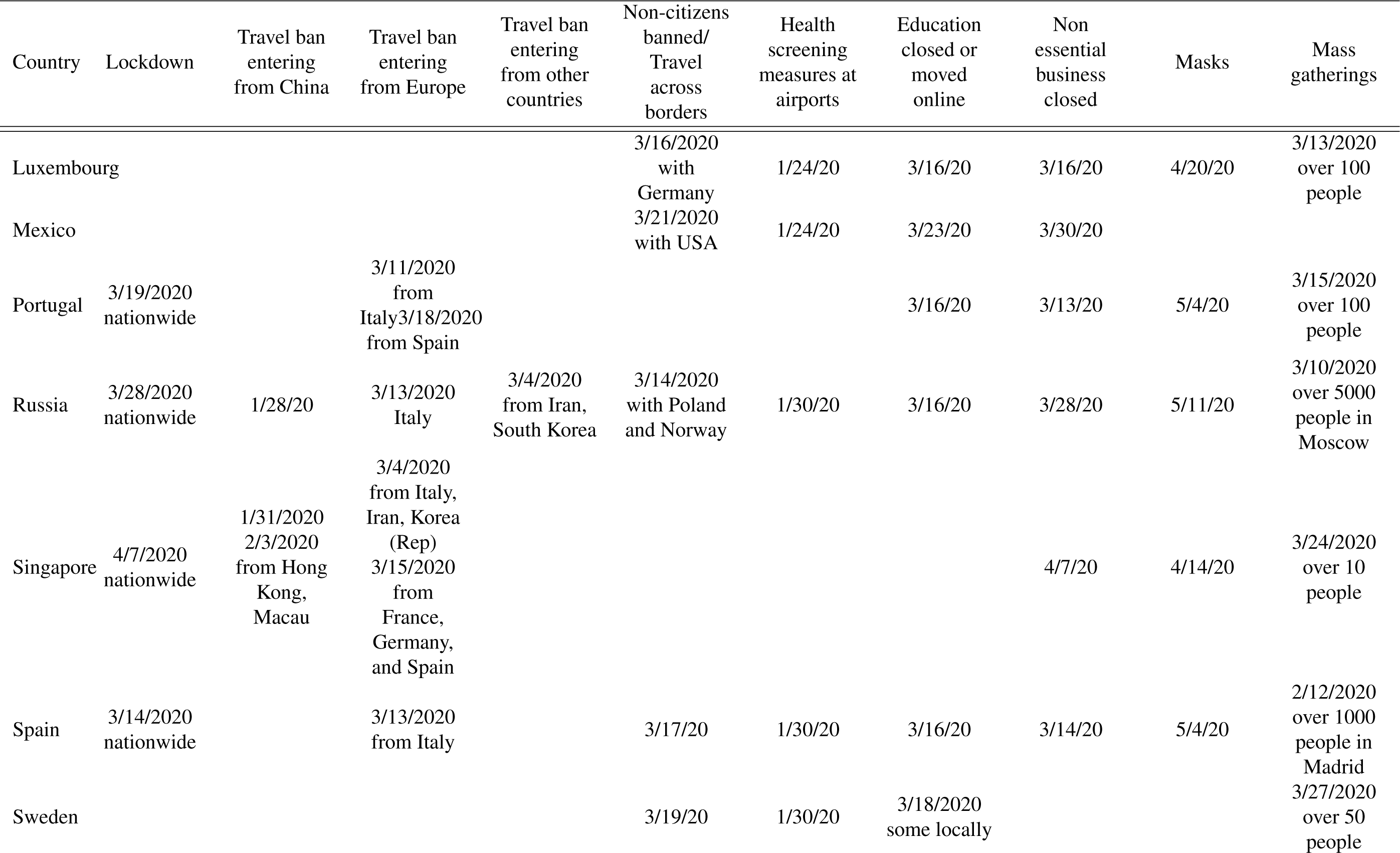
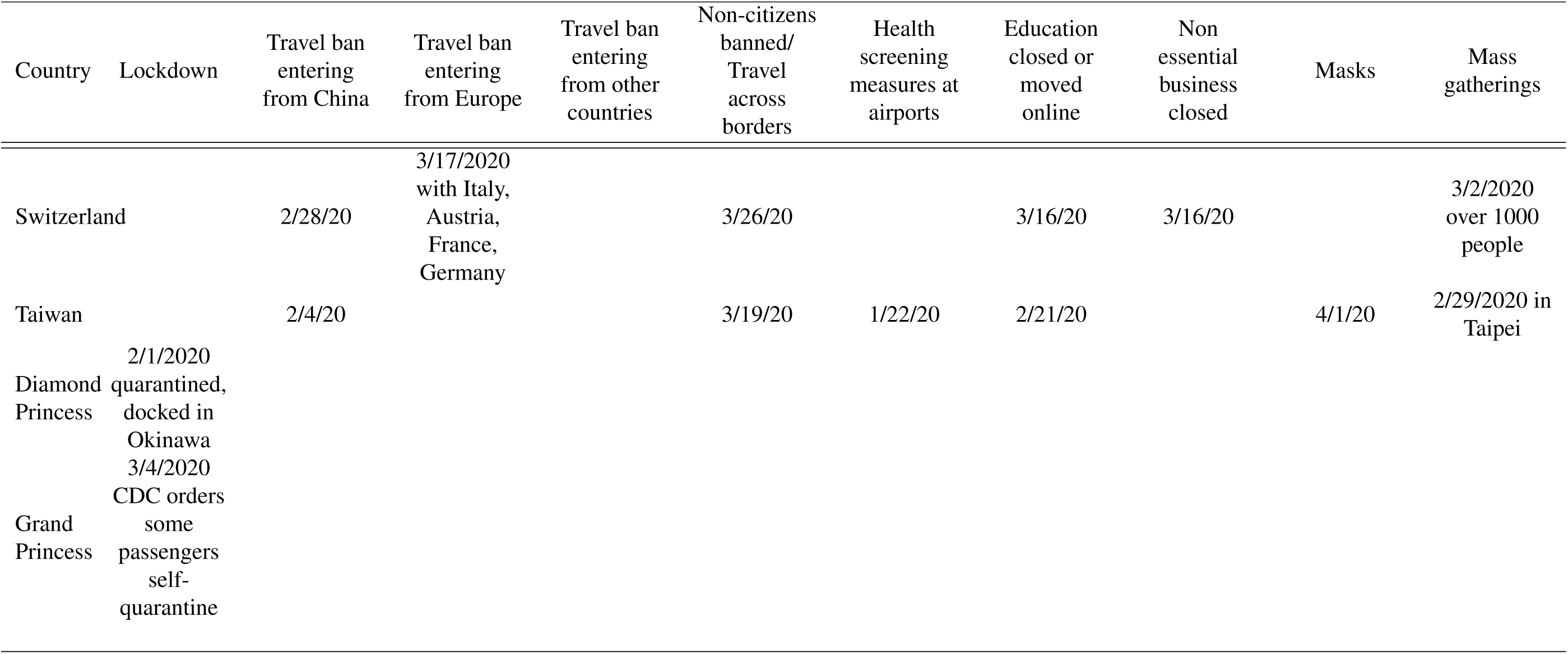
Travel restriction data by country derived from various sources (see Materials and Methods)

**Table S2:**
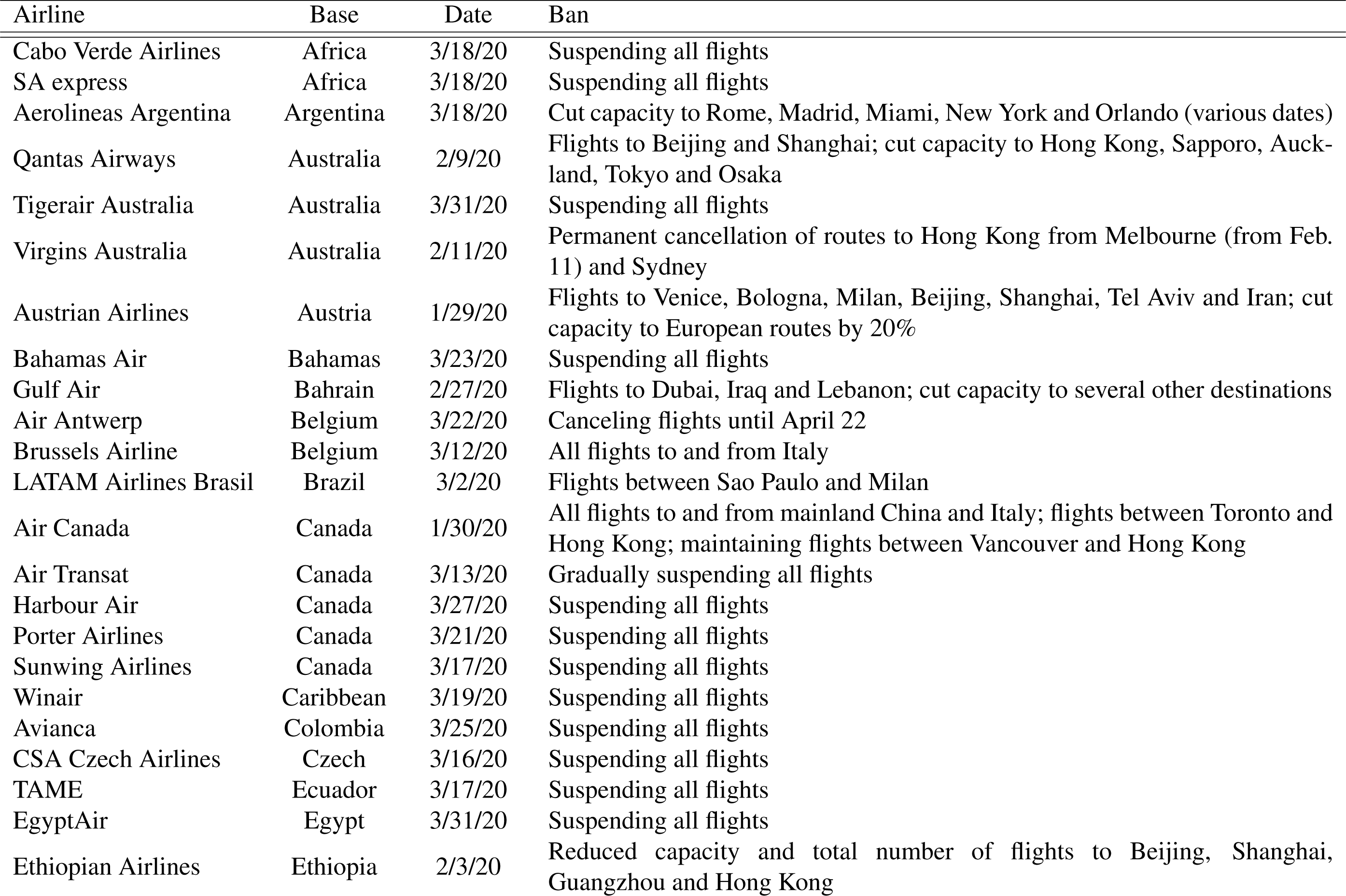
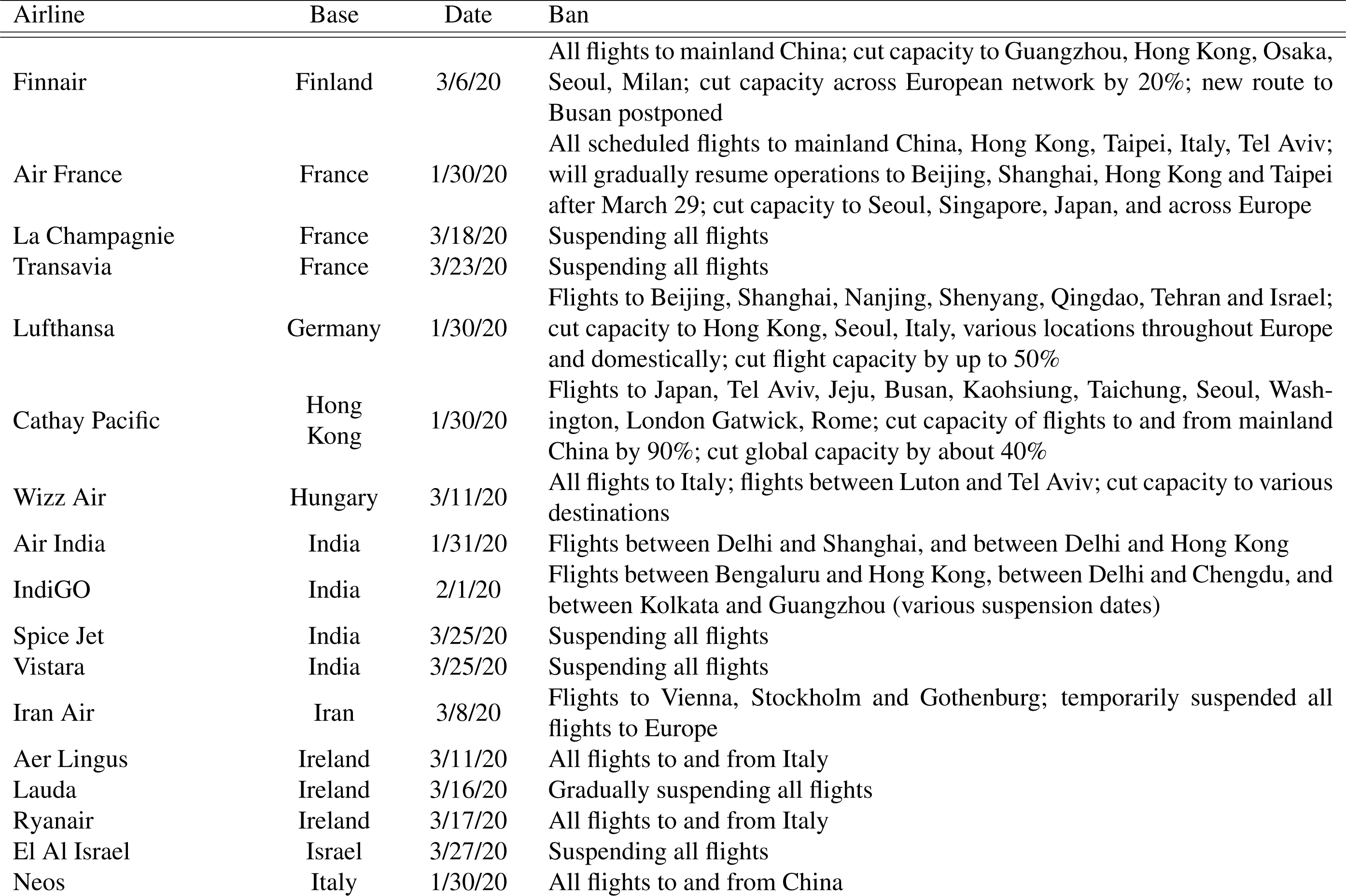
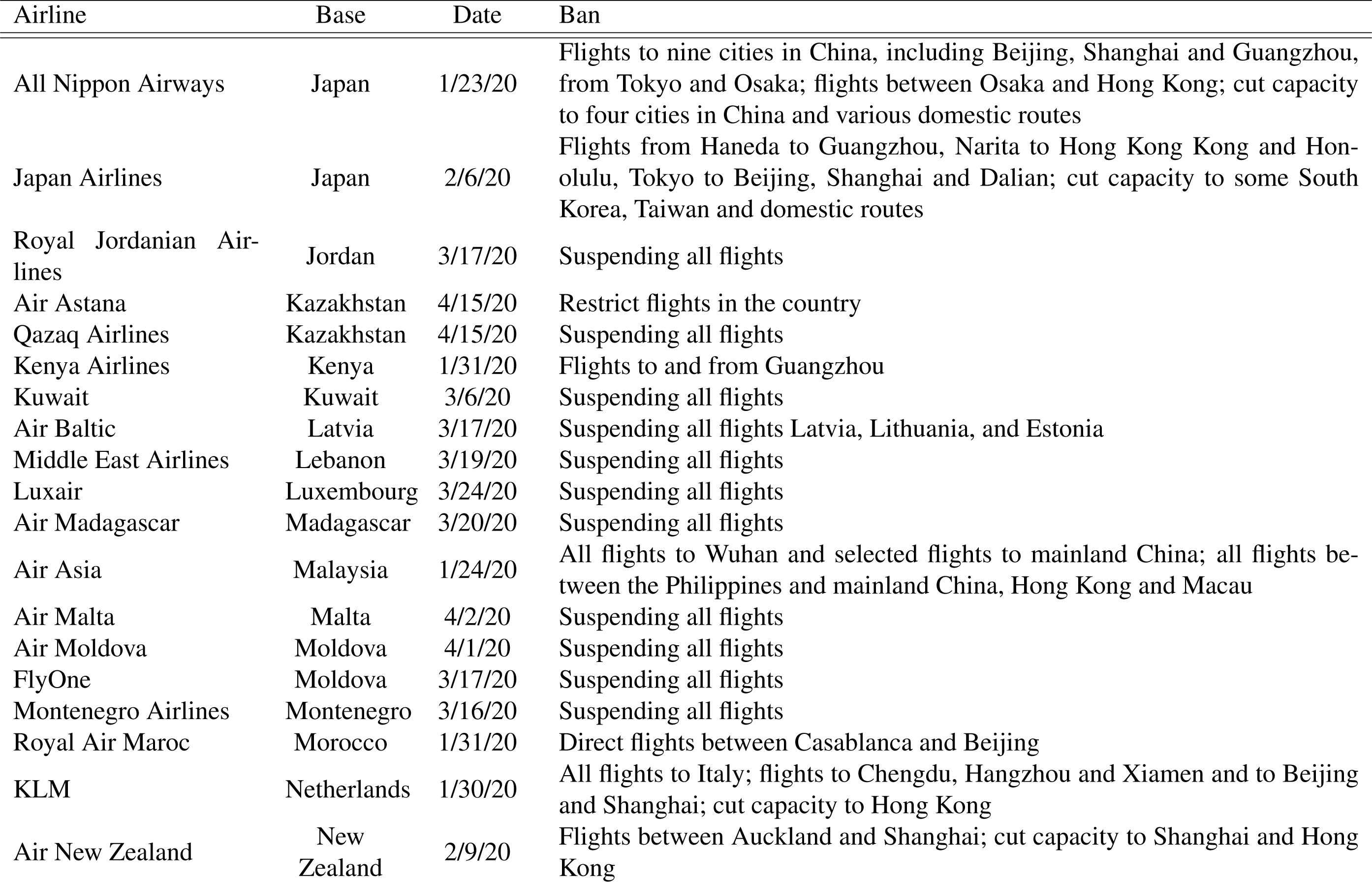
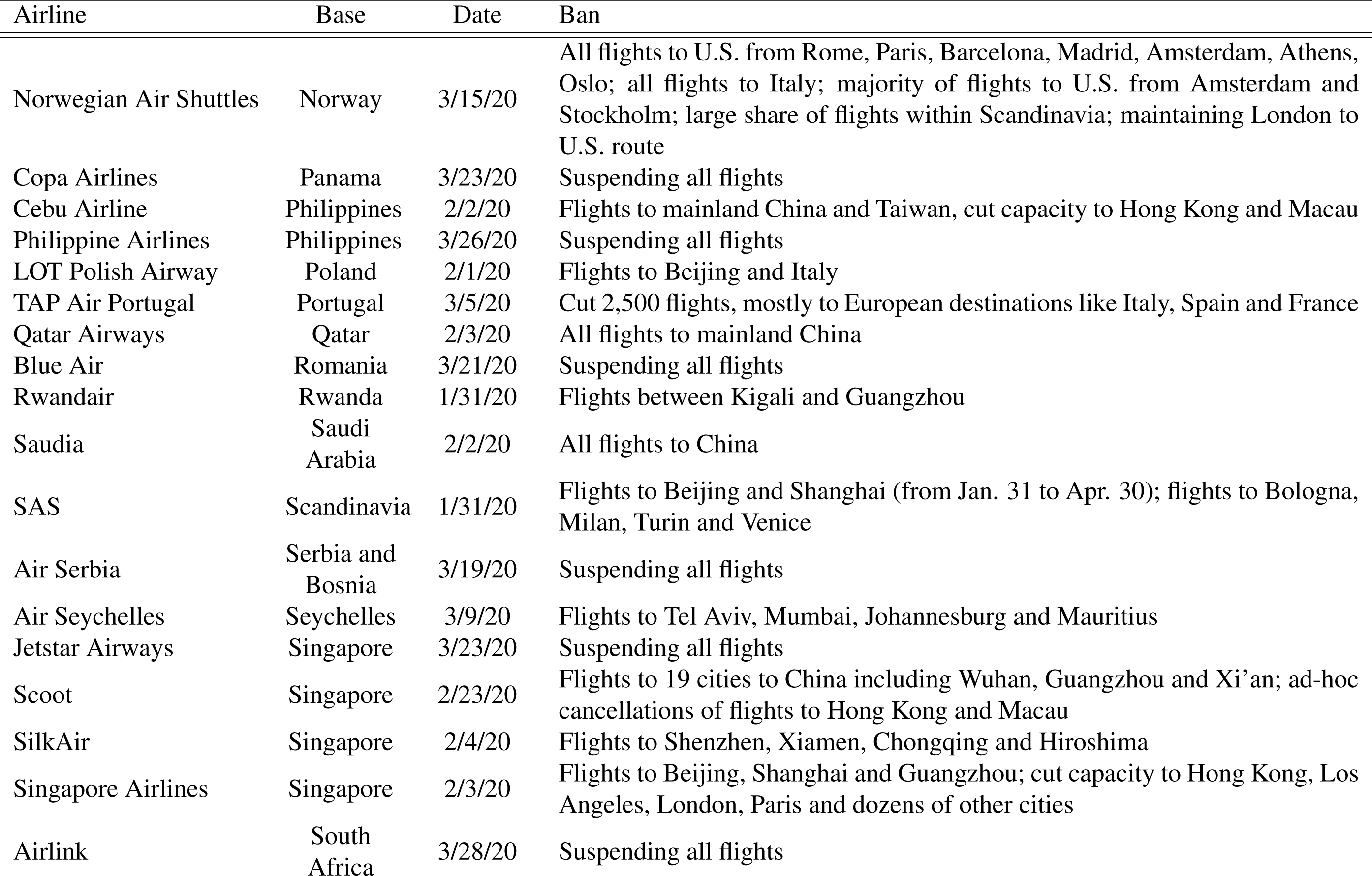
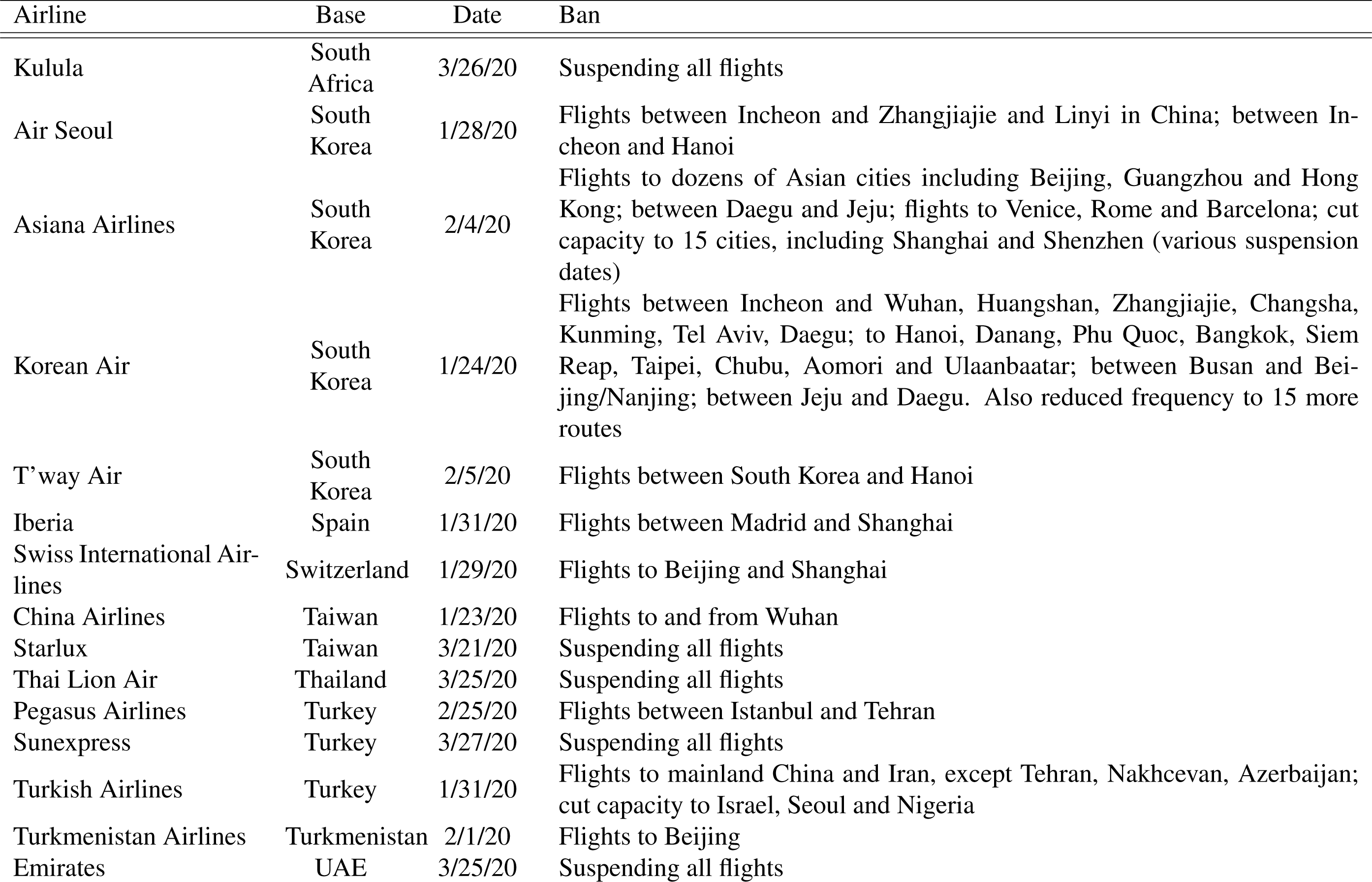
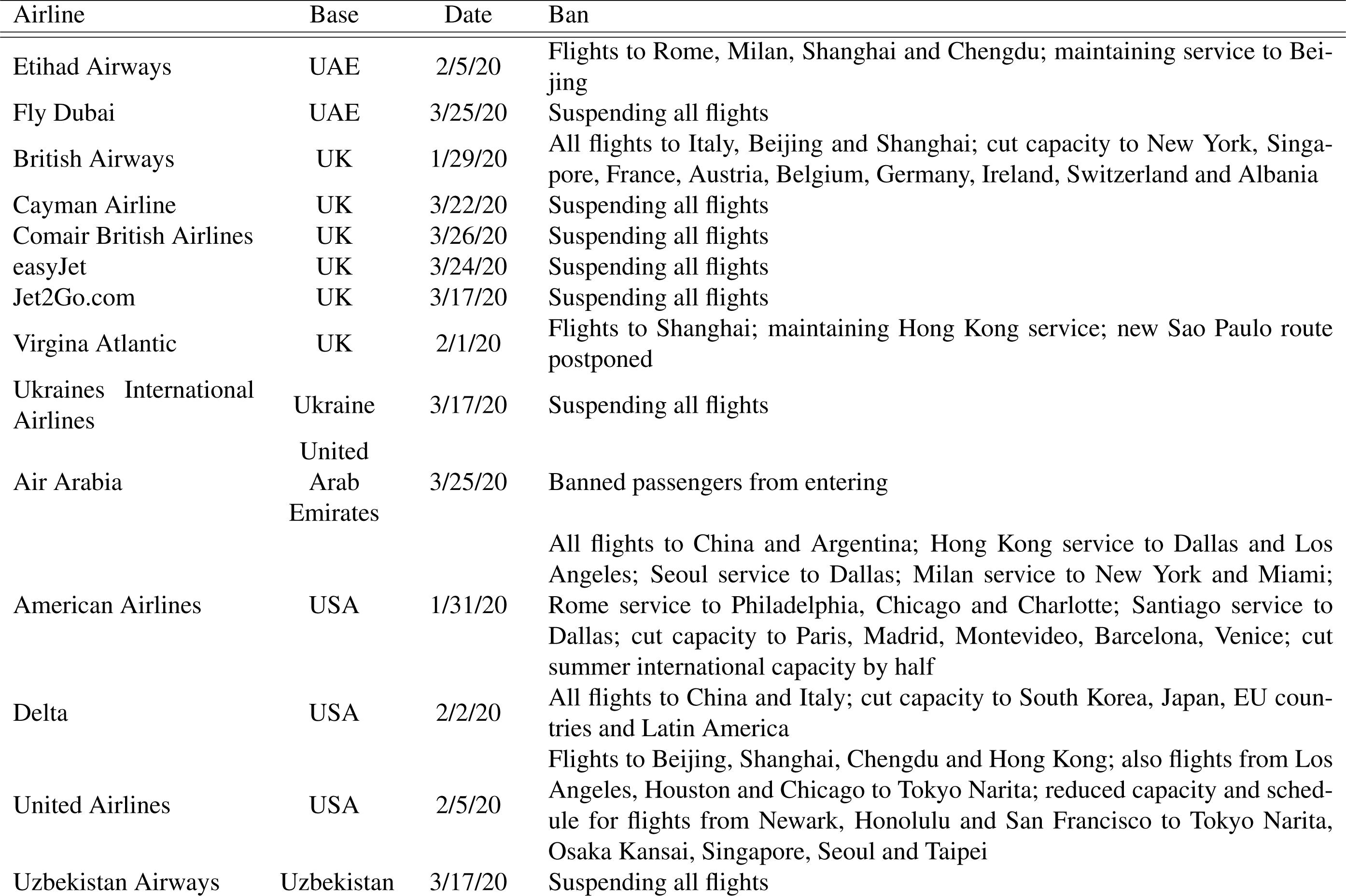
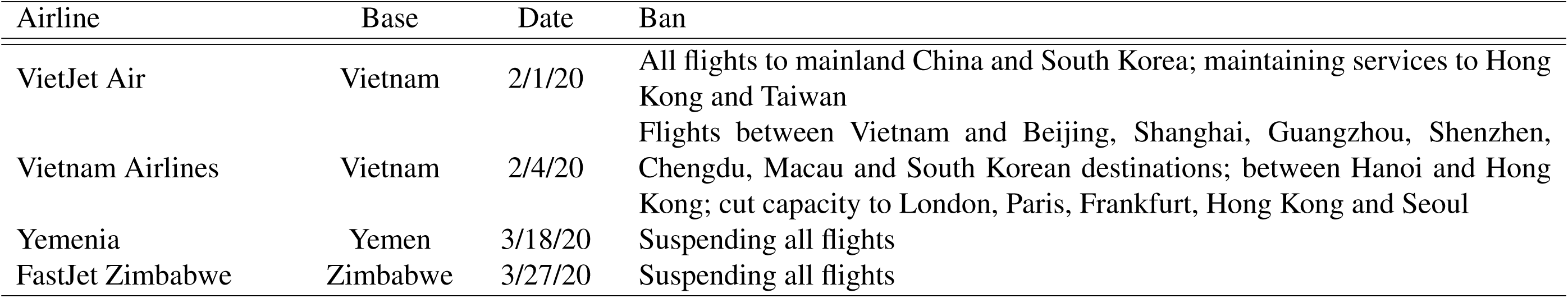
Airline travel data obtained from various sources (see Materials and Methods).

**Table S3:** GISAID sequence ID<u>s collected on April 25</u>, 2020.

| ID |
| --- |
| EPI_ISL_417541 |
| EPI_ISL_419448 |
| EPI_ISL_424900 |
| EPI_ISL_419301 |
| EPI_ISL_428832 |
| EPI_ISL_413850 |
| EPI_ISL_419738 |
| EPI_ISL_424528 |
| EPI_ISL_415529 |
| EPI_ISL_421922 |
| EPI_ISL_421900 |
| EPI_ISL_427334 |
| EPI_ISL_414562 |
| EPI_ISL_404227 |
| EPI_ISL_422032 |
| EPI_ISL_417234 |
| EPI_ISL_426763 |
| EPI_ISL_420731 |
| EPI_ISL_417386 |
| EPI_ISL_417267 |
| EPI_ISL_422888 |
| EPI_ISL_429882 |
| EPI_ISL_419824 |
| EPI_ISL_424227 |
| EPI_ISL_426307 |
| EPI_ISL_425762 |
| EPI_ISL_417813 |
| EPI_ISL_422602 |
| EPI_ISL_427208 |
| EPI_ISL_424092 |
| EPI_ISL_421934 |
| EPI_ISL_423205 |
| EPI_ISL_422019 |
| EPI_ISL_421746 |
| EPI_ISL_420003 |
| EPI_ISL_424215 |
| EPI_ISL_421411 |
| EPI_ISL_416718 |

Table S3: GISAID sequence IDs collected on April 25, 2020.
| ID |
| --- |
| EPI_ISL_417669 |
| EPI_ISL_402129 |
| EPI_ISL_425857 |
| EPI_ISL_426978 |
| EPI_ISL_423952 |
| EPI_ISL_420773 |
| EPI_ISL_423332 |
| EPI_ISL_419436 |
| EPI_ISL_420691 |
| EPI_ISL_420516 |
| EPI_ISL_426689 |
| EPI_ISL_429483 |
| EPI_ISL_420174 |
| EPI_ISL_419674 |
| EPI_ISL_429022 |
| EPI_ISL_429089 |
| EPI_ISL_420569 |
| EPI_ISL_423642 |
| EPI_ISL_426852 |
| EPI_ISL_422042 |
| EPI_ISL_423384 |
| EPI_ISL_425872 |
| EPI_ISL_426979 |
| EPI_ISL_417466 |
| EPI_ISL_422071 |
| EPI_ISL_428319 |
| EPI_ISL_423221 |
| EPI_ISL_422421 |
| EPI_ISL_419907 |
| EPI_ISL_427381 |
| EPI_ISL_424498 |
| EPI_ISL_427764 |
| EPI_ISL_418433 |
| EPI_ISL_426642 |
| EPI_ISL_417348 |
| EPI_ISL_424379 |
| EPI_ISL_430097 |
| EPI_ISL_419582 |

Table S3: GISAID sequence IDs collected on April 25, 2020.
| ID |
| --- |
| EPI_ISL_424482 |
| EPI_ISL_428262 |
| EPI_ISL_417092 |
| EPI_ISL_429131 |
| EPI_ISL_423161 |
| EPI_ISL_419585 |
| EPI_ISL_418885 |
| EPI_ISL_421314 |
| EPI_ISL_417332 |
| EPI_ISL_418823 |
| EPI_ISL_426941 |
| EPI_ISL_416589 |
| EPI_ISL_426085 |
| EPI_ISL_423310 |
| EPI_ISL_423375 |
| EPI_ISL_418290 |
| EPI_ISL_419935 |
| EPI_ISL_417878 |
| EPI_ISL_420478 |
| EPI_ISL_421879 |
| EPI_ISL_414526 |
| EPI_ISL_419902 |
| EPI_ISL_428272 |
| EPI_ISL_422977 |
| EPI_ISL_417181 |
| EPI_ISL_425048 |
| EPI_ISL_411952 |
| EPI_ISL_427497 |
| EPI_ISL_421427 |
| EPI_ISL_427199 |
| EPI_ISL_425596 |
| EPI_ISL_415532 |
| EPI_ISL_425238 |
| EPI_ISL_420114 |
| EPI_ISL_419532 |
| EPI_ISL_425056 |
| EPI_ISL_419712 |
| EPI_ISL_420612 |

Table S3: GISAID sequence IDs collected on April 25, 2020.
| ID |
| --- |
| EPI_ISL_423000 |
| EPI_ISL_416711 |
| EPI_ISL_417514 |
| EPI_ISL_416482 |
| EPI_ISL_418641 |
| EPI_ISL_421479 |
| EPI_ISL_418033 |
| EPI_ISL_422081 |
| EPI_ISL_424032 |
| EPI_ISL_421004 |
| EPI_ISL_419265 |
| EPI_ISL_425694 |
| EPI_ISL_424476 |
| EPI_ISL_429155 |
| EPI_ISL_413853 |
| EPI_ISL_417524 |
| EPI_ISL_424050 |
| EPI_ISL_429067 |
| EPI_ISL_426729 |
| EPI_ISL_425242 |
| EPI_ISL_416365 |
| EPI_ISL_417301 |
| EPI_ISL_420847 |
| EPI_ISL_422330 |
| EPI_ISL_421260 |
| EPI_ISL_421630 |
| EPI_ISL_425206 |
| EPI_ISL_421738 |
| EPI_ISL_423474 |
| EPI_ISL_426954 |
| EPI_ISL_419559 |
| EPI_ISL_426774 |
| EPI_ISL_417663 |
| EPI_ISL_420749 |
| EPI_ISL_422889 |
| EPI_ISL_427488 |
| EPI_ISL_420477 |
| EPI_ISL_426811 |

Table S3: GISAID sequence IDs collected on April 25, 2020.
| ID |
| --- |
| EPI_ISL_424221 |
| EPI_ISL_427263 |
| EPI_ISL_406531 |
| EPI_ISL_416699 |
| EPI_ISL_421617 |
| EPI_ISL_418896 |
| EPI_ISL_425383 |
| EPI_ISL_418967 |
| EPI_ISL_423477 |
| EPI_ISL_417690 |
| EPI_ISL_429162 |
| EPI_ISL_422927 |
| EPI_ISL_416676 |
| EPI_ISL_427160 |
| EPI_ISL_419499 |
| EPI_ISL_417937 |
| EPI_ISL_418894 |
| EPI_ISL_427088 |
| EPI_ISL_417224 |
| EPI_ISL_424317 |
| EPI_ISL_417850 |
| EPI_ISL_425535 |
| EPI_ISL_427247 |
| EPI_ISL_425688 |
| EPI_ISL_420360 |
| EPI_ISL_420549 |
| EPI_ISL_417222 |
| EPI_ISL_417007 |
| EPI_ISL_426092 |
| EPI_ISL_415538 |
| EPI_ISL_418931 |
| EPI_ISL_421388 |
| EPI_ISL_423135 |
| EPI_ISL_426965 |
| EPI_ISL_428233 |
| EPI_ISL_422397 |
| EPI_ISL_424546 |
| EPI_ISL_424642 |

Table S3: GISAID sequence IDs collected on April 25, 2020.
| ID |
| --- |
| EPI_ISL_429491 |
| EPI_ISL_421691 |
| EPI_ISL_429782 |
| EPI_ISL_419561 |
| EPI_ISL_425988 |
| EPI_ISL_419936 |
| EPI_ISL_422939 |
| EPI_ISL_423638 |
| EPI_ISL_429531 |
| EPI_ISL_424301 |
| EPI_ISL_427290 |
| EPI_ISL_422913 |
| EPI_ISL_425904 |
| EPI_ISL_415595 |
| EPI_ISL_421002 |
| EPI_ISL_416444 |
| EPI_ISL_422128 |
| EPI_ISL_421560 |
| EPI_ISL_422527 |
| EPI_ISL_419749 |
| EPI_ISL_417122 |
| EPI_ISL_422567 |
| EPI_ISL_419834 |
| EPI_ISL_423435 |
| EPI_ISL_417144 |
| EPI_ISL_415743 |
| EPI_ISL_429504 |
| EPI_ISL_427549 |
| EPI_ISL_429744 |
| EPI_ISL_418965 |
| EPI_ISL_422667 |
| EPI_ISL_417163 |
| EPI_ISL_417350 |
| EPI_ISL_422839 |
| EPI_ISL_427773 |
| EPI_ISL_420236 |
| EPI_ISL_429717 |
| EPI_ISL_427744 |

Table S3: GISAID sequence IDs collected on April 25, 2020.
| ID |
| --- |
| EPI_ISL_423968 |
| EPI_ISL_425362 |
| EPI_ISL_422747 |
| EPI_ISL_415650 |
| EPI_ISL_422917 |
| EPI_ISL_420346 |
| EPI_ISL_417022 |
| EPI_ISL_417364 |
| EPI_ISL_416450 |
| EPI_ISL_427474 |
| EPI_ISL_427353 |
| EPI_ISL_423096 |
| EPI_ISL_423063 |
| EPI_ISL_423125 |
| EPI_ISL_422949 |
| EPI_ISL_427110 |
| EPI_ISL_420527 |
| EPI_ISL_418388 |
| EPI_ISL_416698 |
| EPI_ISL_424609 |
| EPI_ISL_418770 |
| EPI_ISL_414486 |
| EPI_ISL_428254 |
| EPI_ISL_418192 |
| EPI_ISL_422880 |
| EPI_ISL_425055 |
| EPI_ISL_418114 |
| EPI_ISL_423854 |
| EPI_ISL_427787 |
| EPI_ISL_426502 |
| EPI_ISL_429281 |
| EPI_ISL_425820 |
| EPI_ISL_419173 |
| EPI_ISL_430021 |
| EPI_ISL_425682 |
| EPI_ISL_420585 |
| EPI_ISL_413653 |
| EPI_ISL_430070 |

Table S3: GISAID sequence IDs collected on April 25, 2020.
| ID |
| --- |
| EPI_ISL_424342 |
| EPI_ISL_426306 |
| EPI_ISL_425430 |
| EPI_ISL_424574 |
| EPI_ISL_421289 |
| EPI_ISL_420611 |
| EPI_ISL_414366 |
| EPI_ISL_426617 |
| EPI_ISL_421207 |
| EPI_ISL_420532 |
| EPI_ISL_418294 |
| EPI_ISL_419832 |
| EPI_ISL_421281 |
| EPI_ISL_417361 |
| EPI_ISL_423176 |
| EPI_ISL_417641 |
| EPI_ISL_424499 |
| EPI_ISL_414621 |
| EPI_ISL_423379 |
| EPI_ISL_418971 |
| EPI_ISL_426993 |
| EPI_ISL_428757 |
| EPI_ISL_419460 |
| EPI_ISL_424272 |
| EPI_ISL_414043 |
| EPI_ISL_428468 |
| EPI_ISL_428232 |
| EPI_ISL_418260 |
| EPI_ISL_419568 |
| EPI_ISL_429564 |
| EPI_ISL_421473 |
| EPI_ISL_427365 |
| EPI_ISL_423823 |
| EPI_ISL_425951 |
| EPI_ISL_424354 |
| EPI_ISL_424103 |
| EPI_ISL_424960 |
| EPI_ISL_417584 |

Table S3: GISAID sequence IDs collected on April 25, 2020.
| ID |
| --- |
| EPI_ISL_422323 |
| EPI_ISL_420058 |
| EPI_ISL_420208 |
| EPI_ISL_423124 |
| EPI_ISL_427084 |
| EPI_ISL_425673 |
| EPI_ISL_417481 |
| EPI_ISL_423074 |
| EPI_ISL_417536 |
| EPI_ISL_420248 |
| EPI_ISL_420209 |
| EPI_ISL_413607 |
| EPI_ISL_420483 |
| EPI_ISL_429359 |
| EPI_ISL_423936 |
| EPI_ISL_417193 |
| EPI_ISL_417477 |
| EPI_ISL_418834 |
| EPI_ISL_423560 |
| EPI_ISL_416488 |
| EPI_ISL_424449 |
| EPI_ISL_417246 |
| EPI_ISL_425521 |
| EPI_ISL_427555 |
| EPI_ISL_424162 |
| EPI_ISL_425135 |
| EPI_ISL_412898 |
| EPI_ISL_421855 |
| EPI_ISL_422303 |
| EPI_ISL_429464 |
| EPI_ISL_426555 |
| EPI_ISL_426964 |
| EPI_ISL_418796 |
| EPI_ISL_418240 |
| EPI_ISL_420461 |
| EPI_ISL_429969 |
| EPI_ISL_416368 |
| EPI_ISL_427205 |

Table S3: GISAID sequence IDs collected on April 25, 2020.
| ID |
| --- |
| EPI_ISL_426685 |
| EPI_ISL_426014 |
| EPI_ISL_418218 |
| EPI_ISL_413858 |
| EPI_ISL_427786 |
| EPI_ISL_420798 |
| EPI_ISL_424956 |
| EPI_ISL_419984 |
| EPI_ISL_419899 |
| EPI_ISL_425984 |
| EPI_ISL_429485 |
| EPI_ISL_429282 |
| EPI_ISL_421291 |
| EPI_ISL_415710 |
| EPI_ISL_429467 |
| EPI_ISL_418346 |
| EPI_ISL_425913 |
| EPI_ISL_414633 |
| EPI_ISL_420460 |
| EPI_ISL_429097 |
| EPI_ISL_415479 |
| EPI_ISL_429142 |
| EPI_ISL_417816 |
| EPI_ISL_417836 |
| EPI_ISL_414539 |
| EPI_ISL_426288 |
| EPI_ISL_416427 |
| EPI_ISL_418652 |
| EPI_ISL_425891 |
| EPI_ISL_422651 |
| EPI_ISL_429082 |
| EPI_ISL_420631 |
| EPI_ISL_419564 |
| EPI_ISL_421355 |
| EPI_ISL_421450 |
| EPI_ISL_426471 |
| EPI_ISL_407976 |
| EPI_ISL_418284 |

Table S3: GISAID sequence IDs collected on April 25, 2020.
| ID |
| --- |
| EPI_ISL_428373 |
| EPI_ISL_427789 |
| EPI_ISL_427650 |
| EPI_ISL_420120 |
| EPI_ISL_429511 |
| EPI_ISL_419584 |
| EPI_ISL_424880 |
| EPI_ISL_419720 |
| EPI_ISL_429305 |
| EPI_ISL_414586 |
| EPI_ISL_421417 |
| EPI_ISL_422427 |
| EPI_ISL_418646 |
| EPI_ISL_418159 |
| EPI_ISL_428359 |
| EPI_ISL_423347 |
| EPI_ISL_422871 |
| EPI_ISL_419767 |
| EPI_ISL_426527 |
| EPI_ISL_426564 |
| EPI_ISL_421755 |
| EPI_ISL_418155 |
| EPI_ISL_427034 |
| EPI_ISL_426750 |
| EPI_ISL_420145 |
| EPI_ISL_418826 |
| EPI_ISL_426820 |
| EPI_ISL_428253 |
| EPI_ISL_427017 |
| EPI_ISL_417313 |
| EPI_ISL_420930 |
| EPI_ISL_414564 |
| EPI_ISL_420137 |
| EPI_ISL_419960 |
| EPI_ISL_429065 |
| EPI_ISL_417795 |
| EPI_ISL_416491 |
| EPI_ISL_426082 |

Table S3: GISAID sequence IDs collected on April 25, 2020.
| ID |
| --- |
| EPI_ISL_420200 |
| EPI_ISL_429561 |
| EPI_ISL_423358 |
| EPI_ISL_423212 |
| EPI_ISL_419218 |
| EPI_ISL_425170 |
| EPI_ISL_419524 |
| EPI_ISL_421683 |
| EPI_ISL_427073 |
| EPI_ISL_415977 |
| EPI_ISL_417638 |
| EPI_ISL_429622 |
| EPI_ISL_423592 |
| EPI_ISL_429607 |
| EPI_ISL_419971 |
| EPI_ISL_424065 |
| EPI_ISL_422938 |
| EPI_ISL_418188 |
| EPI_ISL_418045 |
| EPI_ISL_427324 |
| EPI_ISL_425187 |
| EPI_ISL_426567 |
| EPI_ISL_418712 |
| EPI_ISL_421573 |
| EPI_ISL_426091 |
| EPI_ISL_414438 |
| EPI_ISL_419254 |
| EPI_ISL_430019 |
| EPI_ISL_425609 |
| EPI_ISL_422076 |
| EPI_ISL_419482 |
| EPI_ISL_417854 |
| EPI_ISL_419848 |
| EPI_ISL_429015 |
| EPI_ISL_421729 |
| EPI_ISL_424262 |
| EPI_ISL_416524 |
| EPI_ISL_424008 |

Table S3: GISAID sequence IDs collected on April 25, 2020.
| ID |
| --- |
| EPI_ISL_428325 |
| EPI_ISL_421243 |
| EPI_ISL_418720 |
| EPI_ISL_413997 |
| EPI_ISL_417560 |
| EPI_ISL_423044 |
| EPI_ISL_425287 |
| EPI_ISL_427674 |
| EPI_ISL_423122 |
| EPI_ISL_417194 |
| EPI_ISL_422057 |
| EPI_ISL_416721 |
| EPI_ISL_416744 |
| EPI_ISL_429346 |
| EPI_ISL_428347 |
| EPI_ISL_429465 |
| EPI_ISL_428900 |
| EPI_ISL_426074 |
| EPI_ISL_416334 |
| EPI_ISL_423692 |
| EPI_ISL_428800 |
| EPI_ISL_418665 |
| EPI_ISL_425470 |
| EPI_ISL_422746 |
| EPI_ISL_417004 |
| EPI_ISL_419517 |
| EPI_ISL_427531 |
| EPI_ISL_424938 |
| EPI_ISL_423027 |
| EPI_ISL_425784 |
| EPI_ISL_426768 |
| EPI_ISL_424937 |
| EPI_ISL_425511 |
| EPI_ISL_414692 |
| EPI_ISL_428267 |
| EPI_ISL_417426 |
| EPI_ISL_420040 |
| EPI_ISL_418220 |

Table S3: GISAID sequence IDs collected on April 25, 2020.
| ID |
| --- |
| EPI_ISL_422164 |
| EPI_ISL_411950 |
| EPI_ISL_420078 |
| EPI_ISL_421743 |
| EPI_ISL_429884 |
| EPI_ISL_422694 |
| EPI_ISL_420326 |
| EPI_ISL_425720 |
| EPI_ISL_419260 |
| EPI_ISL_422557 |
| EPI_ISL_421432 |
| EPI_ISL_422944 |
| EPI_ISL_417675 |
| EPI_ISL_424059 |
| EPI_ISL_418869 |
| EPI_ISL_418882 |
| EPI_ISL_422222 |
| EPI_ISL_424974 |
| EPI_ISL_427545 |
| EPI_ISL_423216 |
| EPI_ISL_418739 |
| EPI_ISL_423856 |
| EPI_ISL_429634 |
| EPI_ISL_421320 |
| EPI_ISL_427455 |
| EPI_ISL_427071 |
| EPI_ISL_429565 |
| EPI_ISL_424857 |
| EPI_ISL_427164 |
| EPI_ISL_422174 |
| EPI_ISL_422115 |
| EPI_ISL_423941 |
| EPI_ISL_422855 |
| EPI_ISL_426059 |
| EPI_ISL_424846 |
| EPI_ISL_418016 |
| EPI_ISL_427106 |
| EPI_ISL_424209 |

Table S3: GISAID sequence IDs collected on April 25, 2020.
| ID |
| --- |
| EPI_ISL_417296 |
| EPI_ISL_420570 |
| EPI_ISL_424186 |
| EPI_ISL_426460 |
| EPI_ISL_426653 |
| EPI_ISL_428878 |
| EPI_ISL_420842 |
| EPI_ISL_425455 |
| EPI_ISL_423320 |
| EPI_ISL_420305 |
| EPI_ISL_418768 |
| EPI_ISL_418935 |
| EPI_ISL_426164 |
| EPI_ISL_427692 |
| EPI_ISL_418717 |
| EPI_ISL_425923 |
| EPI_ISL_422317 |
| EPI_ISL_420505 |
| EPI_ISL_416737 |
| EPI_ISL_429583 |
| EPI_ISL_416694 |
| EPI_ISL_425053 |
| EPI_ISL_421430 |
| EPI_ISL_429008 |
| EPI_ISL_423817 |
| EPI_ISL_406970 |
| EPI_ISL_421612 |
| EPI_ISL_415602 |
| EPI_ISL_427127 |
| EPI_ISL_414550 |
| EPI_ISL_417270 |
| EPI_ISL_429558 |
| EPI_ISL_424606 |
| EPI_ISL_429569 |
| EPI_ISL_424521 |
| EPI_ISL_413652 |
| EPI_ISL_423596 |
| EPI_ISL_423663 |

Table S3: GISAID sequence IDs collected on April 25, 2020.
| ID |
| --- |
| EPI_ISL_419417 |
| EPI_ISL_419589 |
| EPI_ISL_420117 |
| EPI_ISL_422767 |
| EPI_ISL_426695 |
| EPI_ISL_420010 |
| EPI_ISL_415507 |
| EPI_ISL_421847 |
| EPI_ISL_426683 |
| EPI_ISL_422849 |
| EPI_ISL_426928 |
| EPI_ISL_429631 |
| EPI_ISL_420511 |
| EPI_ISL_429643 |
| EPI_ISL_423639 |
| EPI_ISL_420670 |
| EPI_ISL_421909 |
| EPI_ISL_421453 |
| EPI_ISL_427663 |
| EPI_ISL_420224 |
| EPI_ISL_421311 |
| EPI_ISL_413518 |
| EPI_ISL_421705 |
| EPI_ISL_424041 |
| EPI_ISL_419506 |
| EPI_ISL_414471 |
| EPI_ISL_423693 |
| EPI_ISL_417655 |
| EPI_ISL_422520 |
| EPI_ISL_423195 |
| EPI_ISL_416326 |
| EPI_ISL_425786 |
| EPI_ISL_416584 |
| EPI_ISL_423871 |
| EPI_ISL_414534 |
| EPI_ISL_427063 |
| EPI_ISL_423109 |
| EPI_ISL_420129 |

Table S3: GISAID sequence IDs collected on April 25, 2020.
| ID |
| --- |
| EPI_ISL_423484 |
| EPI_ISL_420775 |
| EPI_ISL_429121 |
| EPI_ISL_418337 |
| EPI_ISL_424406 |
| EPI_ISL_419823 |
| EPI_ISL_426624 |
| EPI_ISL_416693 |
| EPI_ISL_420267 |
| EPI_ISL_423507 |
| EPI_ISL_415148 |
| EPI_ISL_414456 |
| EPI_ISL_423406 |
| EPI_ISL_421924 |
| EPI_ISL_422940 |
| EPI_ISL_429119 |
| EPI_ISL_420020 |
| EPI_ISL_419533 |
| EPI_ISL_423584 |
| EPI_ISL_421198 |
| EPI_ISL_420622 |
| EPI_ISL_429397 |
| EPI_ISL_426982 |
| EPI_ISL_419835 |
| EPI_ISL_421984 |
| EPI_ISL_426116 |
| EPI_ISL_428478 |
| EPI_ISL_420393 |
| EPI_ISL_419751 |
| EPI_ISL_424579 |
| EPI_ISL_420053 |
| EPI_ISL_423314 |
| EPI_ISL_421828 |
| EPI_ISL_422504 |
| EPI_ISL_422893 |
| EPI_ISL_416690 |
| EPI_ISL_424963 |
| EPI_ISL_426113 |

Table S3: GISAID sequence IDs collected on April 25, 2020.
| ID |
| --- |
| EPI_ISL_421946 |
| EPI_ISL_427484 |
| EPI_ISL_416315 |
| EPI_ISL_422118 |
| EPI_ISL_423030 |
| EPI_ISL_426066 |
| EPI_ISL_421970 |
| EPI_ISL_426918 |
| EPI_ISL_423415 |
| EPI_ISL_421622 |
| EPI_ISL_427448 |
| EPI_ISL_429866 |
| EPI_ISL_426558 |
| EPI_ISL_421368 |
| EPI_ISL_418027 |
| EPI_ISL_416448 |
| EPI_ISL_417260 |
| EPI_ISL_427308 |
| EPI_ISL_421393 |
| EPI_ISL_420009 |
| EPI_ISL_423885 |
| EPI_ISL_420276 |
| EPI_ISL_428927 |
| EPI_ISL_425529 |
| EPI_ISL_420553 |
| EPI_ISL_418925 |
| EPI_ISL_423746 |
| EPI_ISL_422730 |
| EPI_ISL_426985 |
| EPI_ISL_425361 |
| EPI_ISL_427366 |
| EPI_ISL_418232 |
| EPI_ISL_425471 |
| EPI_ISL_421421 |
| EPI_ISL_428288 |
| EPI_ISL_421309 |
| EPI_ISL_416432 |
| EPI_ISL_416758 |

Table S3: GISAID sequence IDs collected on April 25, 2020.
| ID |
| --- |
| EPI_ISL_423904 |
| EPI_ISL_426694 |
| EPI_ISL_425241 |
| EPI_ISL_427068 |
| EPI_ISL_424308 |
| EPI_ISL_426506 |
| EPI_ISL_426068 |
| EPI_ISL_423764 |
| EPI_ISL_428352 |
| EPI_ISL_423360 |
| EPI_ISL_421303 |
| EPI_ISL_421390 |
| EPI_ISL_429629 |
| EPI_ISL_426929 |
| EPI_ISL_420214 |
| EPI_ISL_416585 |
| EPI_ISL_419233 |
| EPI_ISL_418981 |
| EPI_ISL_422157 |
| EPI_ISL_425138 |
| EPI_ISL_425301 |
| EPI_ISL_415515 |
| EPI_ISL_421350 |
| EPI_ISL_420630 |
| EPI_ISL_418212 |
| EPI_ISL_429800 |
| EPI_ISL_429290 |
| EPI_ISL_415613 |
| EPI_ISL_415658 |
| EPI_ISL_423497 |
| EPI_ISL_408977 |
| EPI_ISL_417630 |
| EPI_ISL_429624 |
| EPI_ISL_424143 |
| EPI_ISL_420485 |
| EPI_ISL_413020 |
| EPI_ISL_423165 |
| EPI_ISL_425708 |

Table S3: GISAID sequence IDs collected on April 25, 2020.
| ID |
| --- |
| EPI_ISL_420496 |
| EPI_ISL_427317 |
| EPI_ISL_423412 |
| EPI_ISL_423075 |
| EPI_ISL_424038 |
| EPI_ISL_416437 |
| EPI_ISL_417133 |
| EPI_ISL_429219 |
| EPI_ISL_429530 |
| EPI_ISL_429737 |
| EPI_ISL_426807 |
| EPI_ISL_425630 |
| EPI_ISL_423211 |
| EPI_ISL_422320 |
| EPI_ISL_424550 |
| EPI_ISL_426955 |
| EPI_ISL_413615 |
| EPI_ISL_423503 |
| EPI_ISL_417973 |
| EPI_ISL_419976 |
| EPI_ISL_421994 |
| EPI_ISL_425982 |
| EPI_ISL_429458 |
| EPI_ISL_425568 |
| EPI_ISL_424864 |
| EPI_ISL_418508 |
| EPI_ISL_417855 |
| EPI_ISL_423126 |
| EPI_ISL_420656 |
| EPI_ISL_423868 |
| EPI_ISL_425600 |
| EPI_ISL_429099 |
| EPI_ISL_418072 |
| EPI_ISL_429038 |
| EPI_ISL_417470 |
| EPI_ISL_423923 |
| EPI_ISL_425133 |
| EPI_ISL_426560 |

Table S3: GISAID sequence IDs collected on April 25, 2020.
| ID |
| --- |
| EPI_ISL_429517 |
| EPI_ISL_423879 |
| EPI_ISL_417099 |
| EPI_ISL_420685 |
| EPI_ISL_429599 |
| EPI_ISL_423397 |
| EPI_ISL_417876 |
| EPI_ISL_423667 |
| EPI_ISL_419892 |
| EPI_ISL_429080 |
| EPI_ISL_429079 |
| EPI_ISL_418956 |
| EPI_ISL_420086 |
| EPI_ISL_417472 |
| EPI_ISL_427351 |
| EPI_ISL_419593 |
| EPI_ISL_420935 |
| EPI_ISL_428836 |
| EPI_ISL_417544 |
| EPI_ISL_416594 |
| EPI_ISL_417405 |
| EPI_ISL_425862 |
| EPI_ISL_421829 |
| EPI_ISL_425712 |
| EPI_ISL_427103 |
| EPI_ISL_423716 |
| EPI_ISL_414007 |
| EPI_ISL_419558 |
| EPI_ISL_428960 |
| EPI_ISL_416517 |
| EPI_ISL_423429 |
| EPI_ISL_422697 |
| EPI_ISL_420969 |
| EPI_ISL_426734 |
| EPI_ISL_430093 |
| EPI_ISL_426035 |
| EPI_ISL_420512 |
| EPI_ISL_419587 |

Table S3: GISAID sequence IDs collected on April 25, 2020.
| ID |
| --- |
| EPI_ISL_427204 |
| EPI_ISL_417359 |
| EPI_ISL_416581 |
| EPI_ISL_417796 |
| EPI_ISL_415142 |
| EPI_ISL_419684 |
| EPI_ISL_419741 |
| EPI_ISL_416596 |
| EPI_ISL_425401 |
| EPI_ISL_427699 |
| EPI_ISL_422167 |
| EPI_ISL_429299 |
| EPI_ISL_415707 |
| EPI_ISL_423336 |
| EPI_ISL_428670 |
| EPI_ISL_427500 |
| EPI_ISL_425177 |
| EPI_ISL_422603 |
| EPI_ISL_424509 |
| EPI_ISL_419993 |
| EPI_ISL_428709 |
| EPI_ISL_428284 |
| EPI_ISL_421902 |
| EPI_ISL_419943 |
| EPI_ISL_421244 |
| EPI_ISL_419927 |
| EPI_ISL_421720 |
| EPI_ISL_419223 |
| EPI_ISL_418963 |
| EPI_ISL_426289 |
| EPI_ISL_426724 |
| EPI_ISL_424146 |
| EPI_ISL_421776 |
| EPI_ISL_425182 |
| EPI_ISL_414517 |
| EPI_ISL_416443 |
| EPI_ISL_414541 |
| EPI_ISL_422689 |

Table S3: GISAID sequence IDs collected on April 25, 2020.
| ID |
| --- |
| EPI_ISL_417742 |
| EPI_ISL_418015 |
| EPI_ISL_421996 |
| EPI_ISL_414643 |
| EPI_ISL_428317 |
| EPI_ISL_422345 |
| EPI_ISL_420521 |
| EPI_ISL_420318 |
| EPI_ISL_424431 |
| EPI_ISL_422265 |
| EPI_ISL_418293 |
| EPI_ISL_422432 |
| EPI_ISL_424197 |
| EPI_ISL_425178 |
| EPI_ISL_420940 |
| EPI_ISL_417432 |
| EPI_ISL_425692 |
| EPI_ISL_417444 |
| EPI_ISL_429821 |
| EPI_ISL_420226 |
| EPI_ISL_414542 |
| EPI_ISL_418025 |
| EPI_ISL_429010 |
| EPI_ISL_425122 |
| EPI_ISL_421301 |
| EPI_ISL_423957 |
| EPI_ISL_423949 |
| EPI_ISL_428870 |
| EPI_ISL_420154 |
| EPI_ISL_422941 |
| EPI_ISL_423246 |
| EPI_ISL_421876 |
| EPI_ISL_430100 |
| EPI_ISL_426075 |
| EPI_ISL_417215 |
| EPI_ISL_419228 |
| EPI_ISL_418777 |
| EPI_ISL_418953 |

Table S3: GISAID sequence IDs collected on April 25, 2020.
| ID |
| --- |
| EPI_ISL_418920 |
| EPI_ISL_430020 |
| EPI_ISL_424865 |
| EPI_ISL_423054 |
| EPI_ISL_423175 |
| EPI_ISL_427123 |
| EPI_ISL_425953 |
| EPI_ISL_416499 |
| EPI_ISL_429209 |
| EPI_ISL_427081 |
| EPI_ISL_430001 |
| EPI_ISL_423307 |
| EPI_ISL_417204 |
| EPI_ISL_427629 |
| EPI_ISL_420250 |
| EPI_ISL_423010 |
| EPI_ISL_429603 |
| EPI_ISL_424643 |
| EPI_ISL_408479 |
| EPI_ISL_425941 |
| EPI_ISL_423189 |
| EPI_ISL_425449 |
| EPI_ISL_424874 |
| EPI_ISL_416753 |
| EPI_ISL_418994 |
| EPI_ISL_425944 |
| EPI_ISL_423586 |
| EPI_ISL_413589 |
| EPI_ISL_417575 |
| EPI_ISL_430028 |
| EPI_ISL_423803 |
| EPI_ISL_427129 |
| EPI_ISL_418753 |
| EPI_ISL_429853 |
| EPI_ISL_428395 |
| EPI_ISL_417564 |
| EPI_ISL_420985 |
| EPI_ISL_420382 |

Table S3: GISAID sequence IDs collected on April 25, 2020.
| ID |
| --- |
| EPI_ISL_425691 |
| EPI_ISL_427700 |
| EPI_ISL_427065 |
| EPI_ISL_427304 |
| EPI_ISL_416607 |
| EPI_ISL_420321 |
| EPI_ISL_426064 |
| EPI_ISL_421395 |
| EPI_ISL_422253 |
| EPI_ISL_427452 |
| EPI_ISL_423494 |
| EPI_ISL_422422 |
| EPI_ISL_417569 |
| EPI_ISL_418044 |
| EPI_ISL_424128 |
| EPI_ISL_421382 |
| EPI_ISL_422542 |
| EPI_ISL_424176 |
| EPI_ISL_426499 |
| EPI_ISL_424466 |
| EPI_ISL_425358 |
| EPI_ISL_420195 |
| EPI_ISL_424895 |
| EPI_ISL_424414 |
| EPI_ISL_413569 |
| EPI_ISL_424506 |
| EPI_ISL_427566 |
| EPI_ISL_422529 |
| EPI_ISL_403935 |
| EPI_ISL_423637 |
| EPI_ISL_420007 |
| EPI_ISL_412981 |
| EPI_ISL_424397 |
| EPI_ISL_426550 |
| EPI_ISL_417024 |
| EPI_ISL_427385 |
| EPI_ISL_422054 |
| EPI_ISL_415461 |

Table S3: GISAID sequence IDs collected on April 25, 2020.
| ID |
| --- |
| EPI_ISL_426937 |
| EPI_ISL_426519 |
| EPI_ISL_427252 |
| EPI_ISL_428334 |
| EPI_ISL_418695 |
| EPI_ISL_428794 |
| EPI_ISL_429183 |
| EPI_ISL_424384 |
| EPI_ISL_427069 |
| EPI_ISL_425167 |
| EPI_ISL_419724 |
| EPI_ISL_416331 |
| EPI_ISL_423961 |
| EPI_ISL_427187 |
| EPI_ISL_424529 |
| EPI_ISL_424440 |
| EPI_ISL_419800 |
| EPI_ISL_425775 |
| EPI_ISL_425992 |
| EPI_ISL_420016 |
| EPI_ISL_416691 |
| EPI_ISL_427175 |
| EPI_ISL_426651 |
| EPI_ISL_428725 |
| EPI_ISL_418351 |
| EPI_ISL_423006 |
| EPI_ISL_423294 |
| EPI_ISL_421507 |
| EPI_ISL_424451 |
| EPI_ISL_429567 |
| EPI_ISL_426956 |
| EPI_ISL_428919 |
| EPI_ISL_419400 |
| EPI_ISL_420993 |
| EPI_ISL_421398 |
| EPI_ISL_429269 |
| EPI_ISL_419663 |
| EPI_ISL_417368 |

Table S3: GISAID sequence IDs collected on April 25, 2020.
| ID |
| --- |
| EPI_ISL_428697 |
| EPI_ISL_427191 |
| EPI_ISL_423376 |
| EPI_ISL_428384 |
| EPI_ISL_425789 |
| EPI_ISL_417988 |
| EPI_ISL_422570 |
| EPI_ISL_427022 |
| EPI_ISL_422820 |
| EPI_ISL_414689 |
| EPI_ISL_416033 |
| EPI_ISL_417734 |
| EPI_ISL_425523 |

**Table S4:**
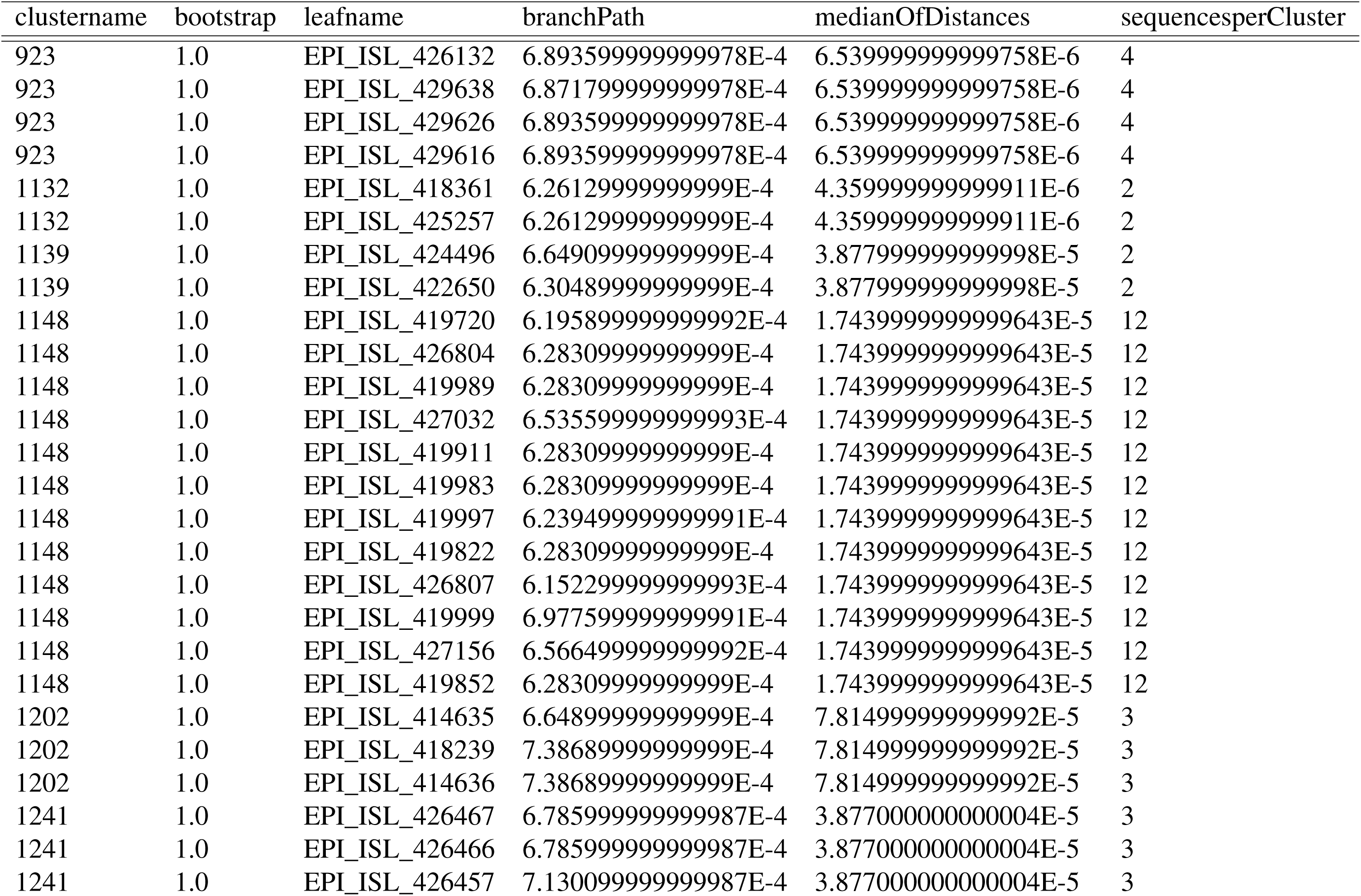
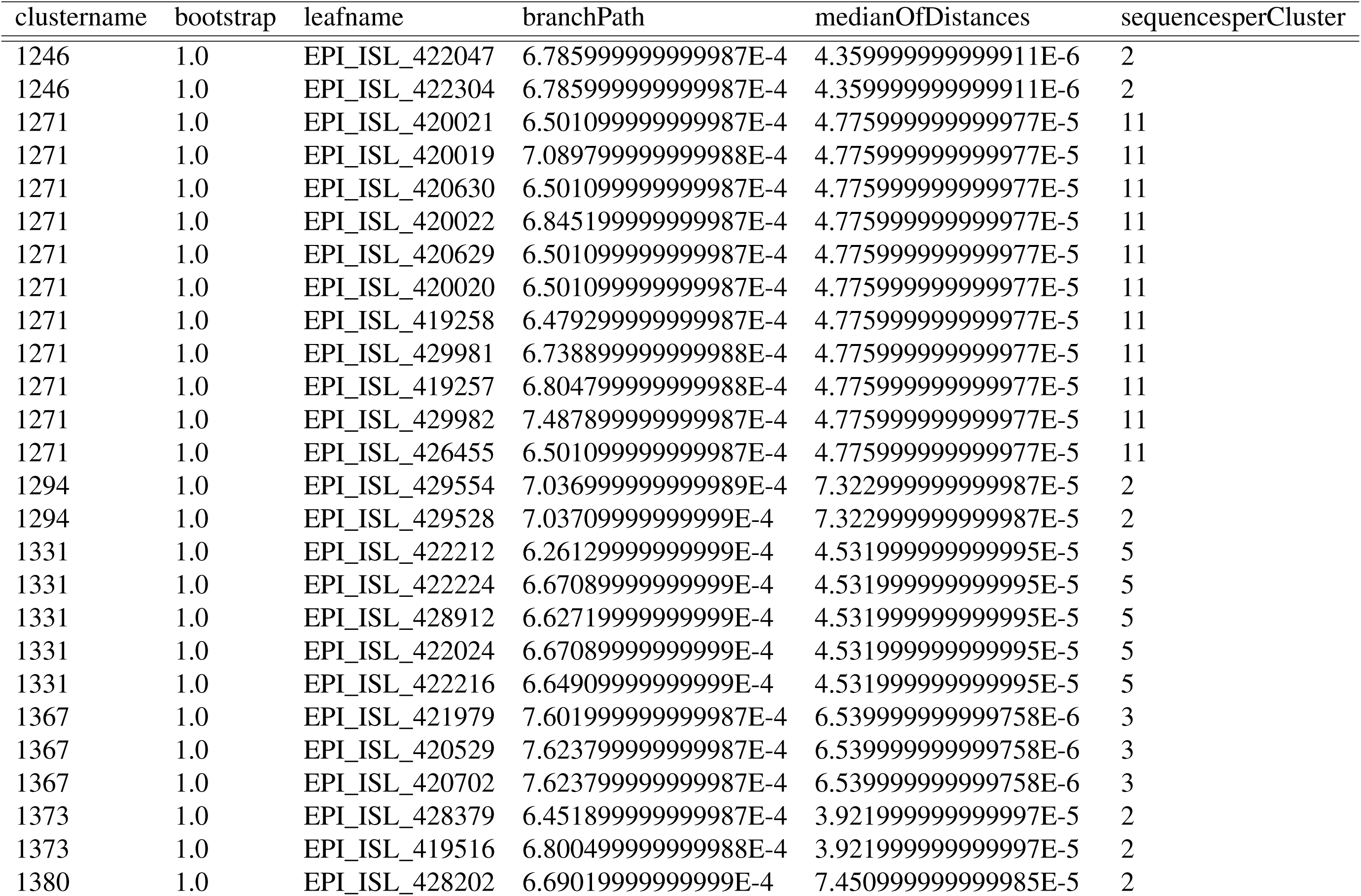
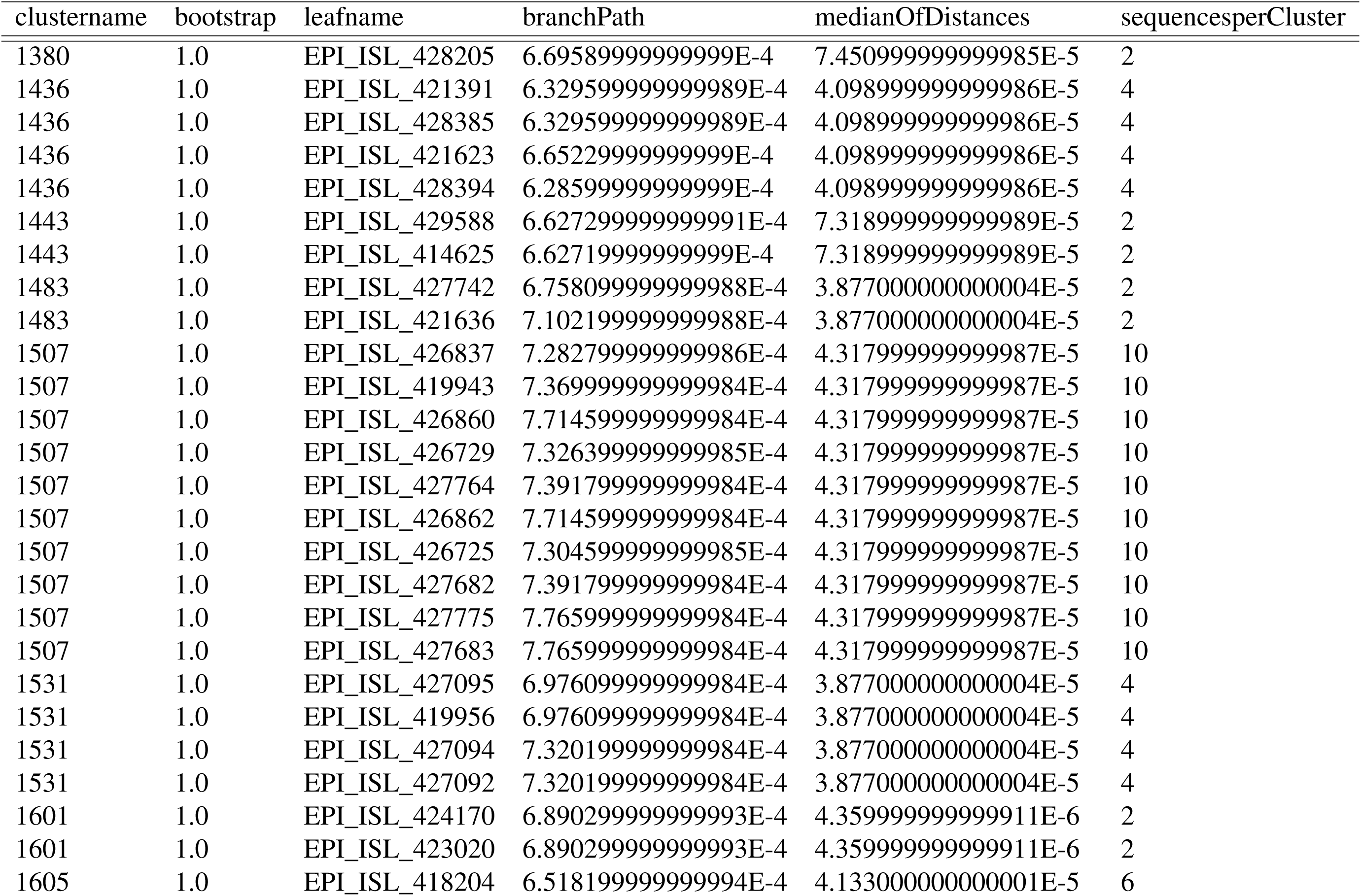
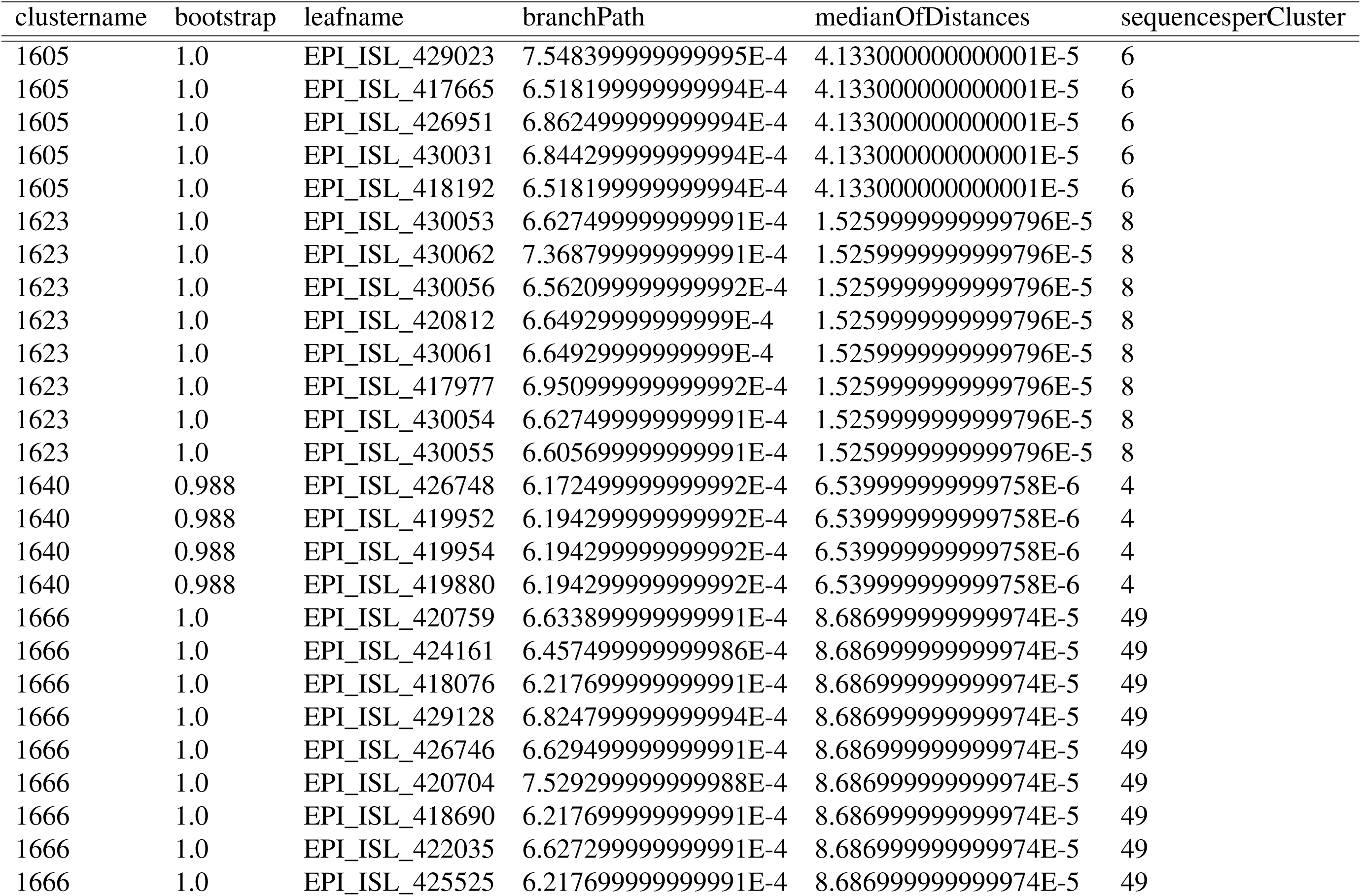
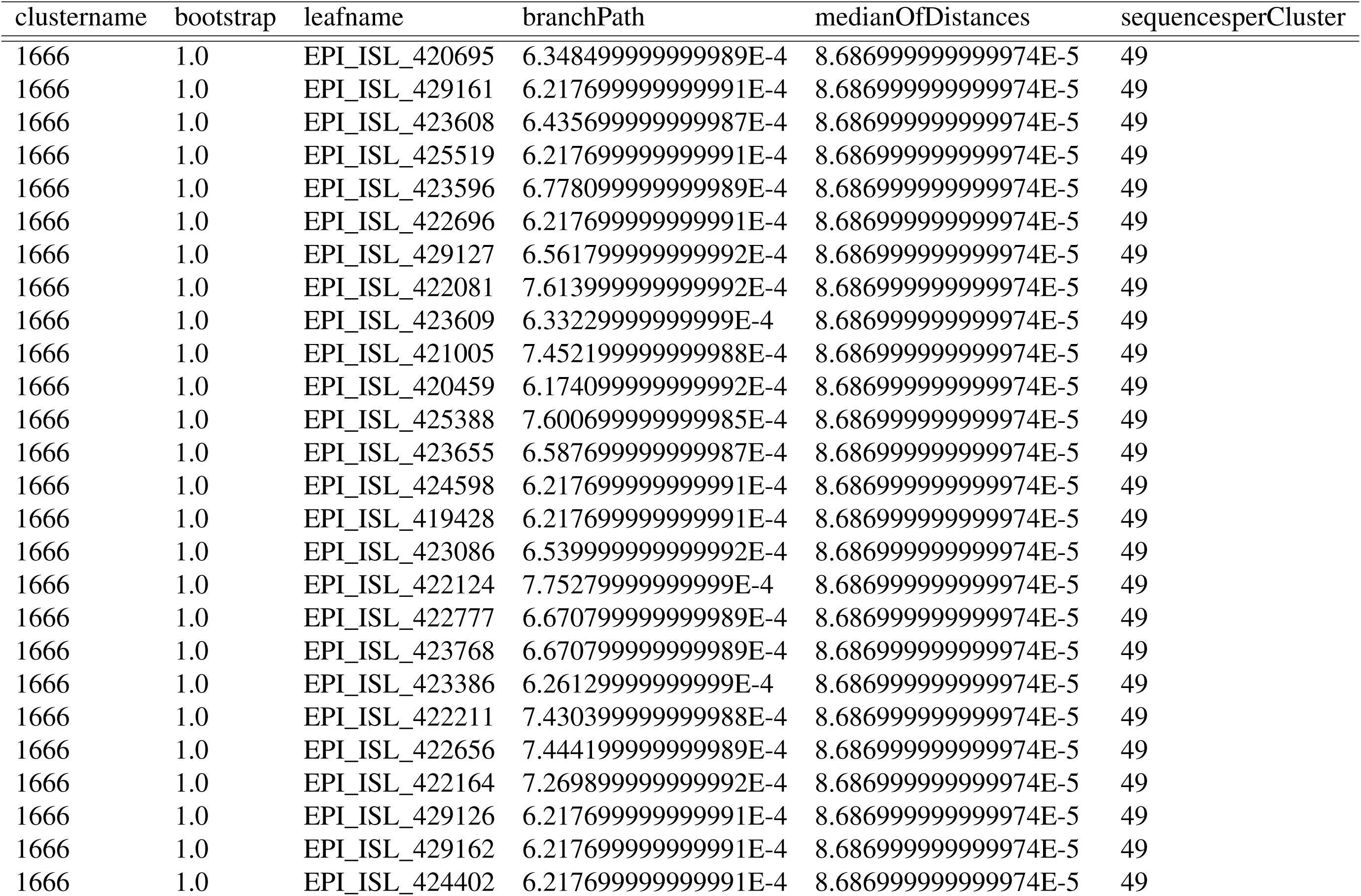
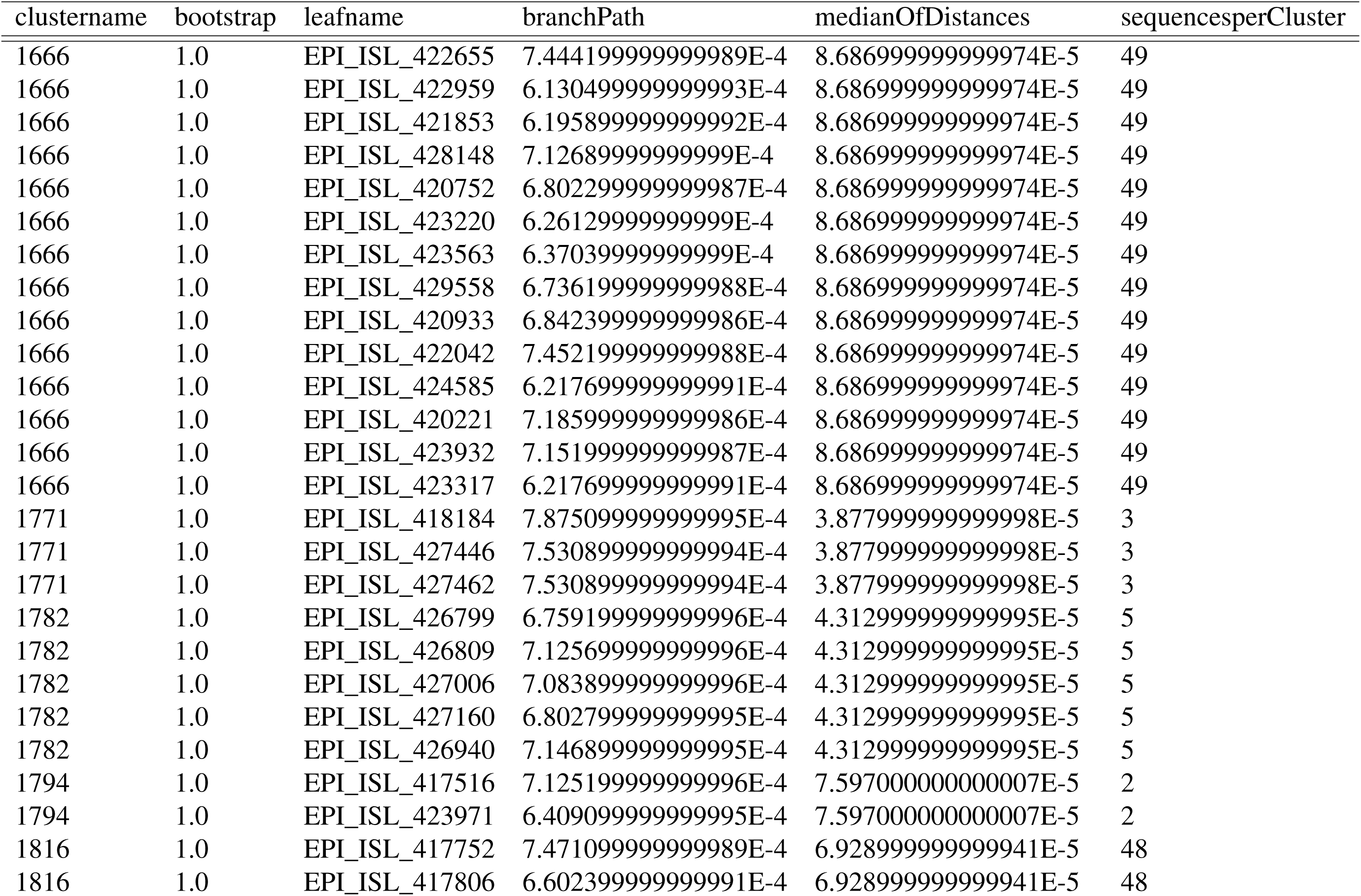
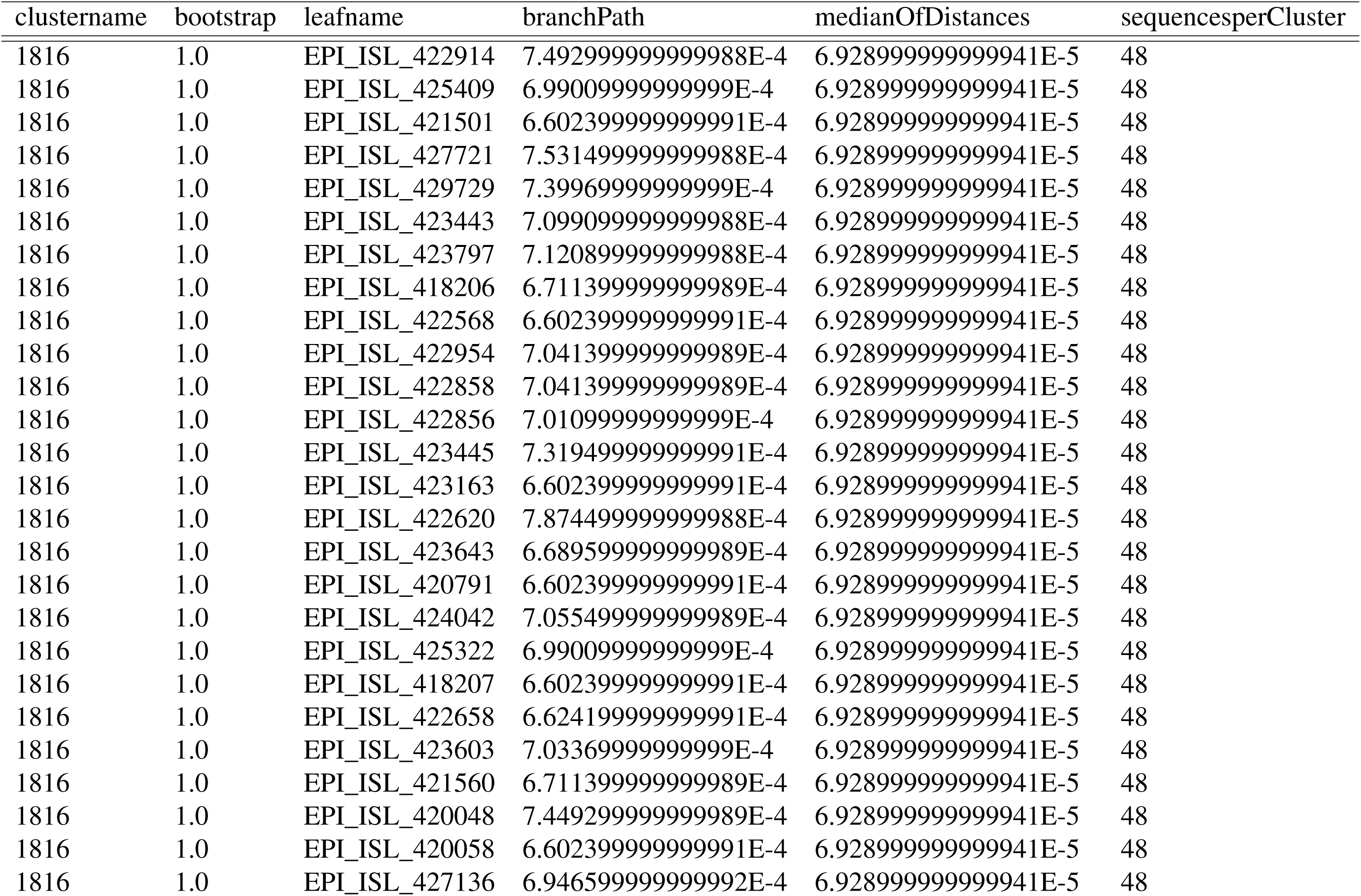
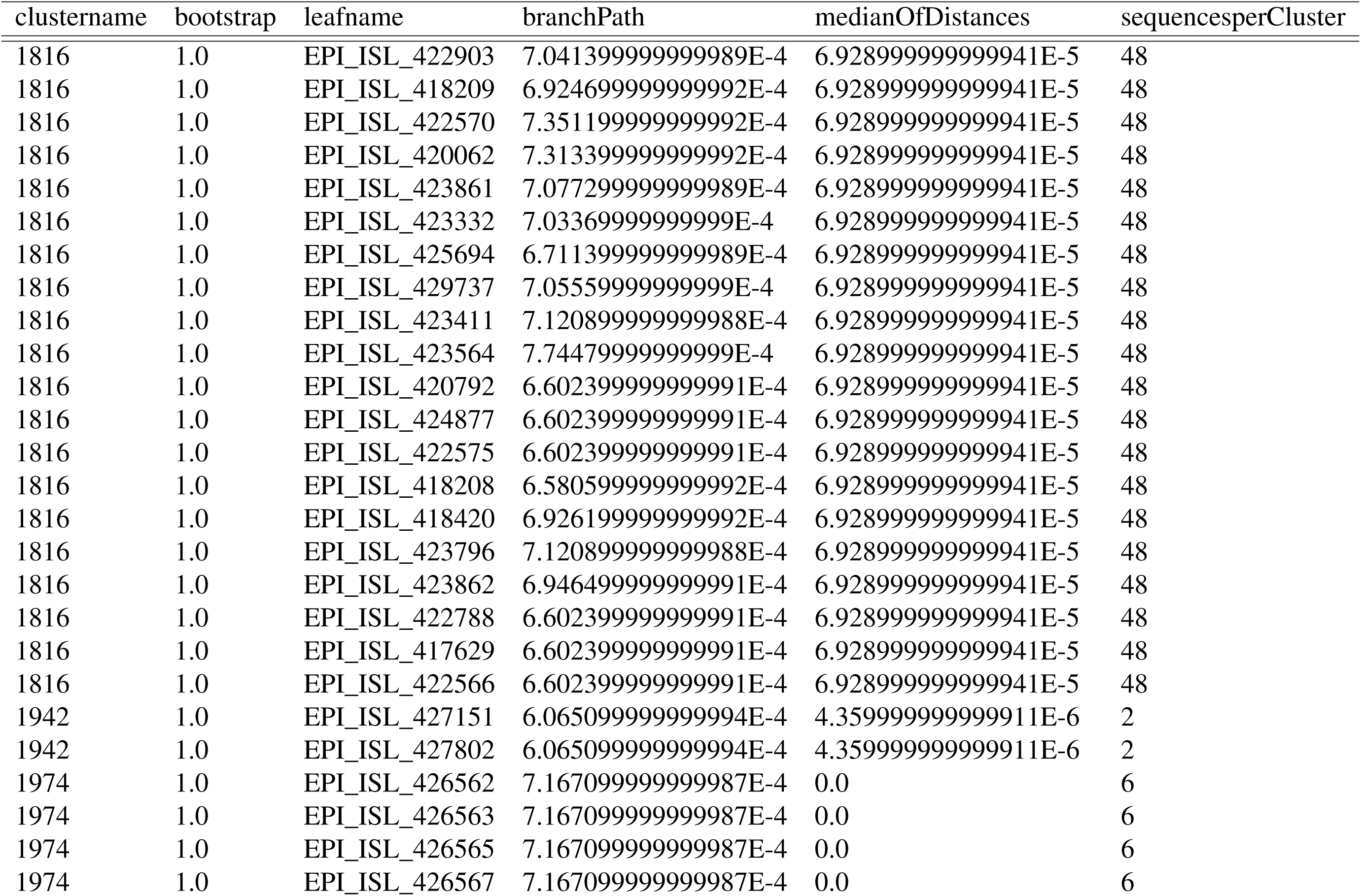
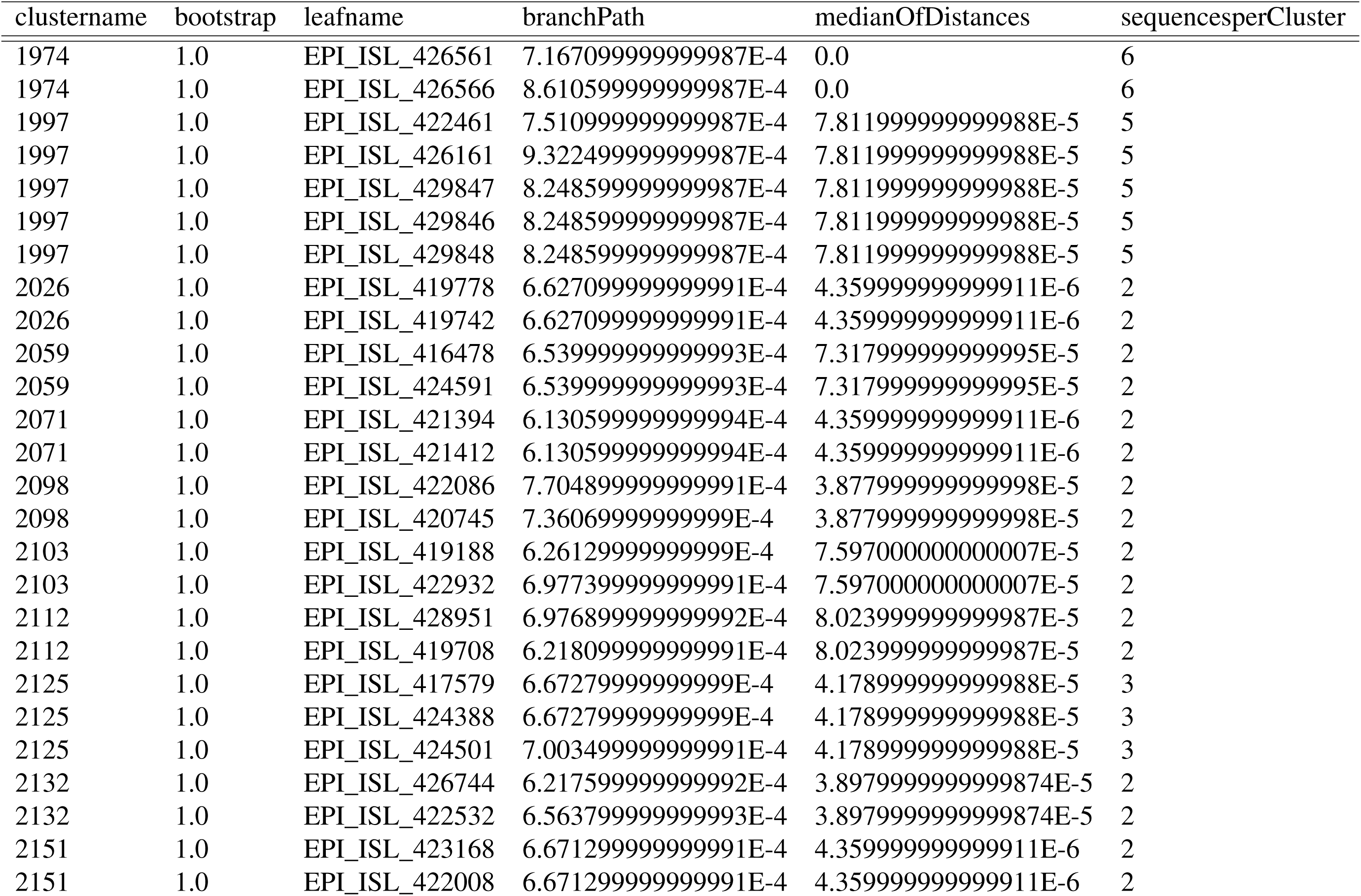
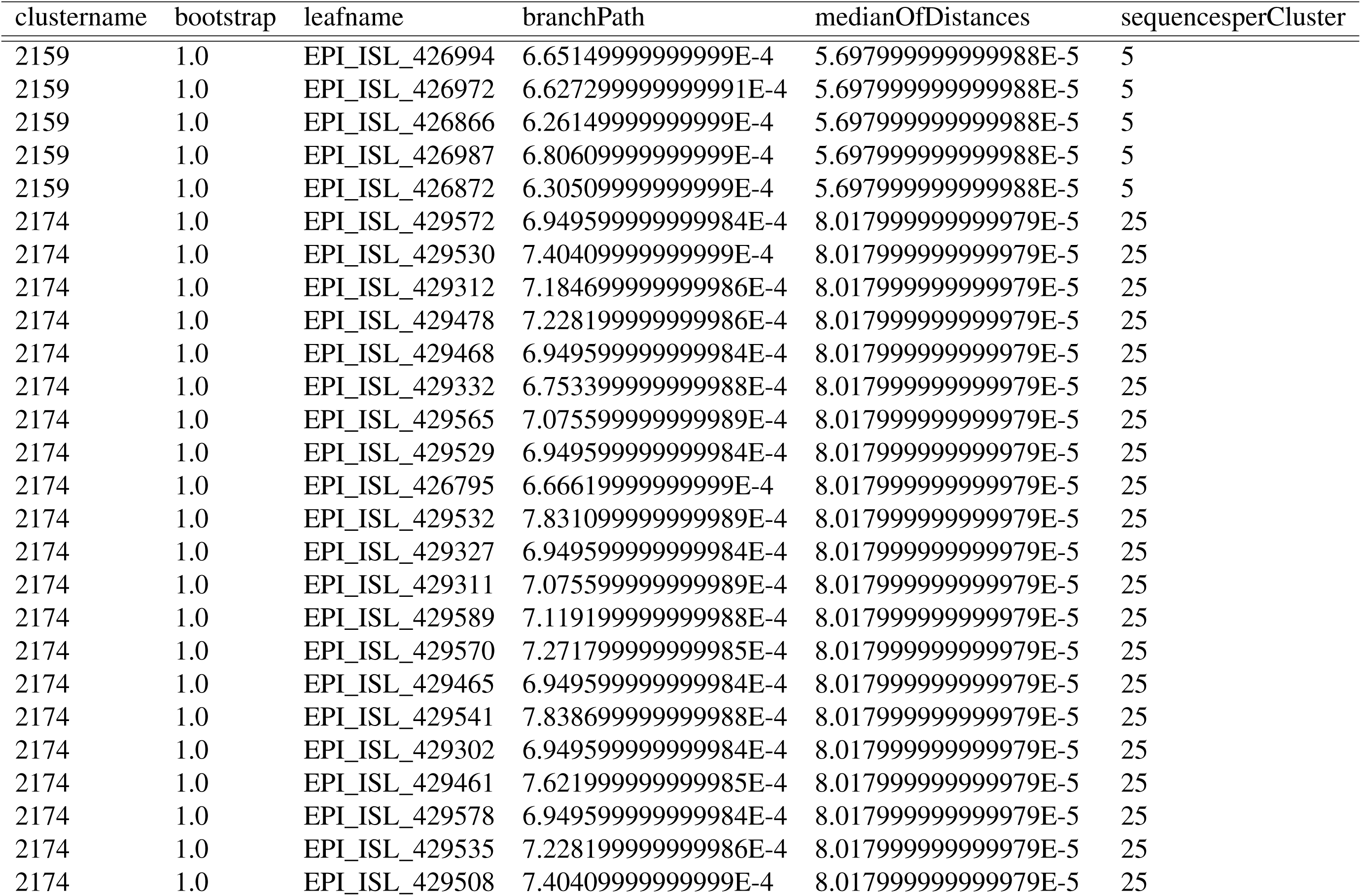
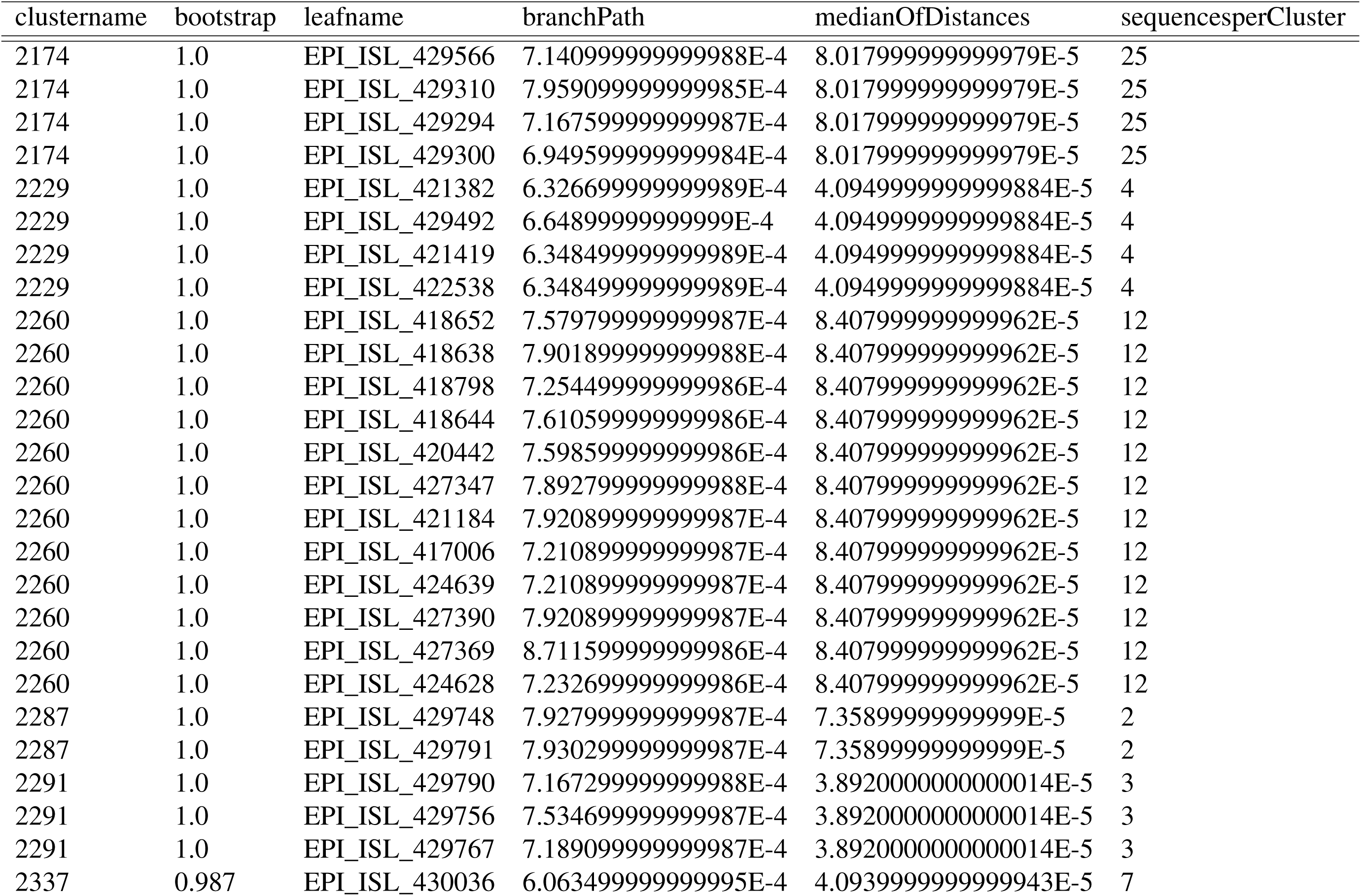
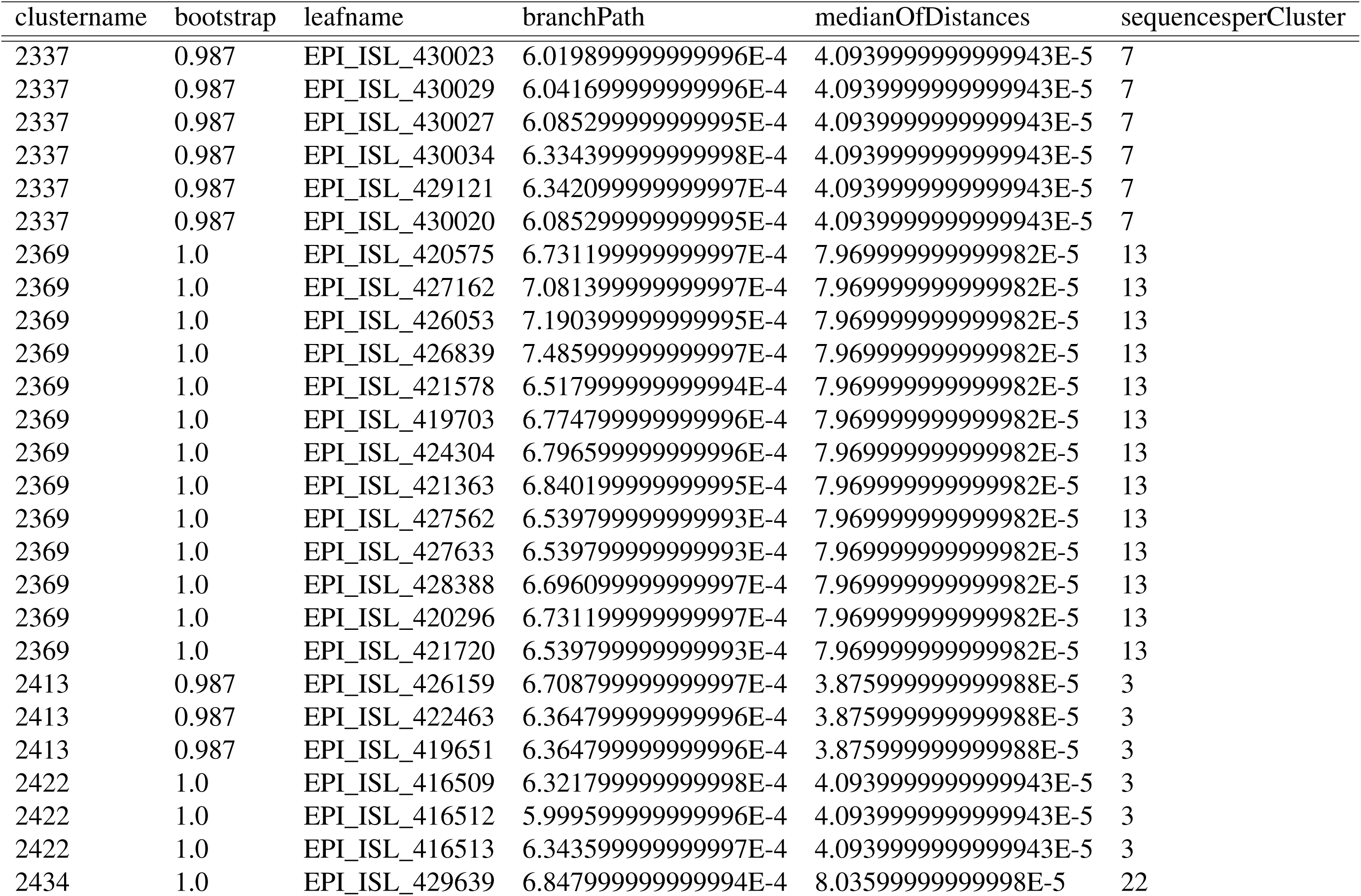
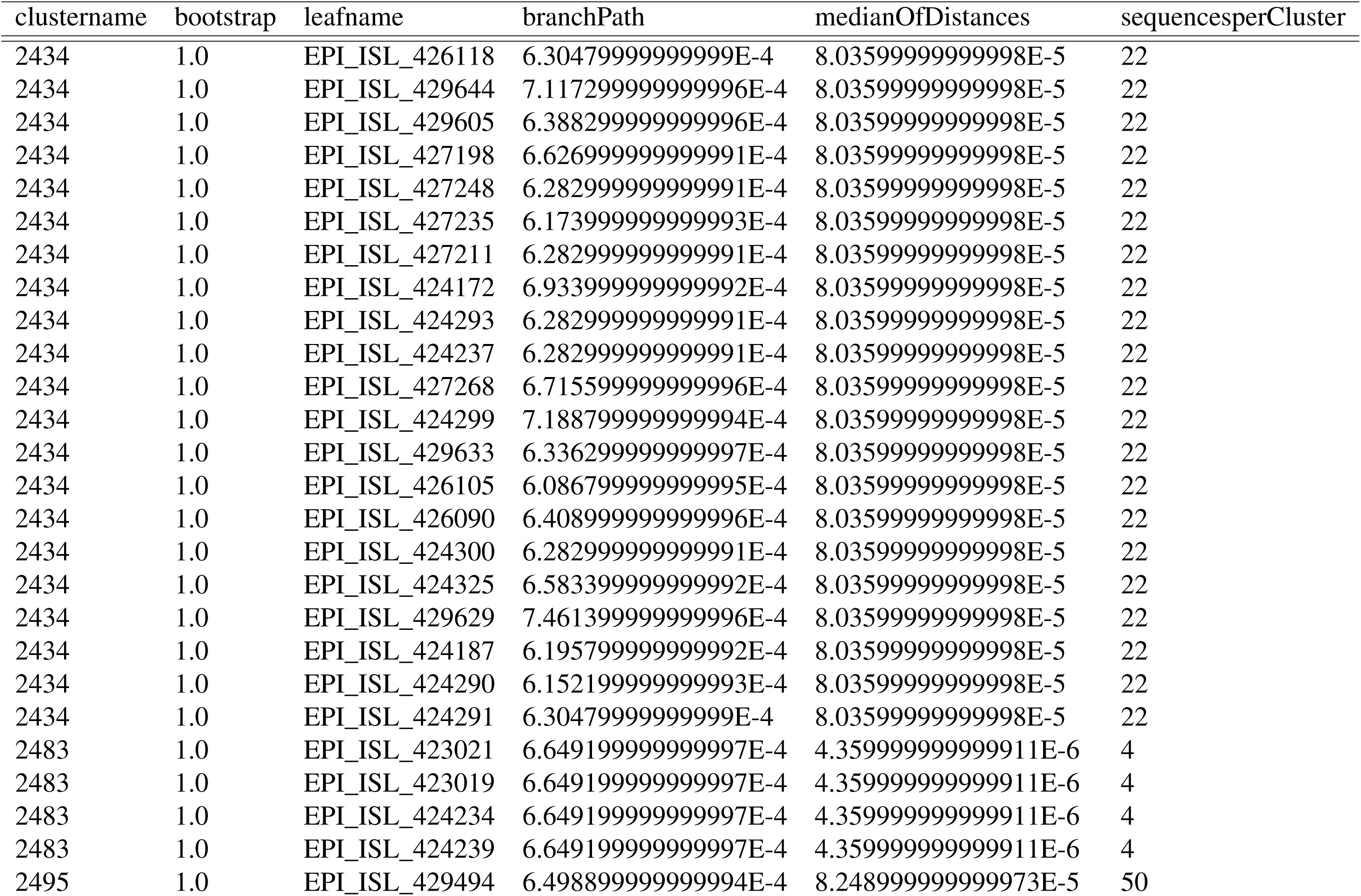
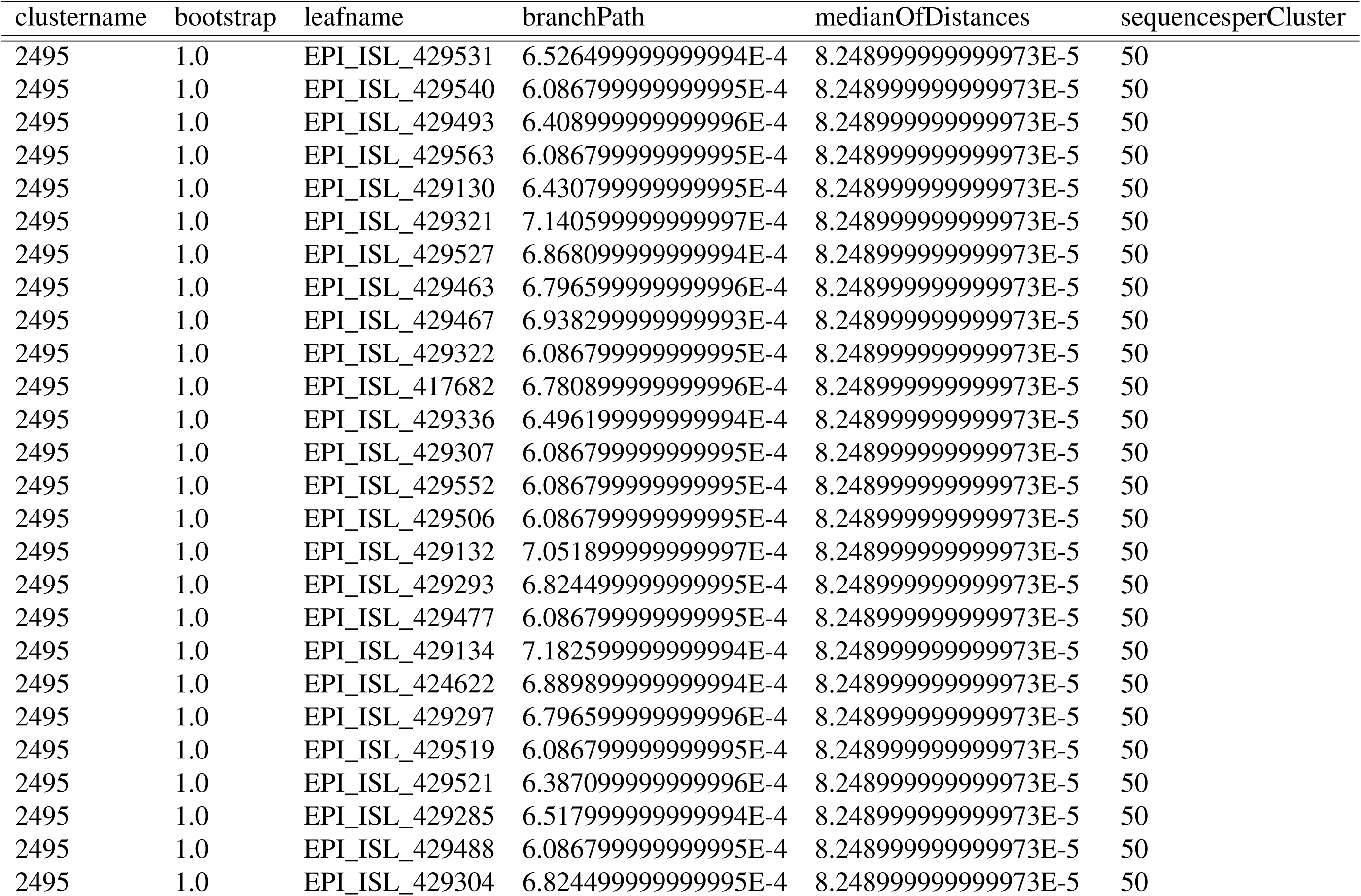
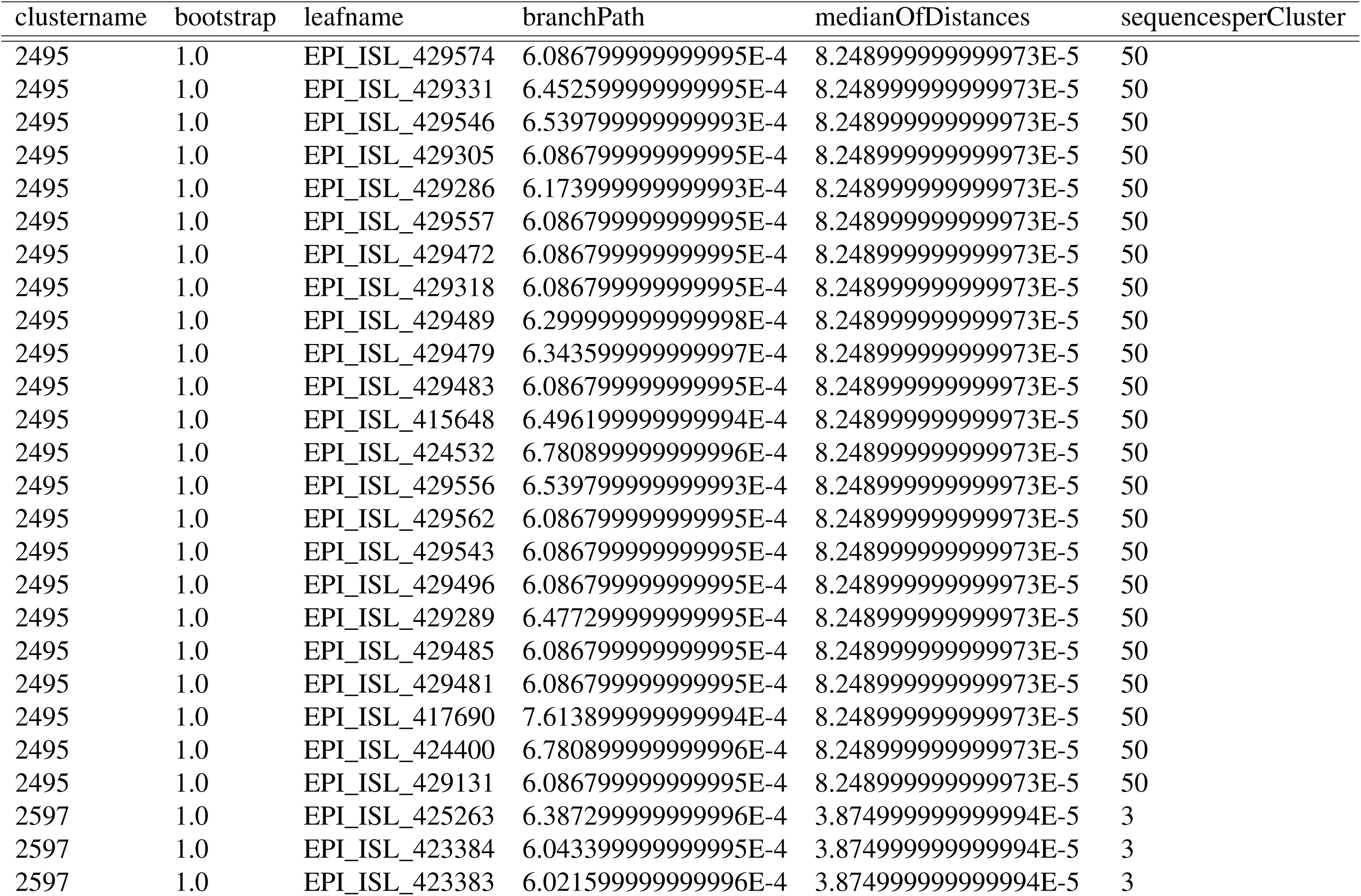
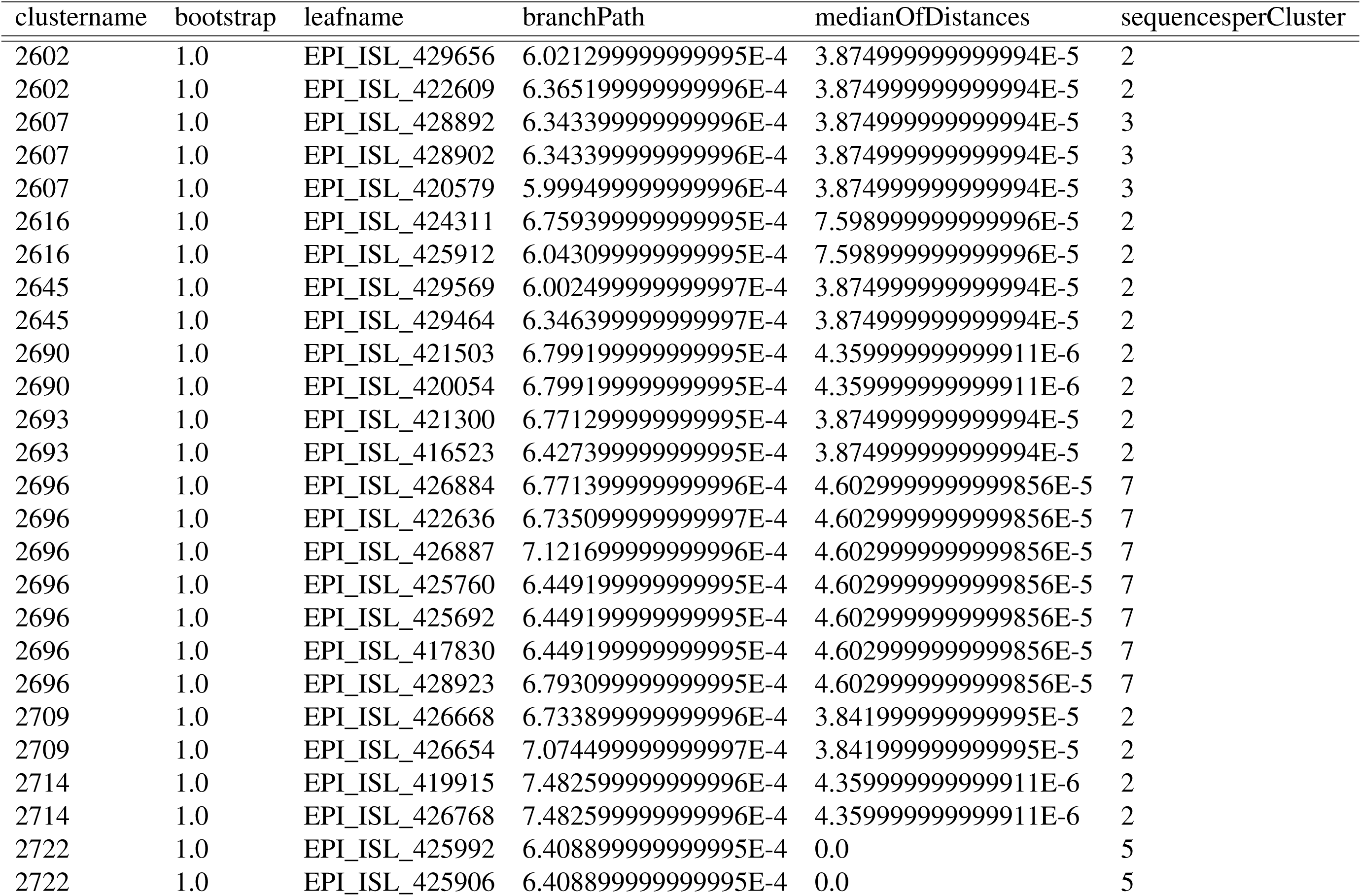
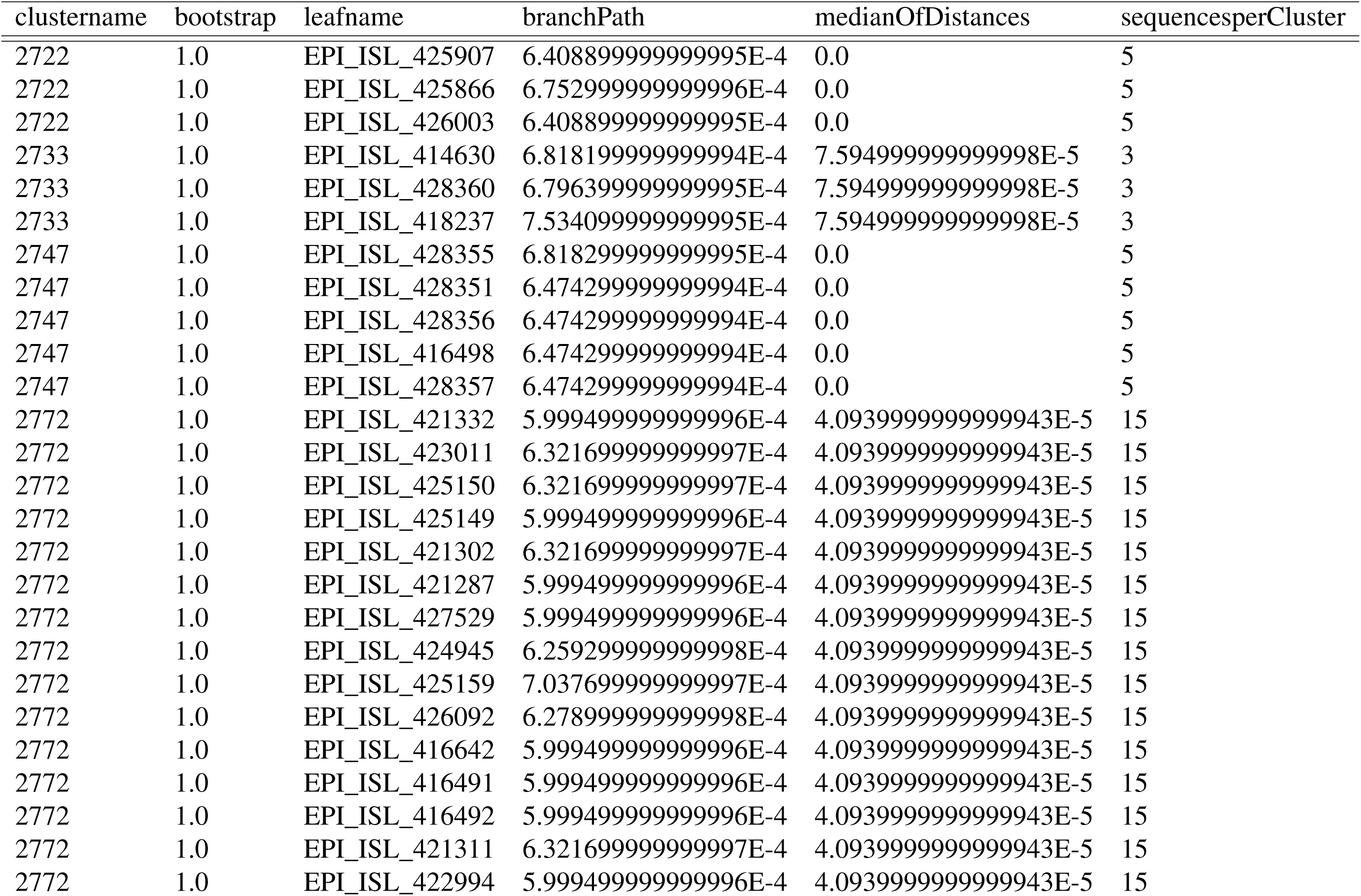
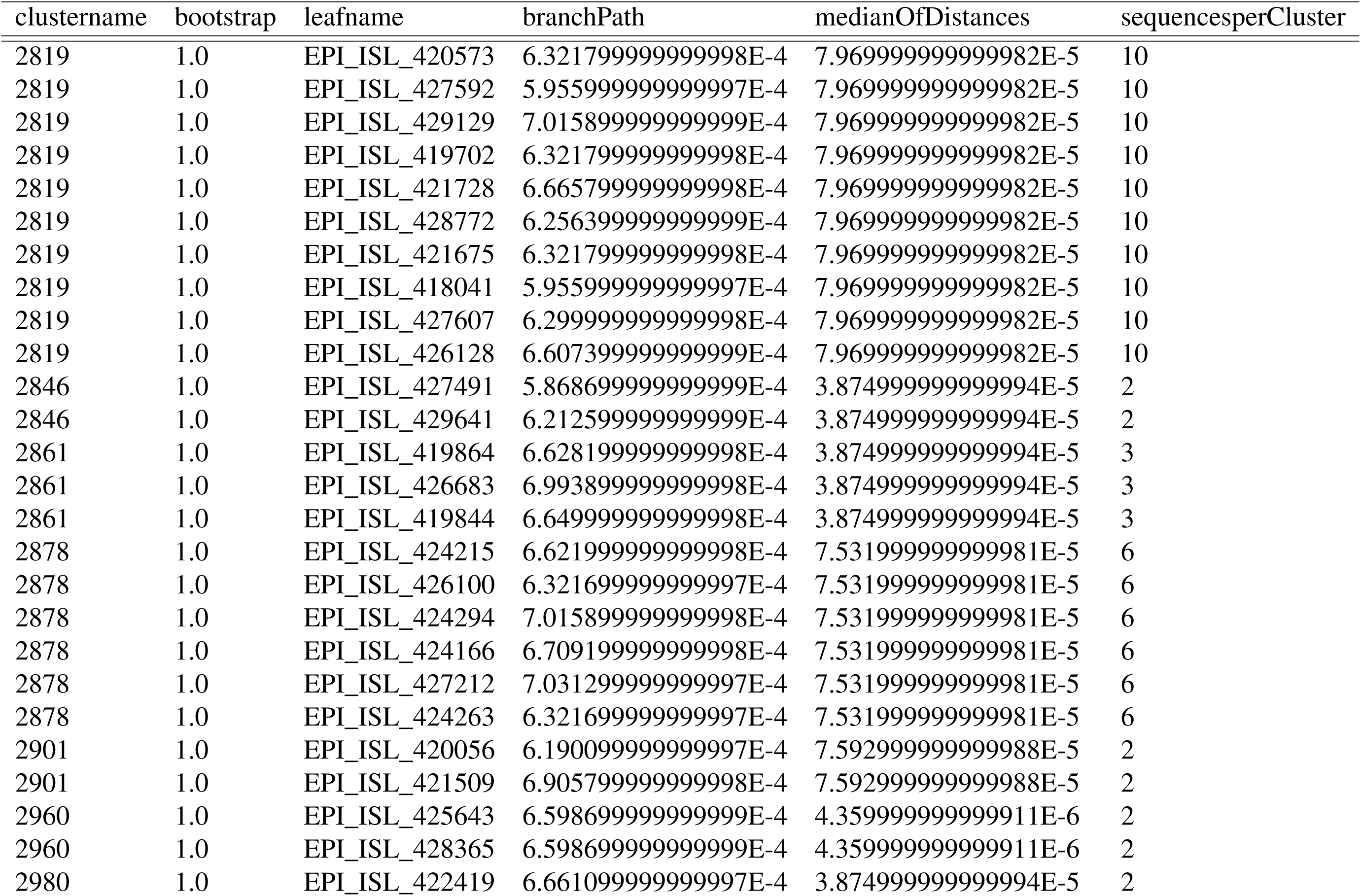
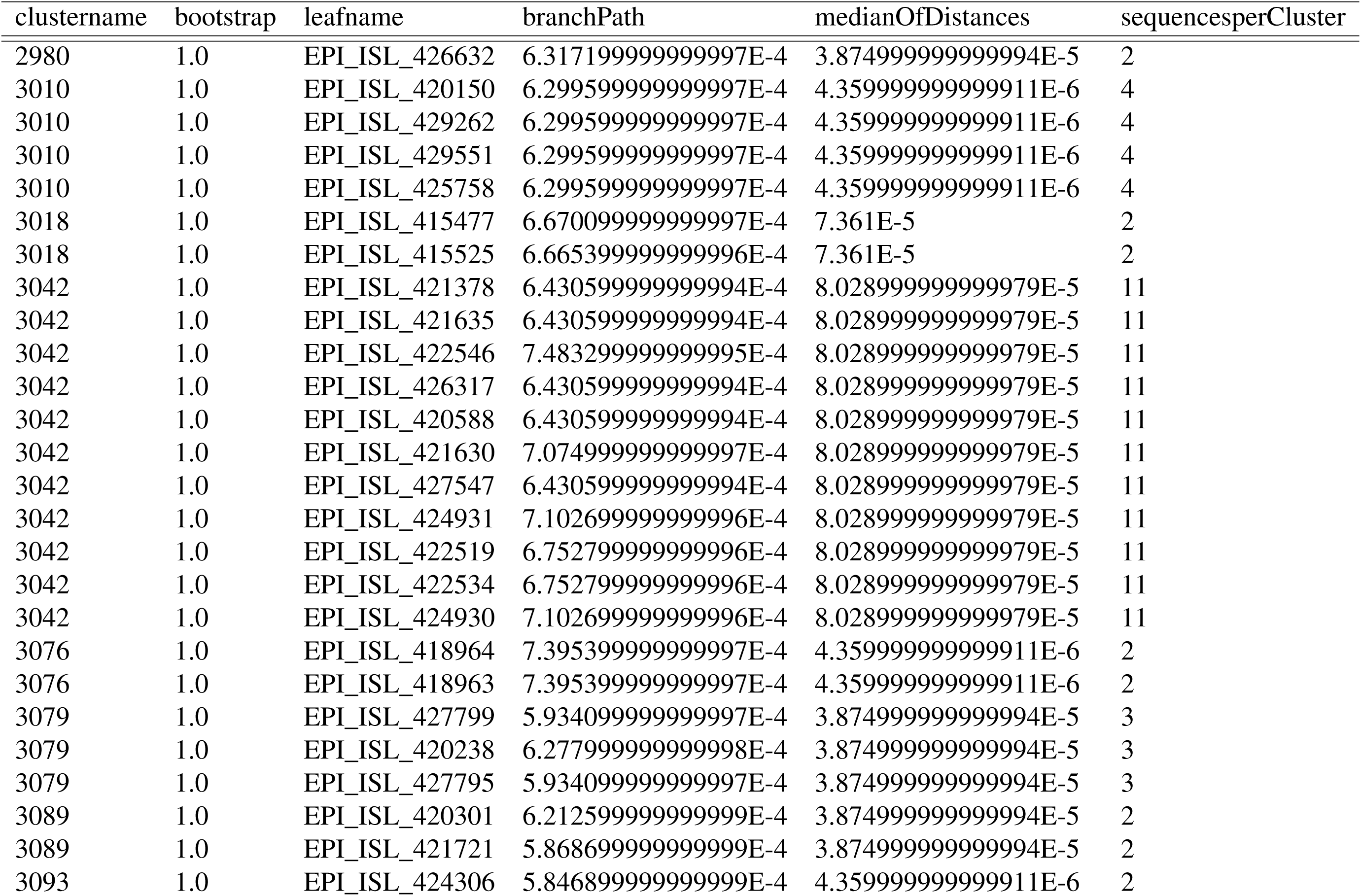
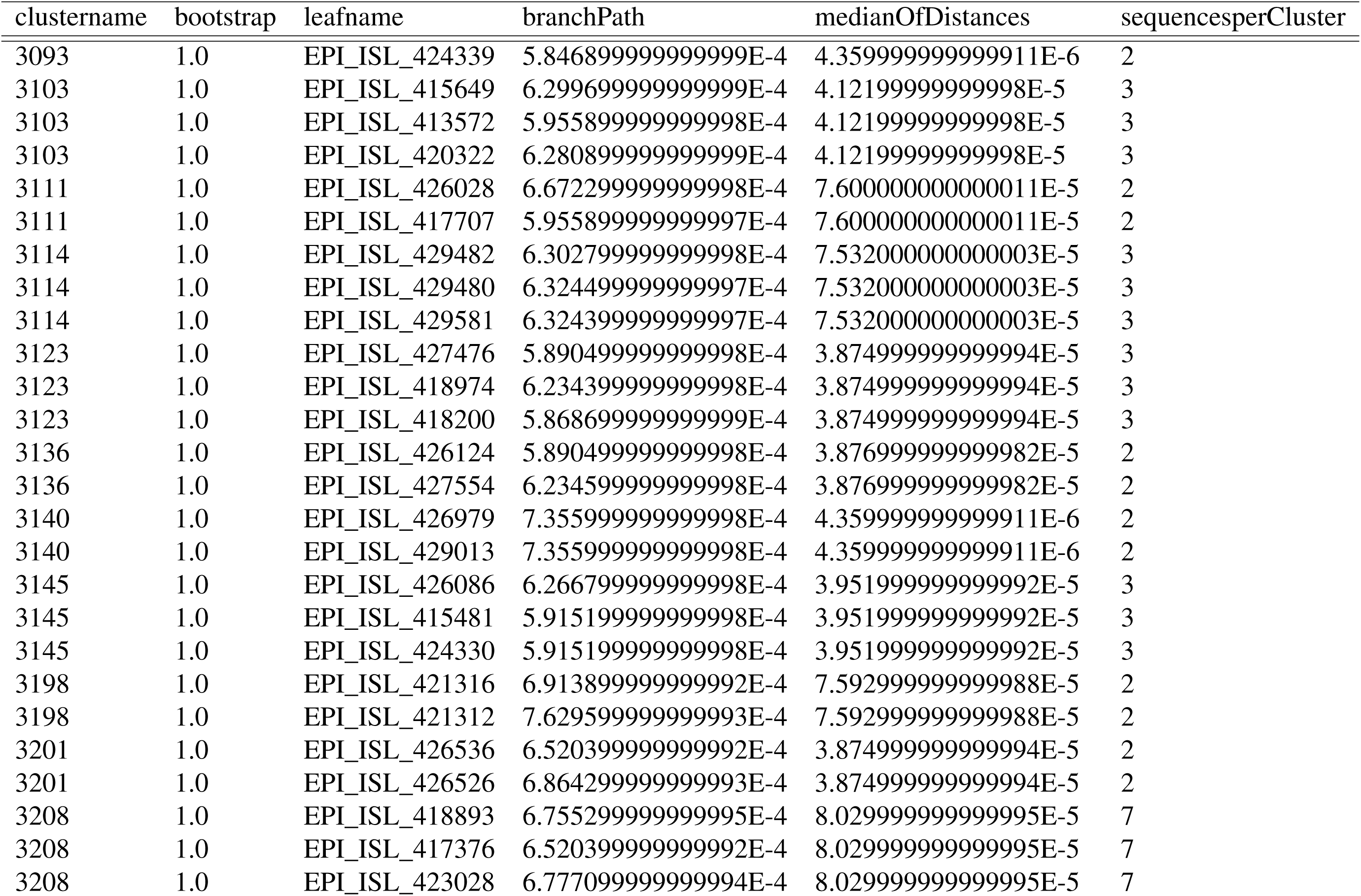
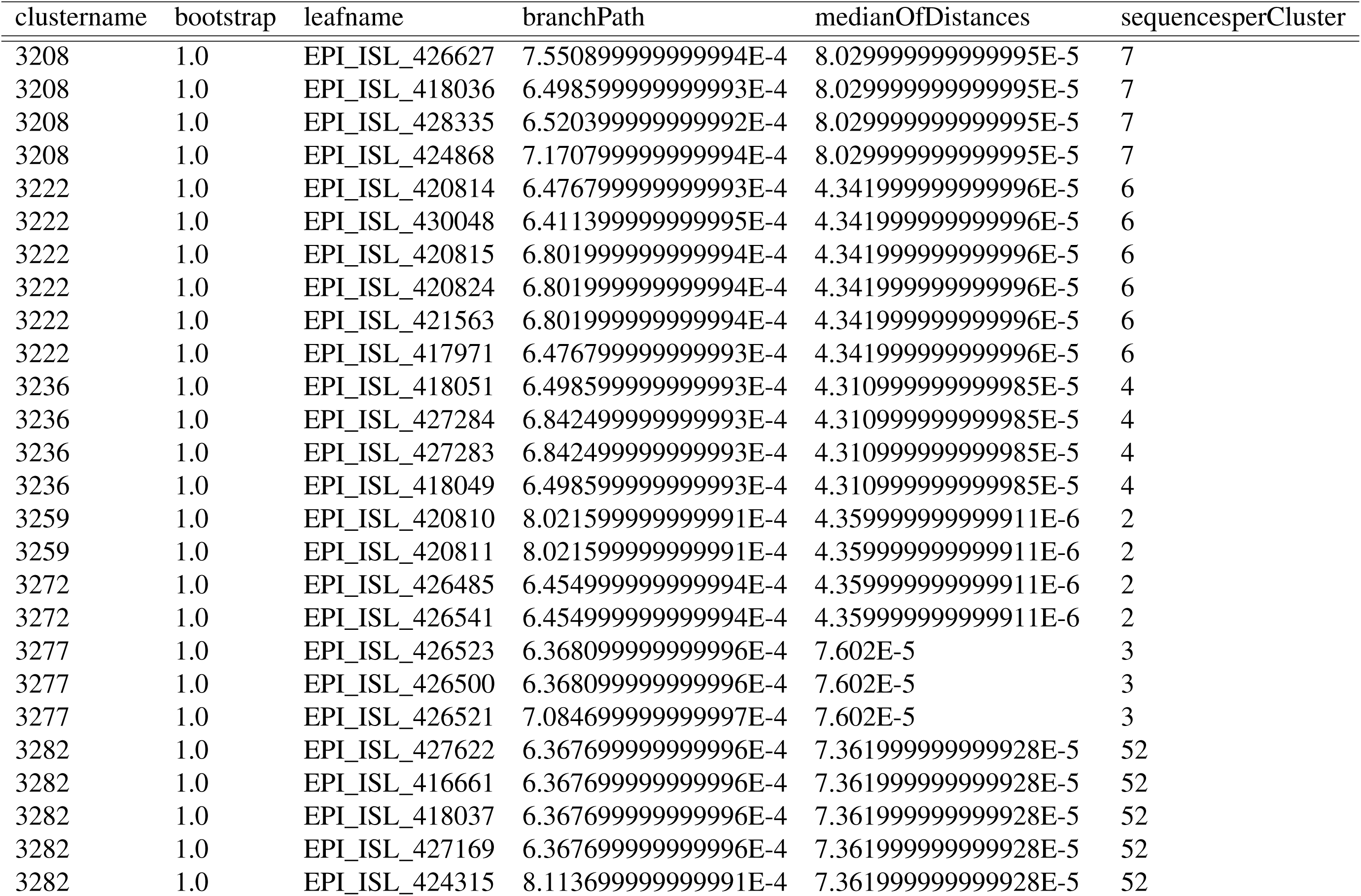
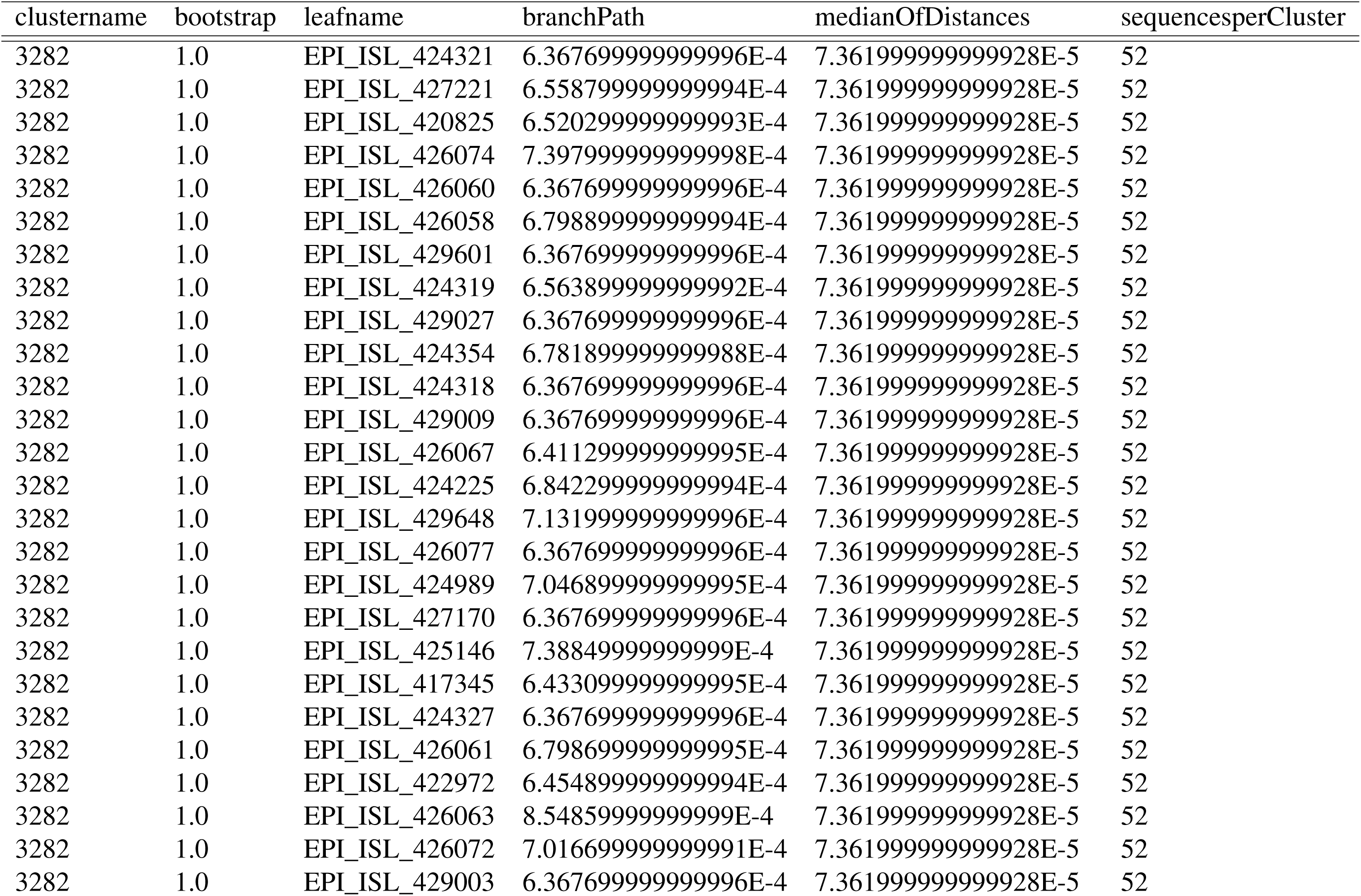
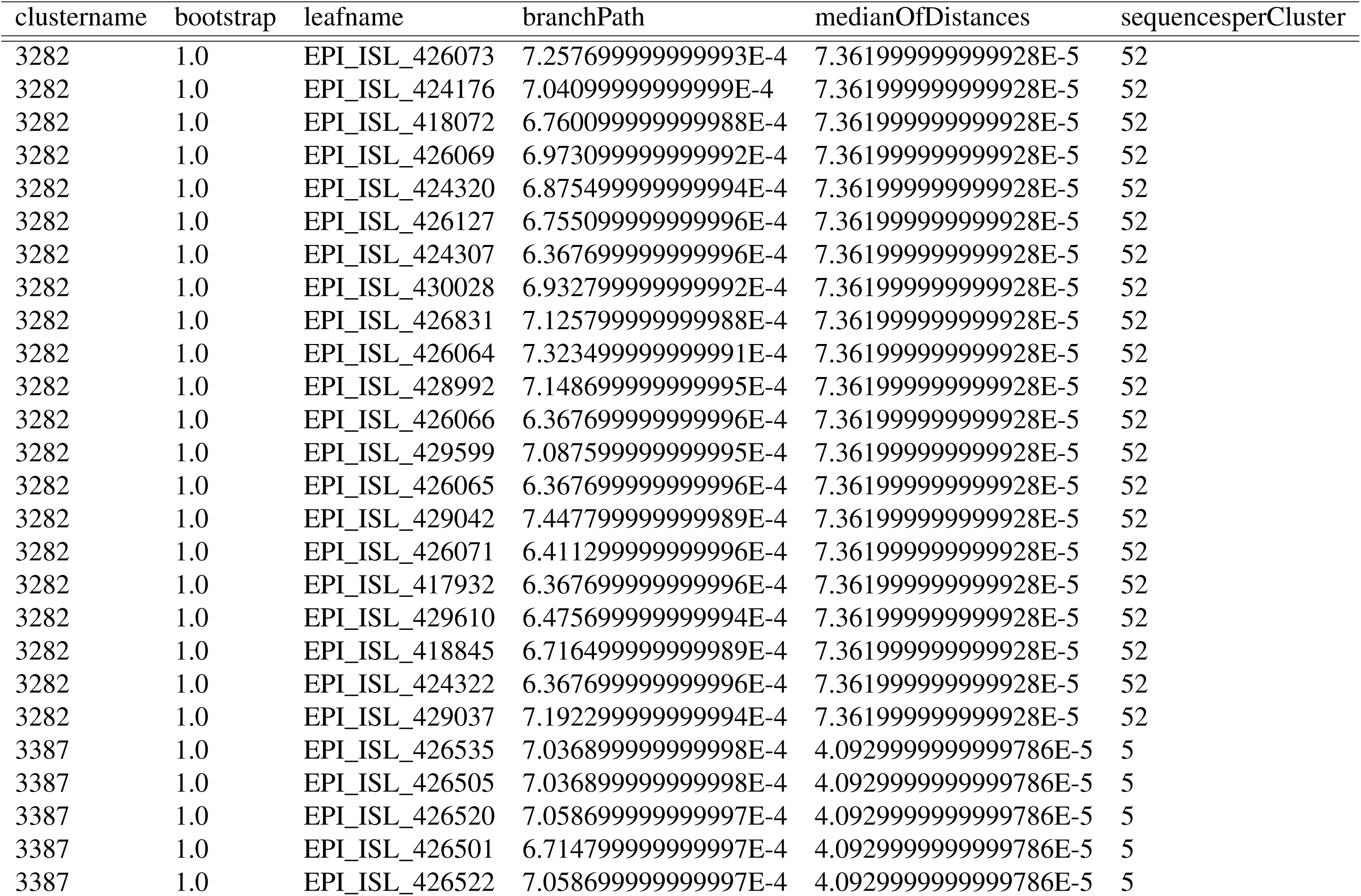
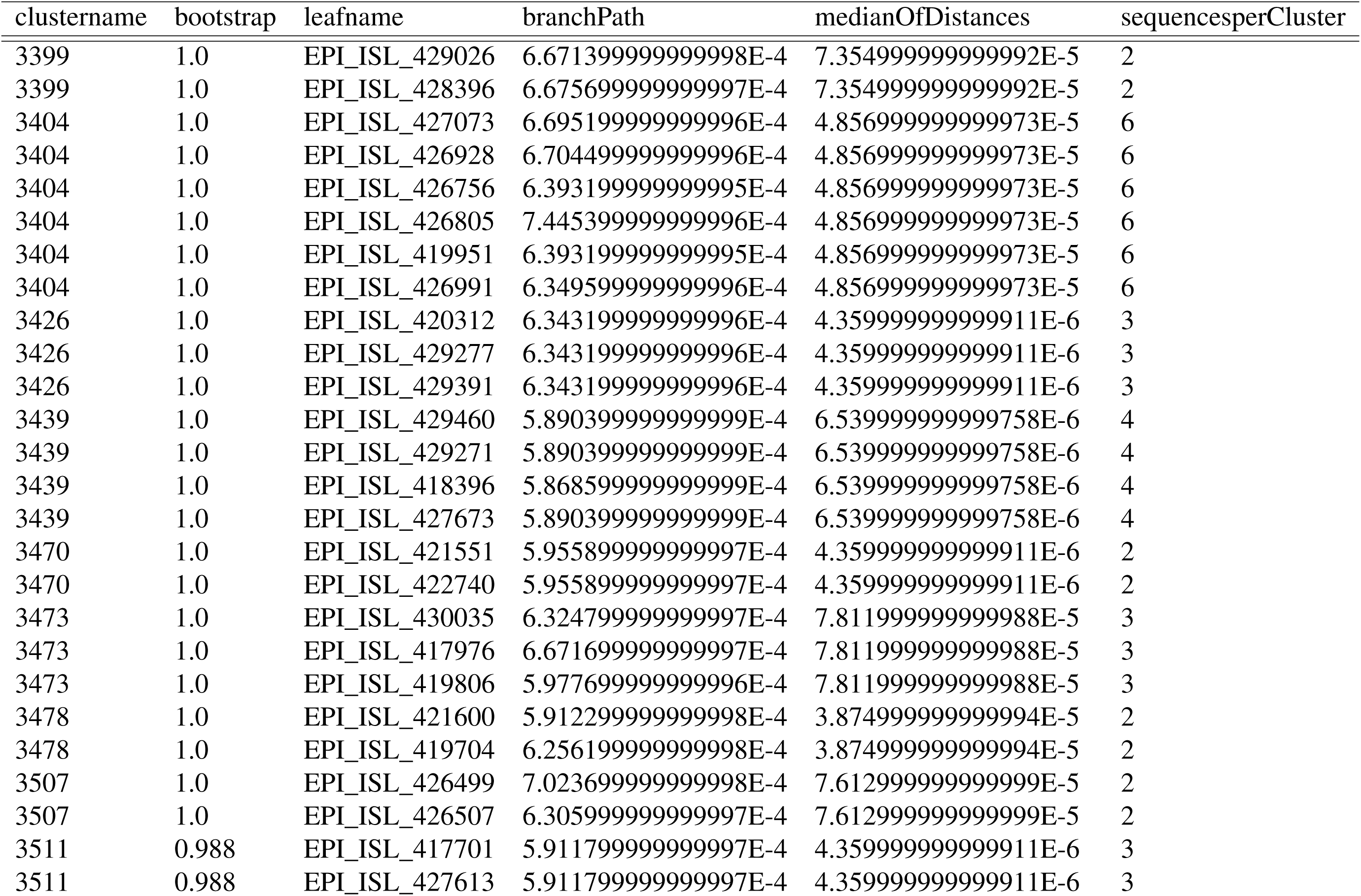
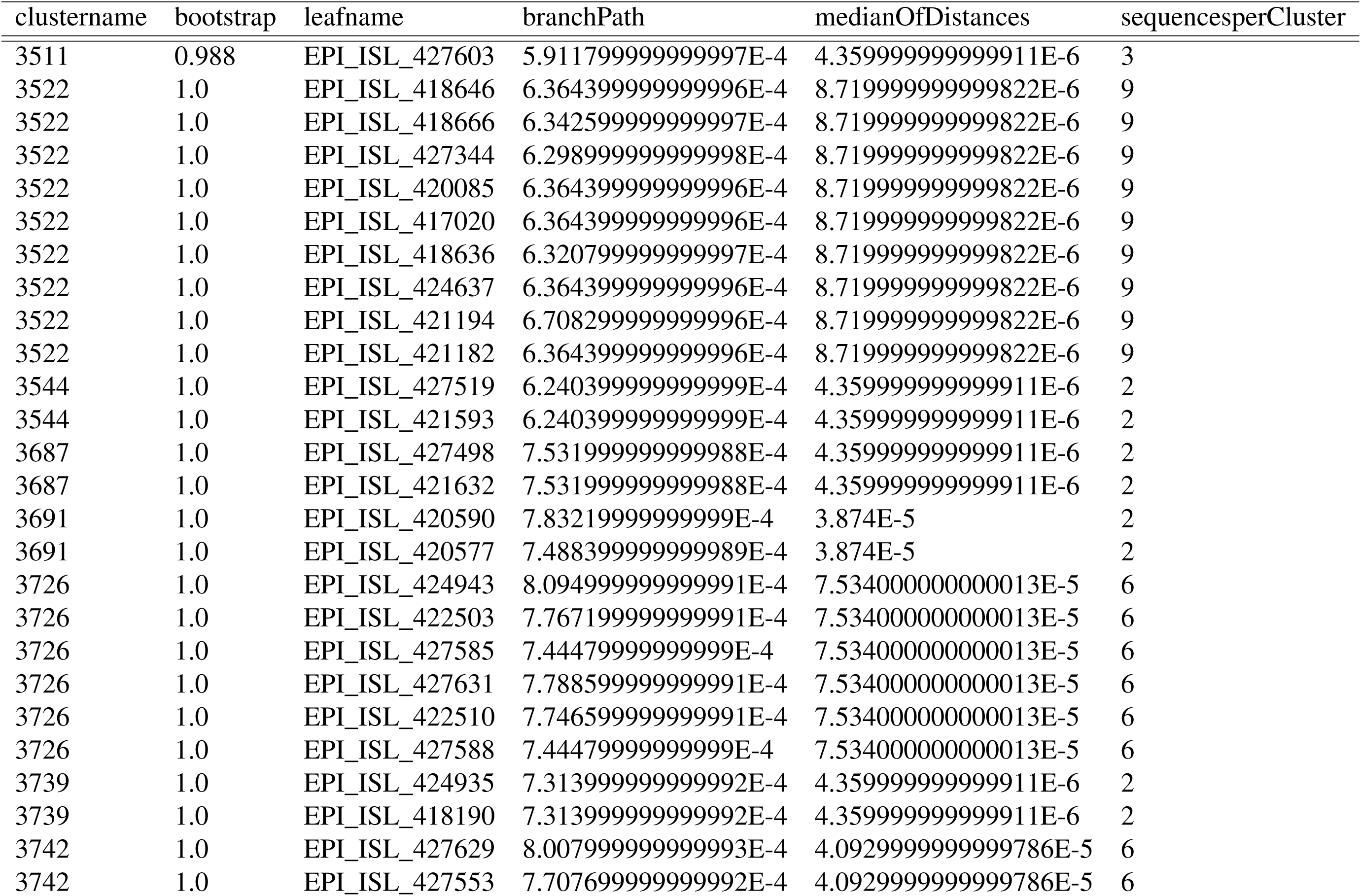
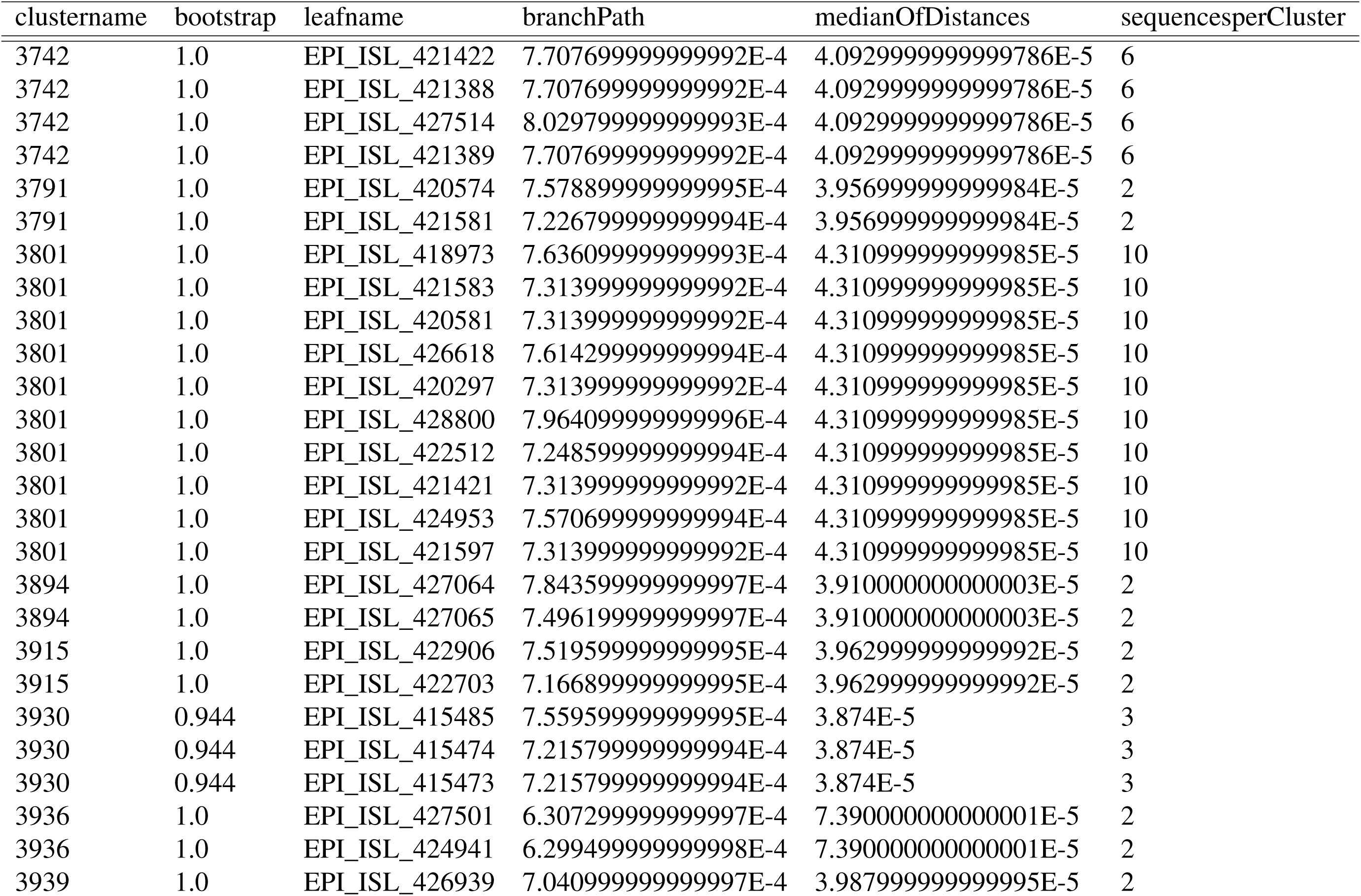
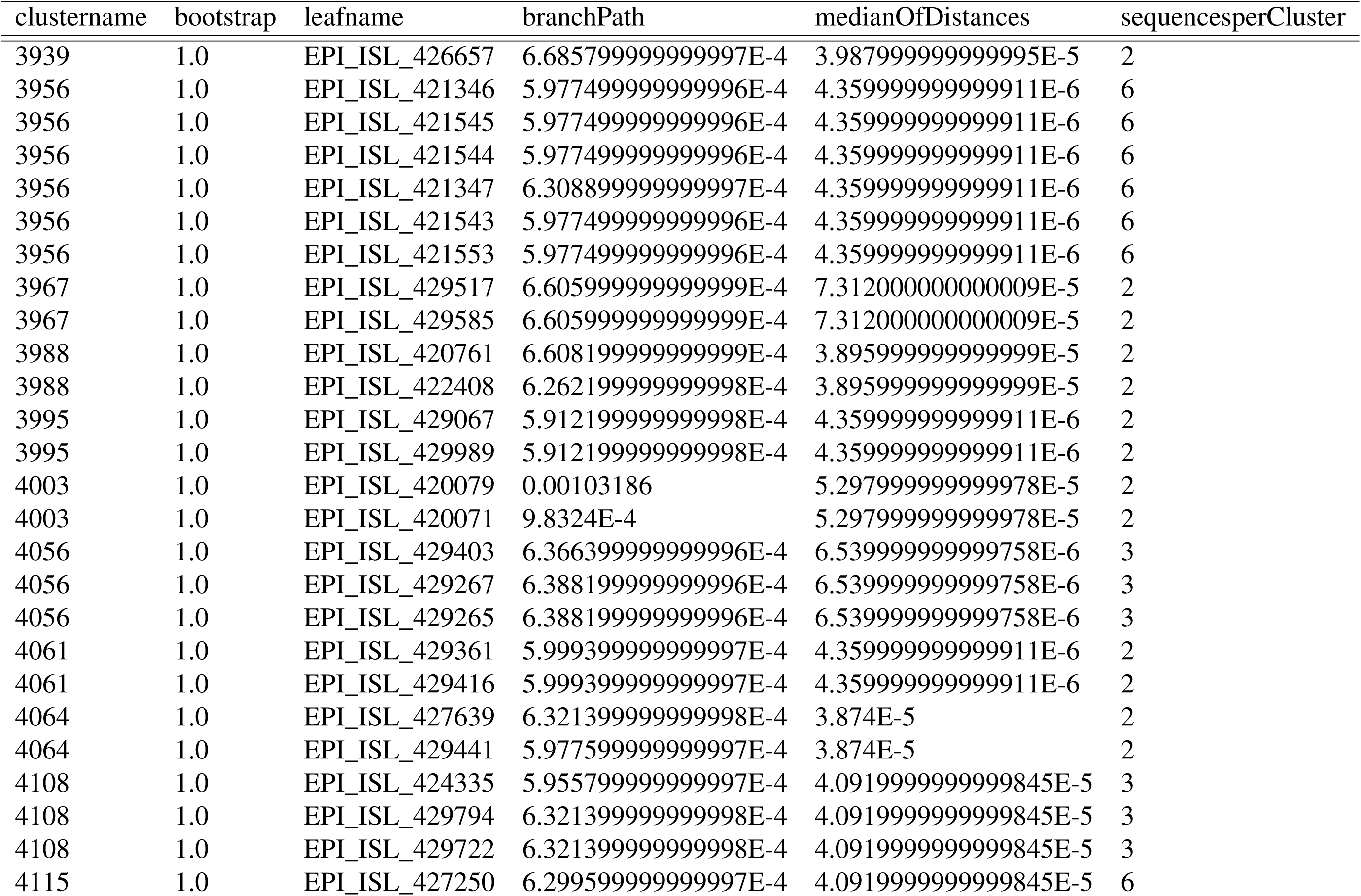
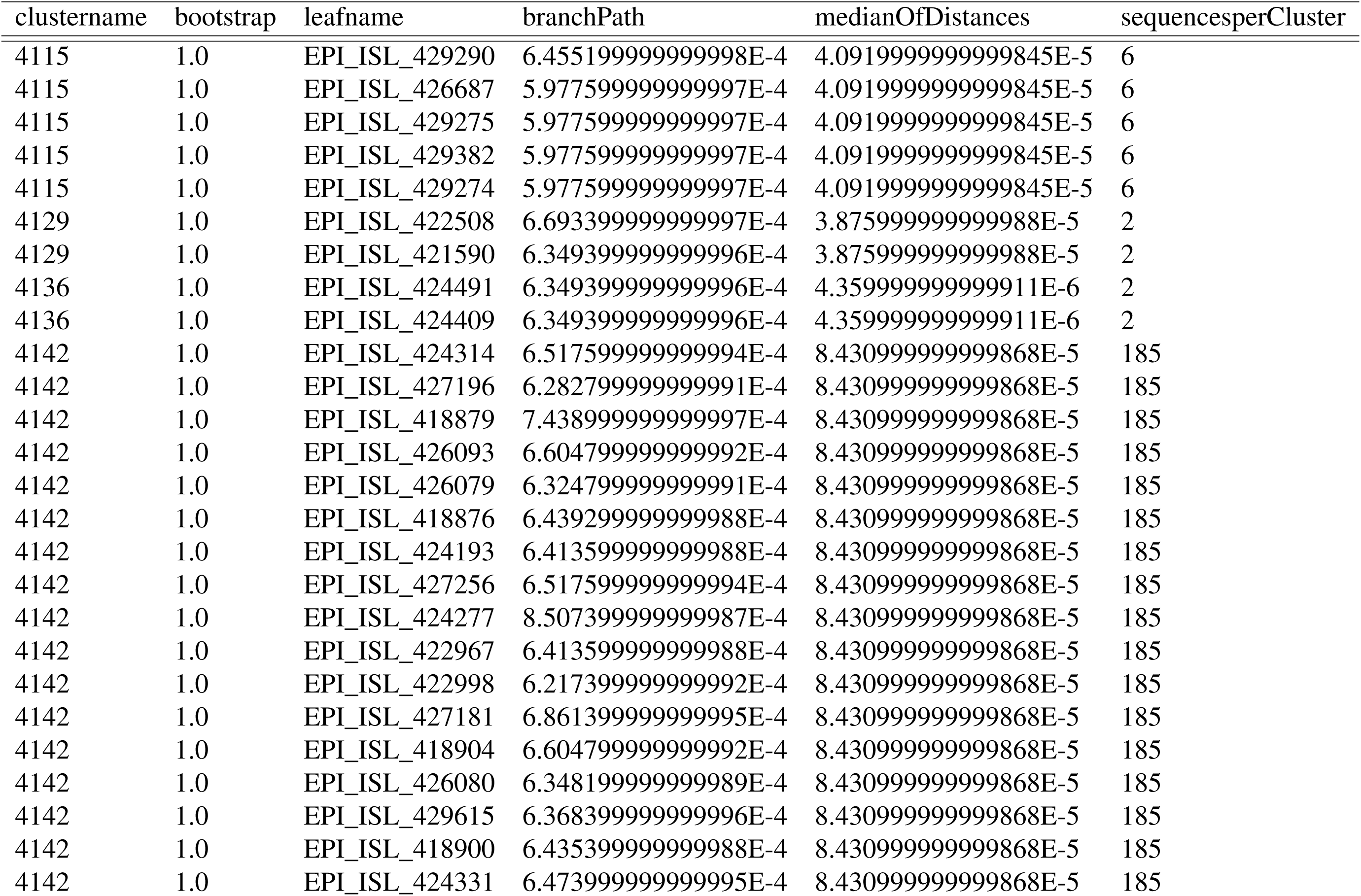
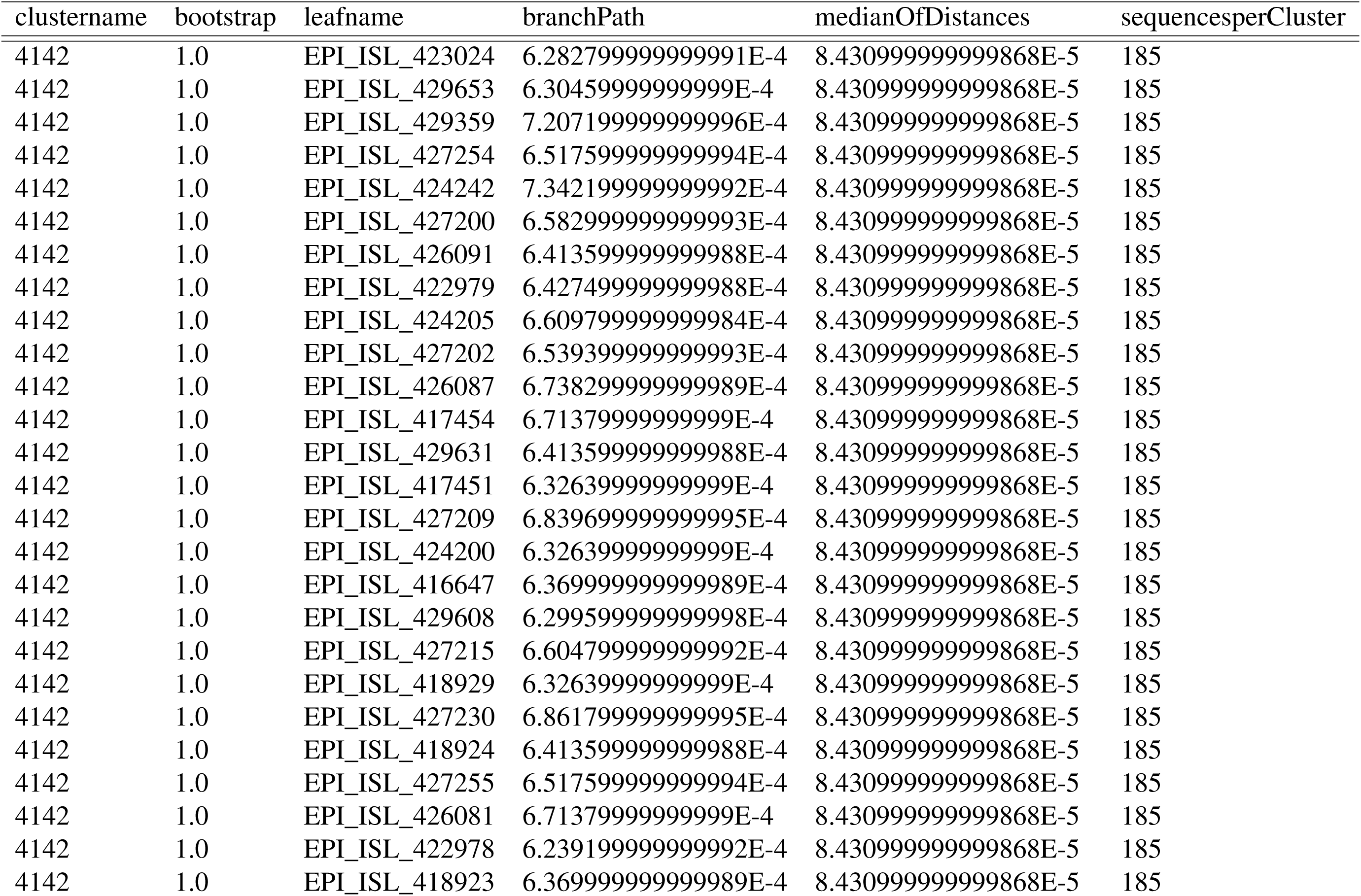
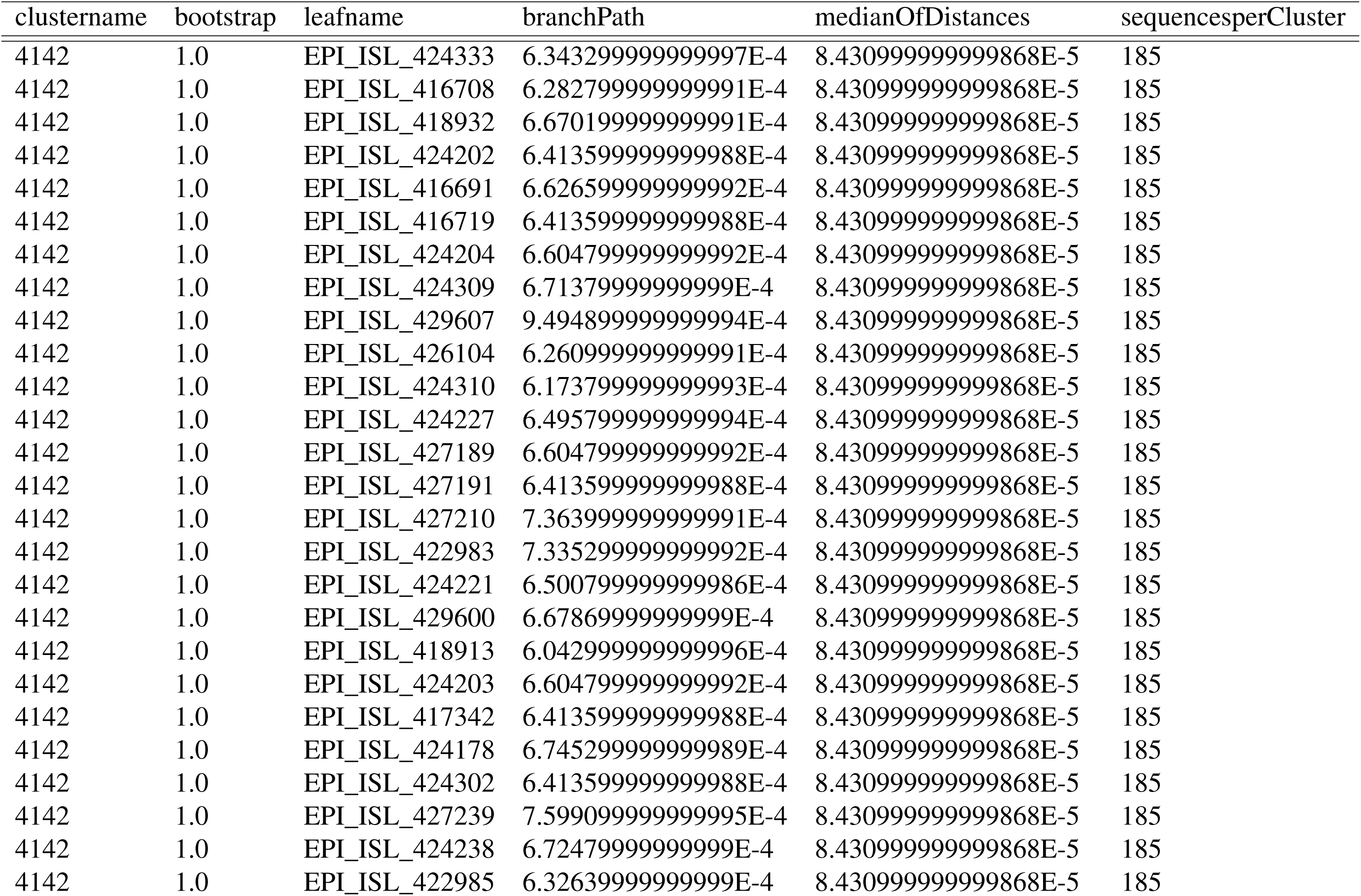
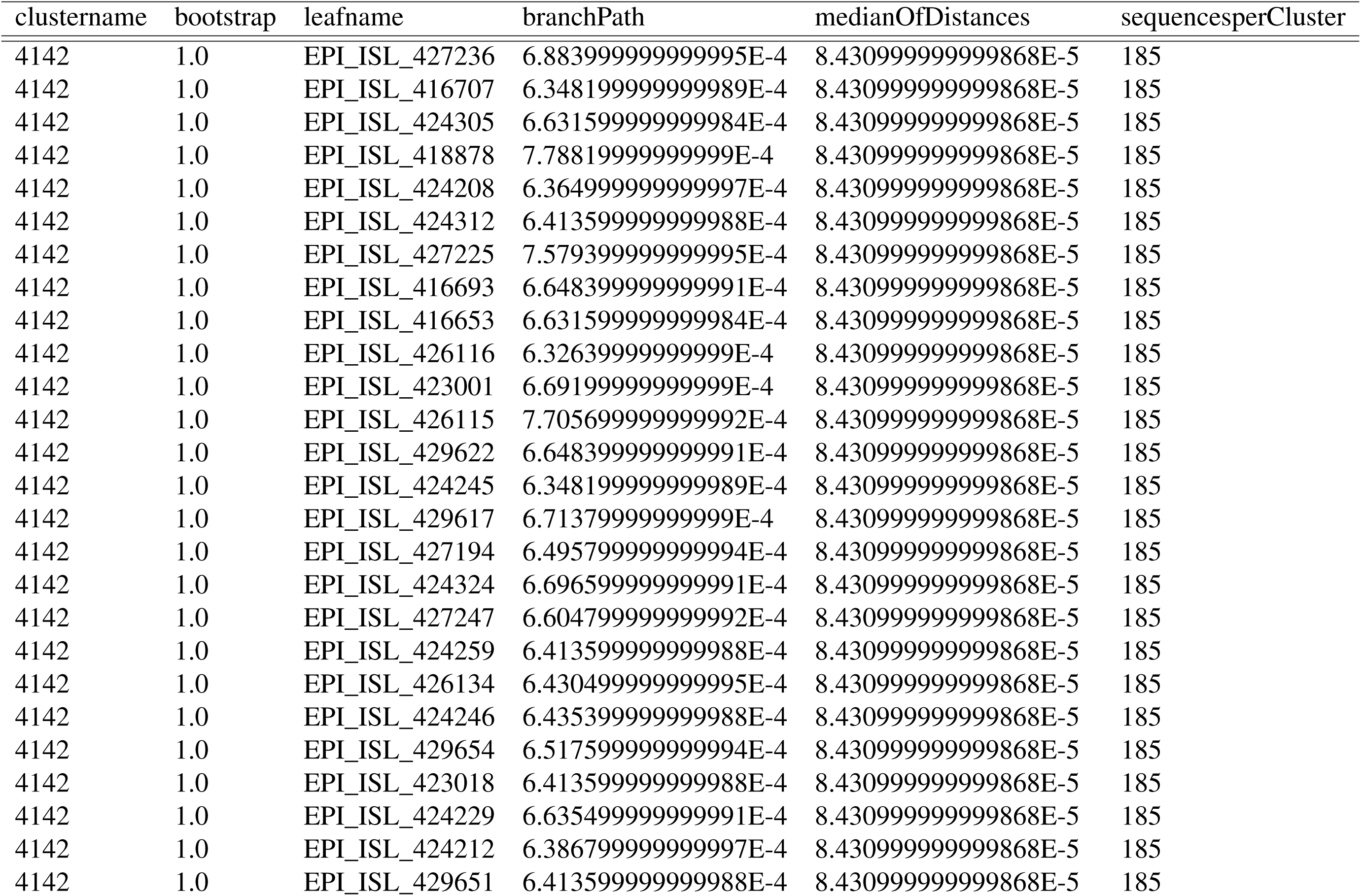
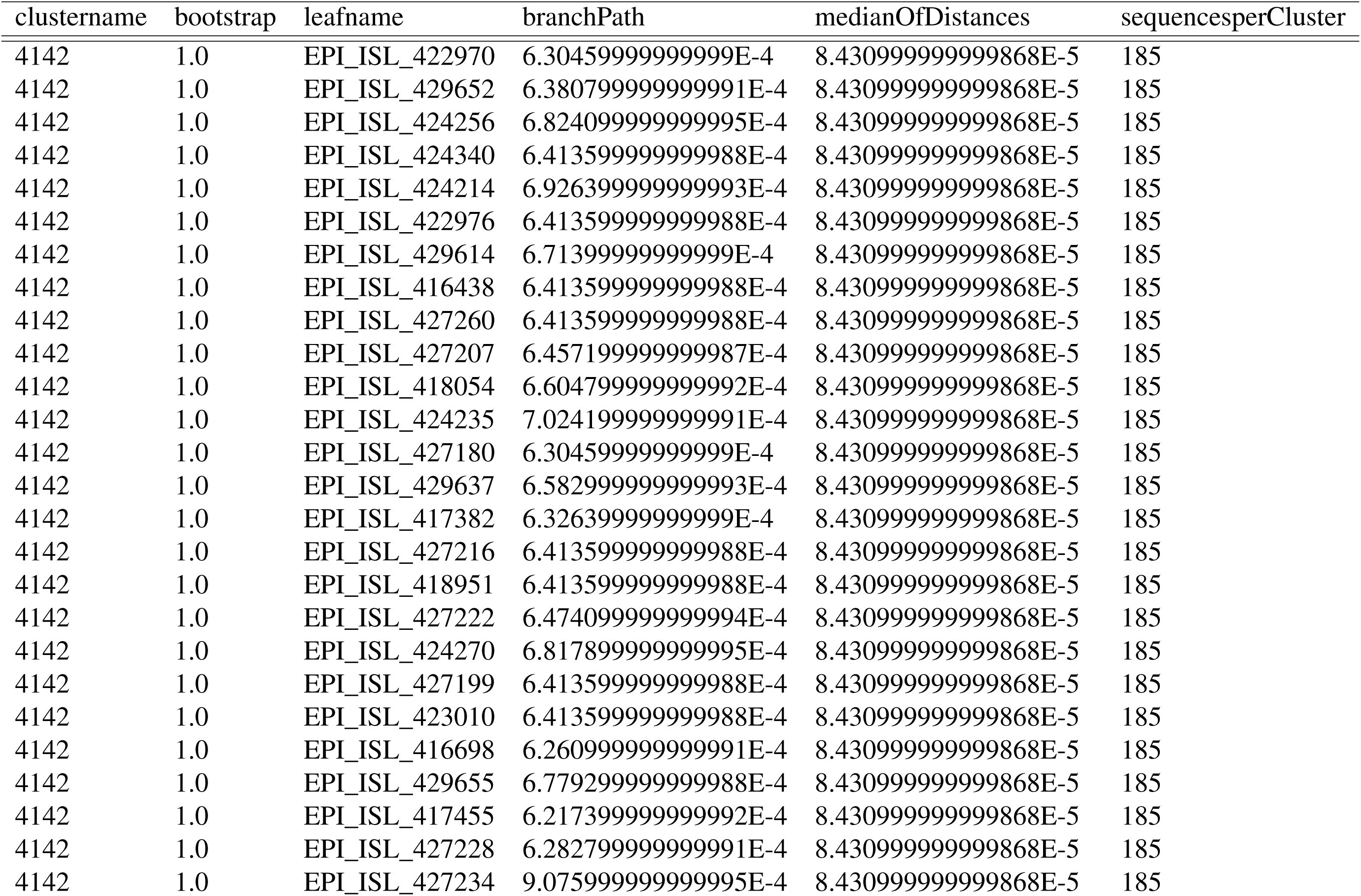
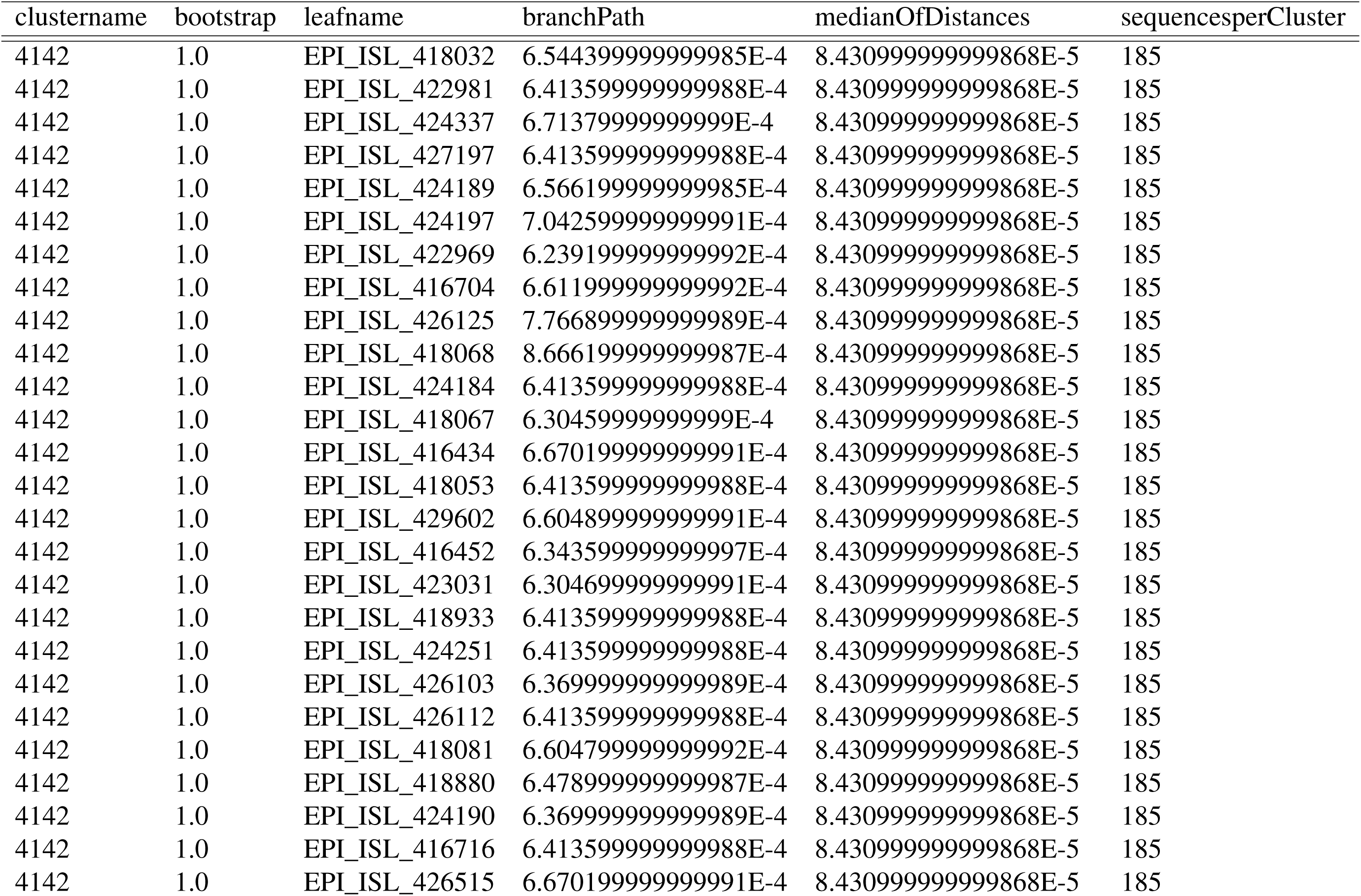
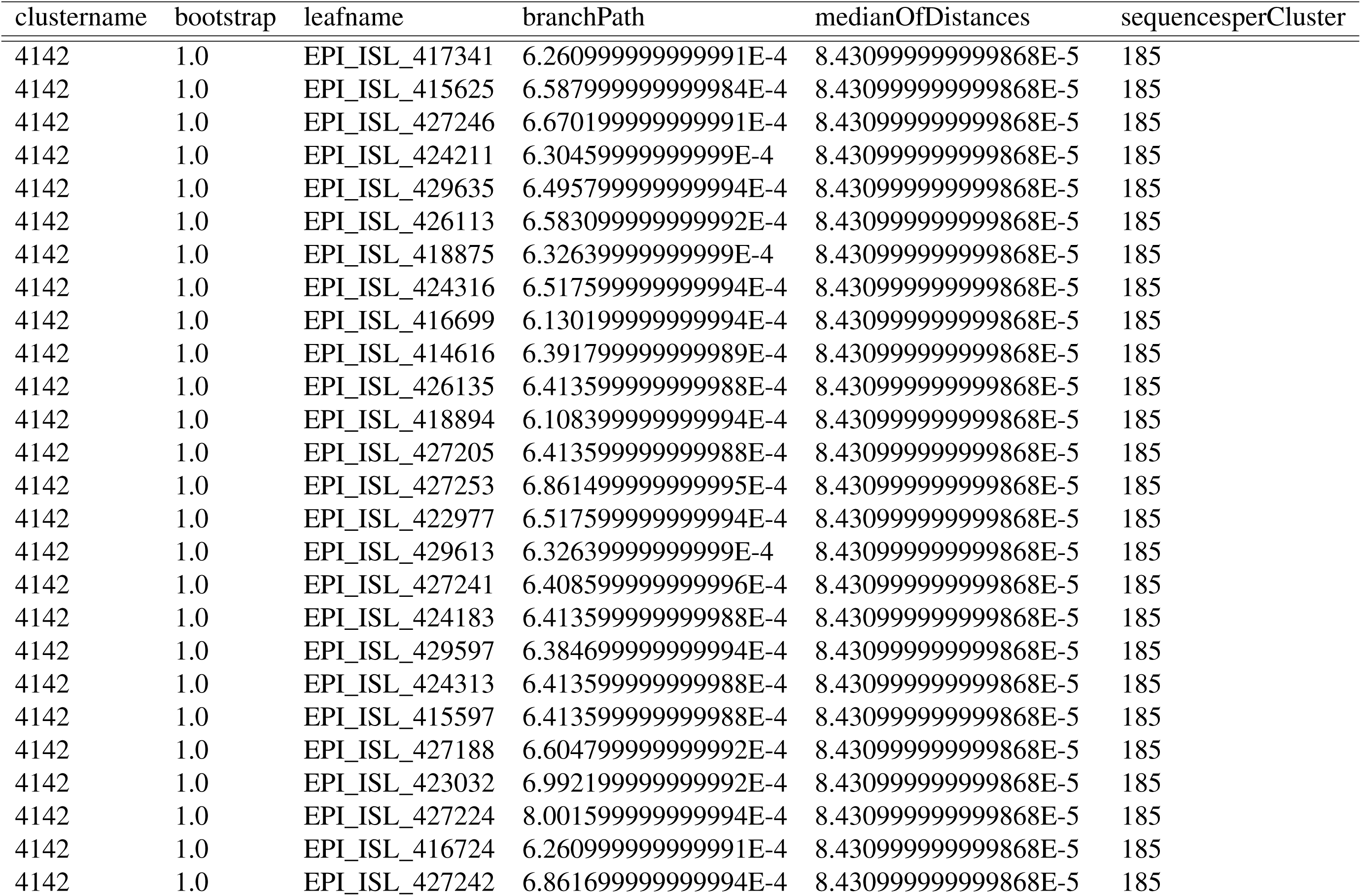
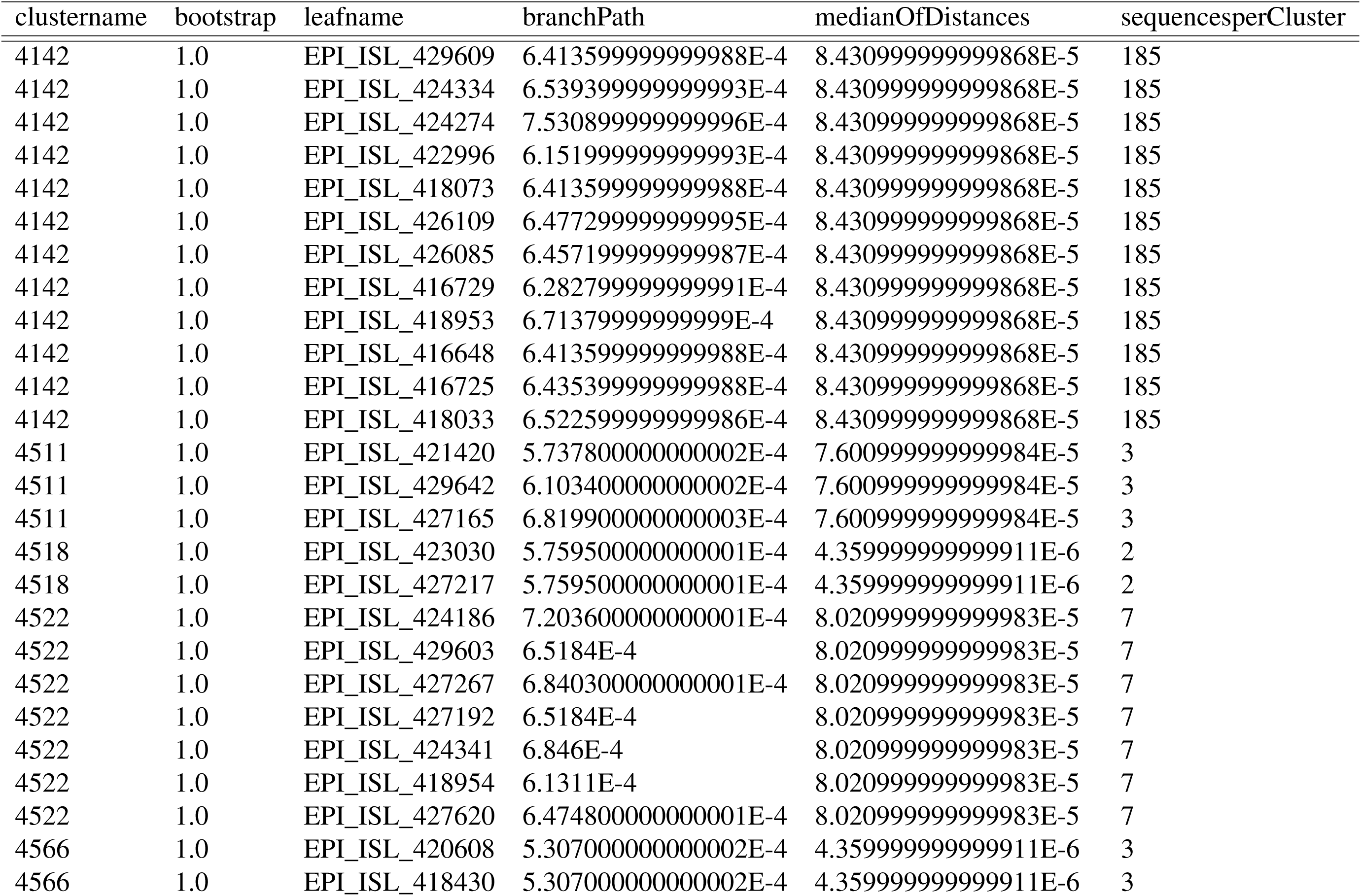
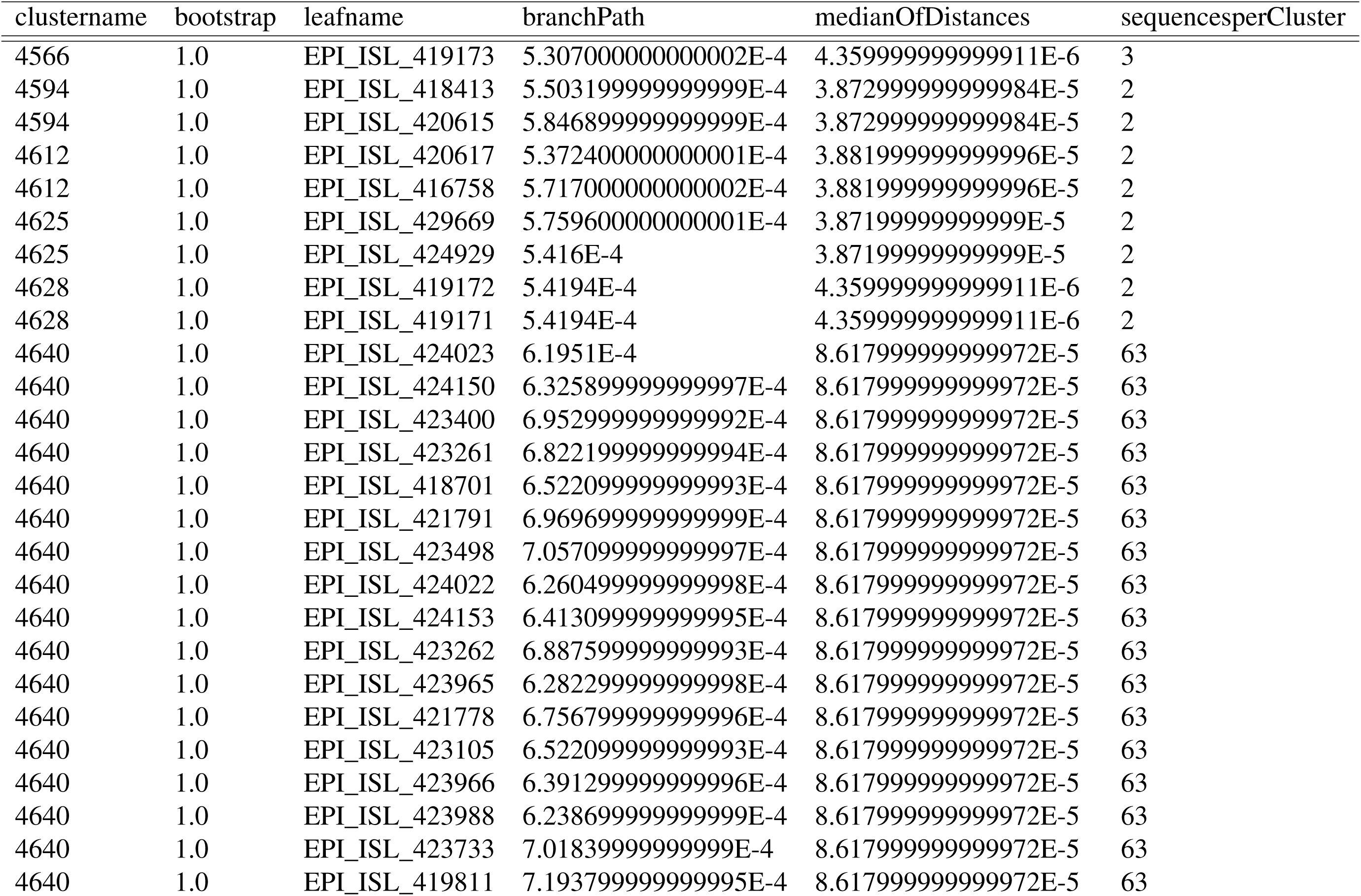
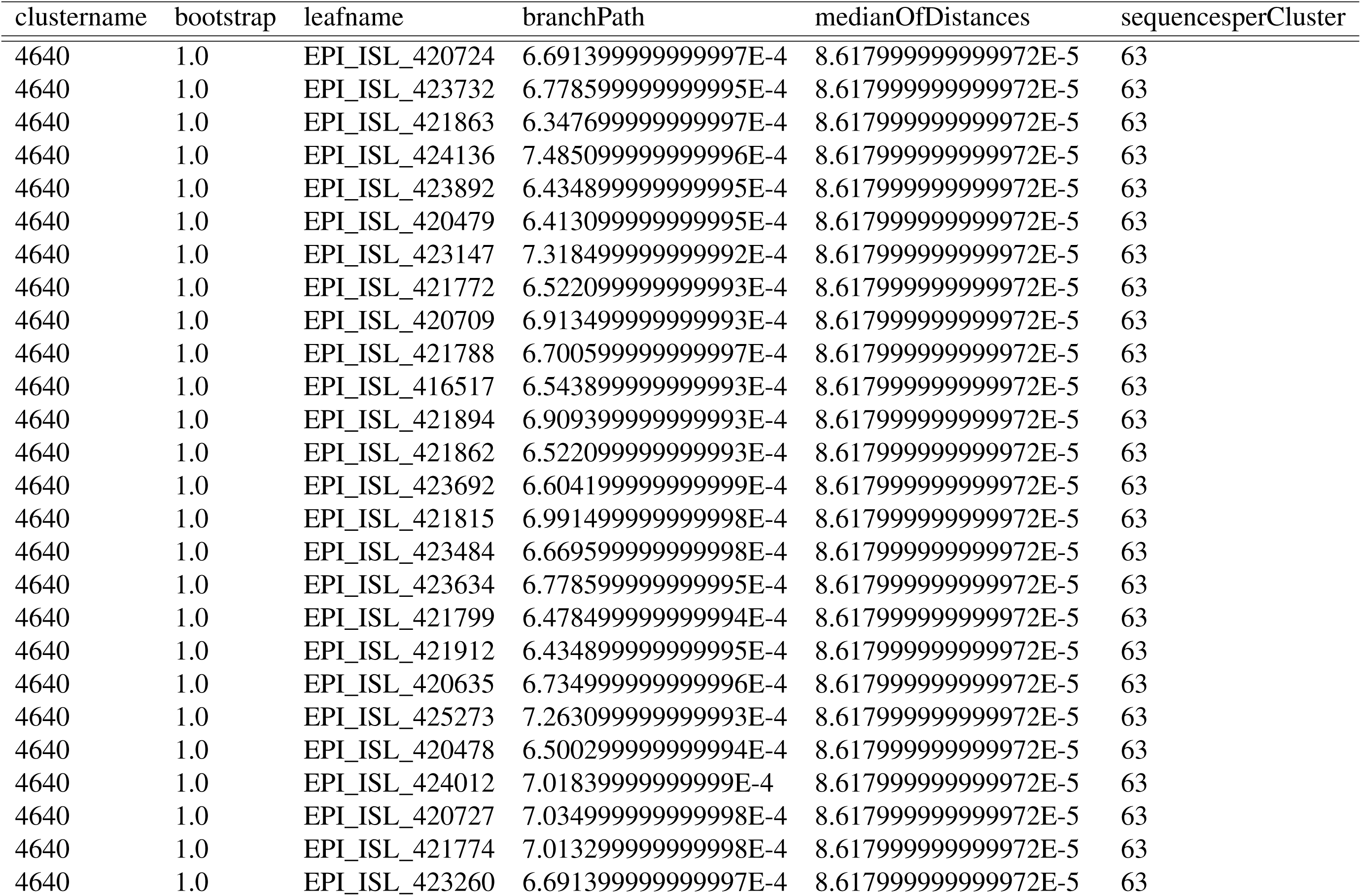
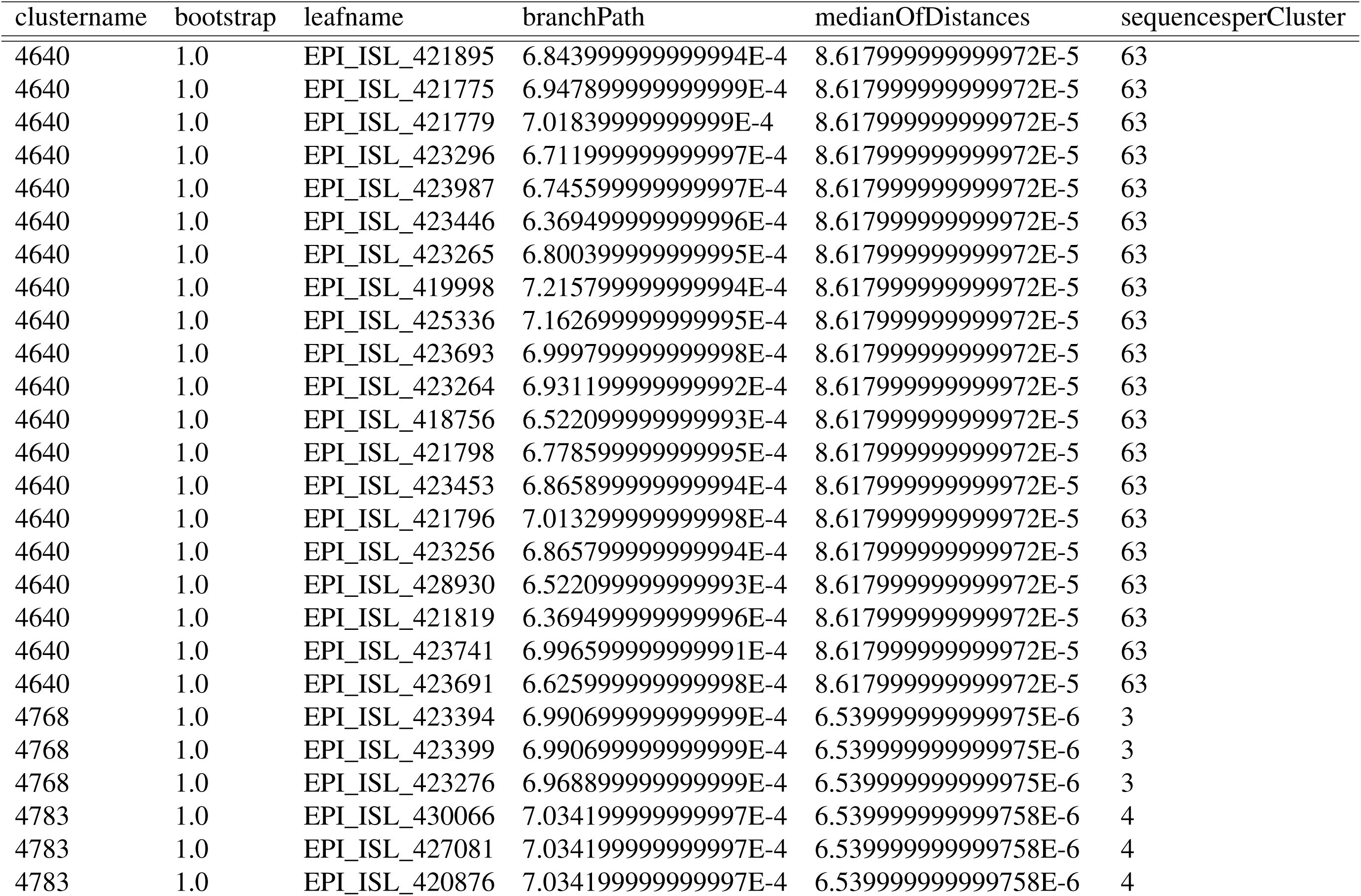
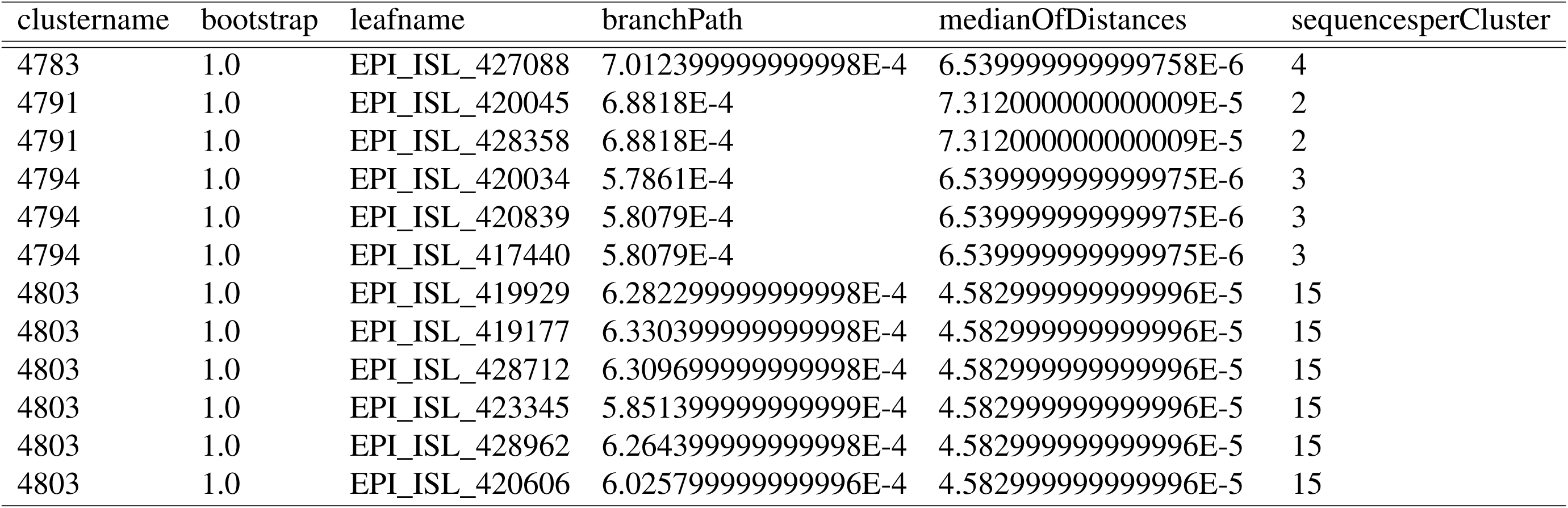
Clusters identified using a genetic distance threshold within 1% of the distribution of patristic distances within the entire tree. The minimum percentile threshold that maximized the number of clusters was chosen as the optimal threshold by performing multiple clustering runs on randomly sampled patristic distance distributions (1 million for each run) in Phylopart v2 (*46*).

